# The structural basis for LRRK2’s activation and autoinhibition

**DOI:** 10.64898/2026.05.31.729085

**Authors:** Amalia Villagran Suarez, Kathryn S Hatch, Tatyana Bodrug, Wei Gai, Katherine J Surridge, Elizabeth Moussikhina, Kendrick HV Nguyen, Marta Sanz-Murillo, Robert Callahan, Erica Xiong, Delisa Ramos, Lawrence Zhu, Verena Dederer, Sebastian Mathea, Janet Iwasa, Stefan Knapp, Kevan M Shokat, Samara L Reck-Peterson, Andres E Leschziner

## Abstract

Mutations in Leucine-Rich Repeat Kinase 2 (LRRK2) are the second most common cause of autosomal-dominant Parkinson’s disease (PD), and increased LRRK2 kinase activity is also observed in idiopathic PD, making LRRK2 a major actionable therapeutic target. LRRK2 is a 286-kDa multidomain enzyme containing a Ras-like GTPase (ROC) and a kinase domain. Using cryo-EM, biochemical reconstitution, and cell-based assays, we show that the ROC GTPase governs switching between autoinhibited and active states: GTP binding promotes activation, whereas GDP binding enforces autoinhibition. Two common PD-linked mutations, G2019S and R1441C/G/H, activate LRRK2 through distinct structural mechanisms, revealing genotype-specific routes to dysregulation. These findings provide a unified framework for understanding LRRK2 regulation with broad therapeutic implications. Stabilizing the GDP-bound state may inhibit LRRK2 by maintaining autoinhibition, whereas promoting the GTP-bound state could be advantageous in specific cellular contexts, such as the lung, where increased LRRK2 kinase activity may play protective or regulatory roles.

## INTRODUCTION

Mutations in Leucine Rich Repeat Kinase 2 (LRRK2) are one of the main causes of familial Parkinson’s Disease (PD)^1^. LRRK2 phosphorylates a subset of Rab GTPases associated with membrane vesicles, including those in the endolysosomal system and secretory pathway^2–4^. PD-linked mutations increase its kinase activity^5–7^, and increased Rab phosphorylation has been reported in postmortem samples from idiopathic PD patients^8^. This has made LRRK2 one of the main actionable targets for PD therapeutics, with several clinical trials targeting LRRK2 currently under way (clinicaltrials.gov).

LRRK2 is a large (286kDa), multi-domain protein, with an N-terminal half comprised of Armadillo (ARM), Ankyrin (ANK), and Leucine Rich Repeats (LRR) domains, and a C-terminal half containing both a Ras-family GTPase (Ras Of Complex, or ROC), and a kinase (Figure 1A). Structural studies revealed that LRRK2 is regulated by autoinhibition: the LRR, which connect to the ROC GTPase domain, cover the kinase’s active site, blocking access by Rab substrates (Figure 1A, 1B)^9^, and preventing the kinase from reaching its active state (Figure 1C)^9^. LRRK2 can form dimers and structural work suggests that the highest affinity dimerization interface is the one mediated by interactions between its C-terminal Of ROC (COR) domains (COR-A and COR-B)^9,10^ (Figure 1D).

**Figure 1.**
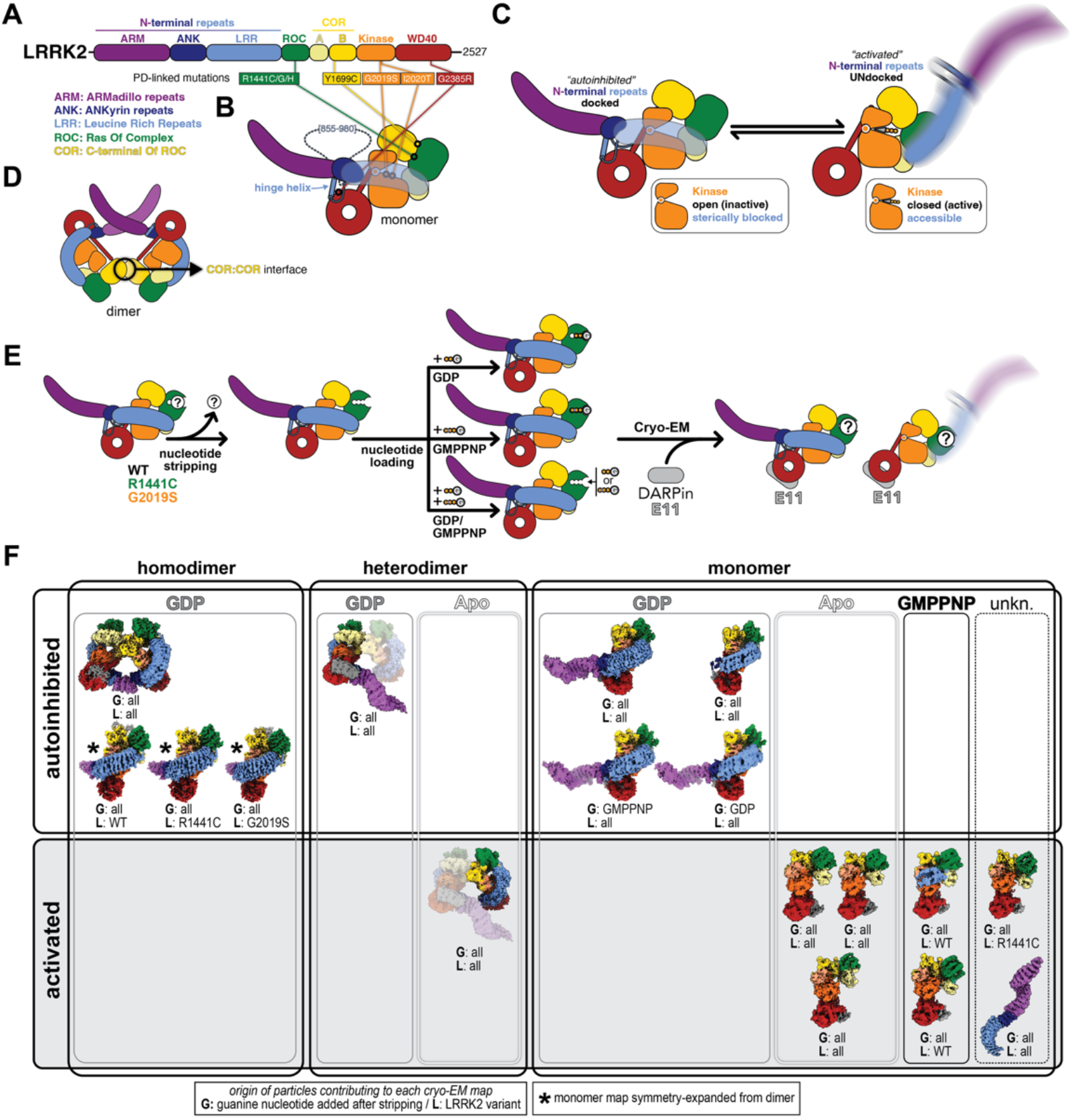
Cryo-EM reveals that GDP loss from the GTPase domain leads to LRRK2 activation. **(A**,**B)** Cartoon representations of the domain architecture (A) and structure of the LRRK2 monomer (B). These cartoons represent the autoinhibited conformation of LRRK2. Common PD-linked mutations are indicated in (A) and (B). We show the LRR as semi-transparent to make the active site of the kinase visible and use the same depiction in (C). **(C)** Cartoon representation of the autoinhibited and activated forms of LRRK2; this nomenclature is used throughout this work. Note: Although the cartoon for activated LRRK2 here shows the kinase in its closed or active conformation, we use ‘activated’ in the text to indicate only the conformation of the N-terminal repeats (i.e. undocked) regardless of the conformation of the kinase. **(D)** Cartoon representation of the LRRK2 dimer, mediated by COR:COR interactions. **(E)** Schematic representation of the strategy we used for our cryo-EM analysis: LRRK2 [WT, R1441C, or G2019S] was stripped of its nucleotide and then incubated with GDP, GMPPNP, or a mixture of both. Before preparing cryo-EM grids, samples were incubated with DARPin E11, which binds to the WD40 domain and reduces the preferred orientation of LRRK2 on cryo-EM grids^23^. Note: E11 is omitted from all cartoons of LRRK2 going forward for simplicity, although it is present in the cryo-EM maps. Data was collected from individual samples, but all data sets were merged and processed together initially, with sorting done after initial structures had been obtained and refined (see Methods and Figures S1-S7 for the data processing pipeline). **(F)** Table of all cryo-EM structures of LRRK2 obtained in this study for which models were built, as well as a map for the N-terminal repeats, for which no model was built. Structures are sorted according to whether they represent an autoinhibited or activated form of LRRK2, by the nucleotide state we determined for their GTPase domains (Apo, GDP, GMPPNP, or unknown), and whether they are dimeric or monomeric. For the heterodimers, the highlighted monomer is the one to which the descriptors apply. We indicate, below each map, what particles contributed to it based on the guanine nucleotide added to the sample (‘G’) and the LRRK2 variant (‘L’) used.

Despite the importance of autoinhibition in LRRK2’s function, its mechanism and regulation are not understood. Several studies have examined the role of the ROC GTPase domain in regulating LRRK2 activity, but the results are not consistent. Earlier work showed that addition of GTP or non-hydrolyzable GTP analogs to cell lysates correlates with increased LRRK2 kinase activity^11,12^, and that pathogenic mutations in the ROC–COR region can enhance GTP binding^11,13^ and affect GTP hydrolysis^14,15^. However, while nucleotide-dependent effects were frequently observed in lysate-based assays, they were reduced or absent when nucleotides were added to immunoprecipitated or purified LRRK2, indicating a strong dependence on experimental and cellular context^11,12^. Structurally, the direct connection between the LRR, which block access to the kinase active site, and the ROC domain makes its nucleotide state a prime candidate to regulate LRRK2. In support of this role, the interface between ROC and the adjacent COR-B domain is a hotspot for PD-linked mutations (Figure 1A, 1B), and some of these mutations have been shown to alter GTP binding^11,13^ or hydrolysis^14,15^. However, no structural studies have systematically examined the effect of the ROC nucleotide state on LRRK2 activation and autoinhibition. All structures of LRRK2 reported to date where the nucleotide state of the ROC domain could be determined^9,10,16–19^, including those capturing an active conformation^18^, were bound to GDP.

Here, we set out to determine how the nucleotide state of the ROC GTPase regulates LRRK2 activation and autoinhibition, structurally and mechanistically. Understanding this regulation could open new therapeutic avenues to allosterically inhibit LRRK2’s kinase activity. The accumulated evidence has shown that different types of kinase inhibitors stabilize different LRRK2 conformations^10,16,17^, and that these conformations have different cellular localizations and/or interactomes^6,10,20–22^. Although the clinical relevance of these differences is not yet clear, they suggest that developing multiple approaches for targeting LRRK2 is desirable. In parallel, there is growing evidence that LRRK2 plays an important role in the lung and that loss of function is linked to both pulmonary fibrosis^23,24^ and tumorigenesis^25^. Thus, allosteric activators of LRRK2 may be useful therapeutics as well.

The foundation of this study is an unbiased structural approach to understand if and how the nucleotide state of LRRK2’s ROC domain correlates with LRRK2’s active and autoinhibited conformations. Throughout this work, we use ‘activated’ to refer to any LRRK2 conformation where its N-terminal repeats are undocked, and ‘autoinhibited’ to refer to those conformations where the N-terminal repeats are docked and thus blocking access to the kinase active site. Note that, in this context, activated refers to accessibility to the kinase active site and not to kinase activity itself.

## RESULTS

### Cryo-EM reveals that GDP loss from the GTPase domain leads to LRRK2 activation

To determine structures of LRRK2 in different nucleotide states, we used a nucleotide exchange protocol to strip off the GDP that is typically bound to purified LRRK2 (see Methods). We then incubated LRRK2 with one of the following: (1) GDP; (2) the non-hydrolyzable GTP analog GMPPNP; or (3) a mixture of GDP and GMPPNP (Figure 1E). We used three different LRRK2 variants: wild-type and the PD-linked mutants G2019S and R1441C (Figure 1A, 1B). G2019S, found in the kinase’s active site, is the most common PD-linked mutation in LRRK2. R1441C is found in the ROC domain at the interface between ROC and COR-B (Figure 1A, 1B). We collected data from the resulting samples and combined all the data sets into one for the initial data processing. Later, we split some of the data according to the LRRK2 variant or the nucleotide used in the samples (see Methods and Figures S1-S7). All samples were incubated with the Designed Ankyrin Repeat Protein (DARPin) E11, which we previously showed binds to the WD40 domain and does not affect kinase activity in vitro^26^. The reason for including DARPin E11 in our samples is that its presence increases the range of orientations LRRK2 adopts on cryo-EM grids, which leads to higher resolution structures^26^.

Data processing revealed a significant level of compositional and conformational heterogeneity in our samples, as expected, including multiple cryo-EM maps representing activated LRRK2 (Figure 1F, and Figure S1). We were able to build models for 16 distinct cryo-EM maps (Figure 1F, and Figures S1-S7). Nine of these maps corresponded to autoinhibited LRRK2 and could be classified into three states (Figure 1F): a LRRK2 monomer with the N-terminal repeats docked; the dimeric form of this species; and a mixed dimer where one LRRK2 protomer had its repeats docked (autoinhibited) while the other had them undocked (activated). The eight maps representing activated LRRK2 could be classified into three structures (Figure 1F): the same mixed dimer (which is counted twice as it contains both autoinhibited and activated LRRK2); structures showing only the C-terminal half of LRRK2 containing the ROC, COR, Kinase, and WD40 domains; and structures showing only the N-terminal repeats.

### Autoinhibited LRRK2 is bound to GDP

Next, we determined the nucleotide state of ROC for the highest resolution map representing the monomeric form of autoinhibited LRRK2 (Figure 2A, and Figures S2, S3). In agreement with previous structures^9,16,17^, this map fit GDP better than GMPPNP (Figure 2B, 2C). We then asked which data sets had contributed to this structure. Surprisingly, we found that particles coming from samples treated with any of the three nucleotide conditions contributed to the map, with a majority coming from samples treated with GMPPNP. We split the data by nucleotide added, and generated new maps comprised of particles from samples that had been treated either with GDP (Figure 2D) or GMPPNP (Figure 2E). This revealed that the map arising from samples treated with GMPPNP also accommodated GDP better than GMPPNP (Figure 2F, 2G). We interpreted this as incomplete nucleotide stripping, with some LRRK2 molecules maintaining their originally bound GDP throughout the process.

**Figure 2.**
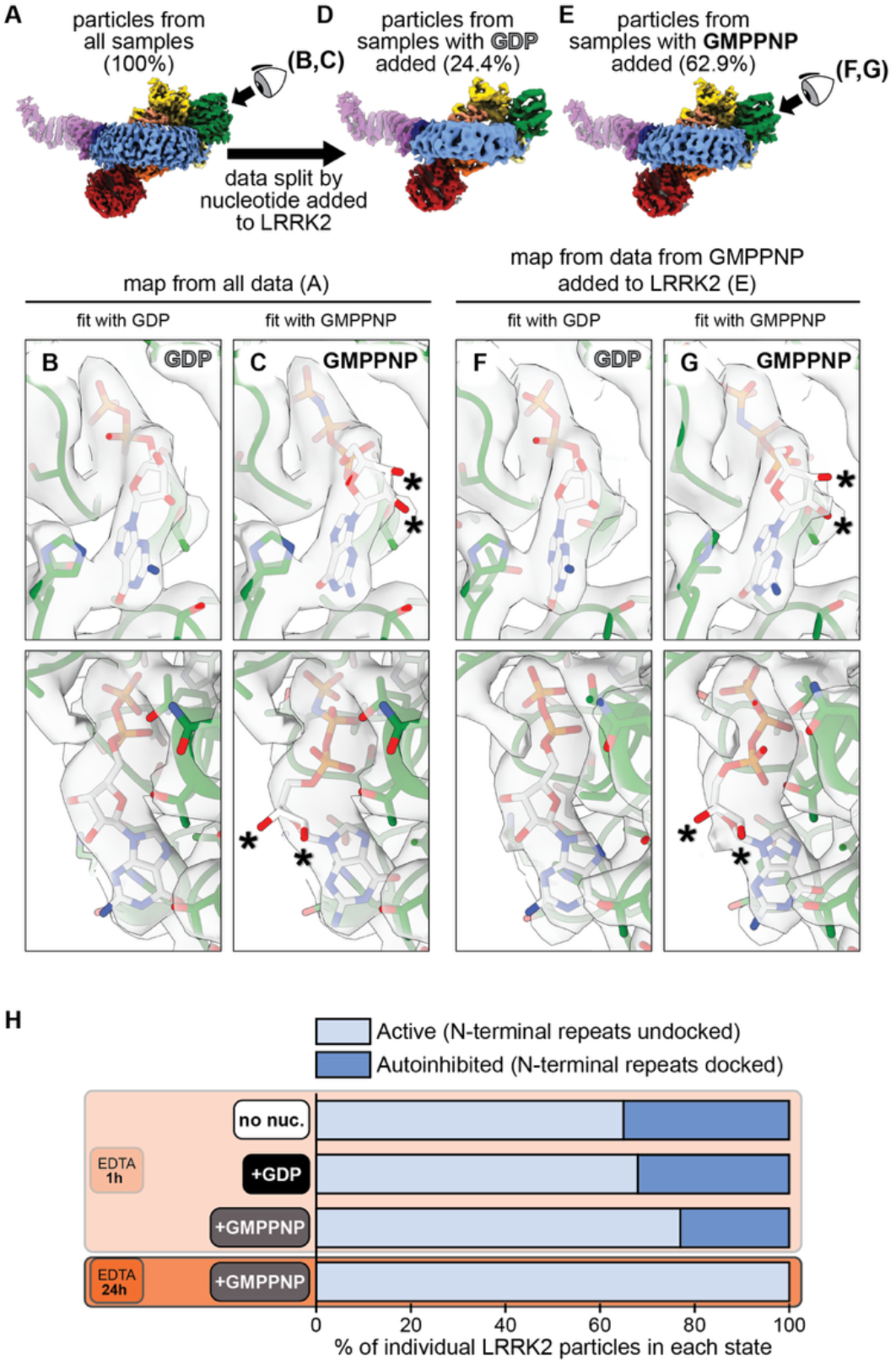
Autoinhibited LRRK2 is bound to GDP. **(A)** Cryo-EM map of autoinhibited LRRK2 comprised of particles coming from all samples (i.e. not sorted based on the nature of the samples). **(B**,**C)** Close-ups of the nucleotide-binding pocket for the map in (A) with the model built and refined containing either GDP (B), or GMPPNP (C). The asterisks mark the 2’-OH and 3’-OH of the ribose in GMPPNP, as these are the features whose fit in the map differs most between the two nucleotides. Once we determined that the map in (A) was comprised of particles coming from samples treated with GDP (24.4%), GMPPNP (62.9%), and GDP + GMPPNP (12.7%), we reconstructed maps containing particles arising from samples treated only with GDP **(D)** or GMPPNP **(E). (F, G)** Close-ups of the nucleotide-binding pocket for the map in (E) with the model built and refined containing either GDP (F), or GMPPNP (G). As in (B,C), asterisks mark the 2’-OH and 3’-OH of the ribose in GMPPNP. (H) Distribution of LRRK2 particles (cryo-EM data) into active and autoinhibited conformations as a function of the length of the nucleotide stripping treatment (EDTA) and nucleotide added after it. (See Figure S8 for data processing pipeline.)

To test this, we prepared a sample where we extended the nucleotide stripping step from 1h to 24h; while samples stripped for 1h showed 23-35% of their particles in the autoinhibited conformation, the sample stripped for 24h had 100% of the particles in the active state (Figure 2H and Figure S8). Taken together, our data show that autoinhibited LRRK2 is in a GDP-bound state.

### Active LRRK2 is either bound to GMPPNP or lacks nucleotide

We next focused on the structures of activated LRRK2, where the N-terminal repeats are undocked and thus not seen in our cryo-EM maps because the repeats are flexible relative to the C-terminal half of LRRK2. (Figure 3A-3F, and Figures S4, S6). The activated structures of LRRK2 also differed from the GDP-bound, autoinhibited form of LRRK2 in the conformational heterogeneity of their ROC and COR-A/B domains, which show flexibility relative to the kinase and to each other (Figure 3G-3J and Figure S9).

**Figure 3.**
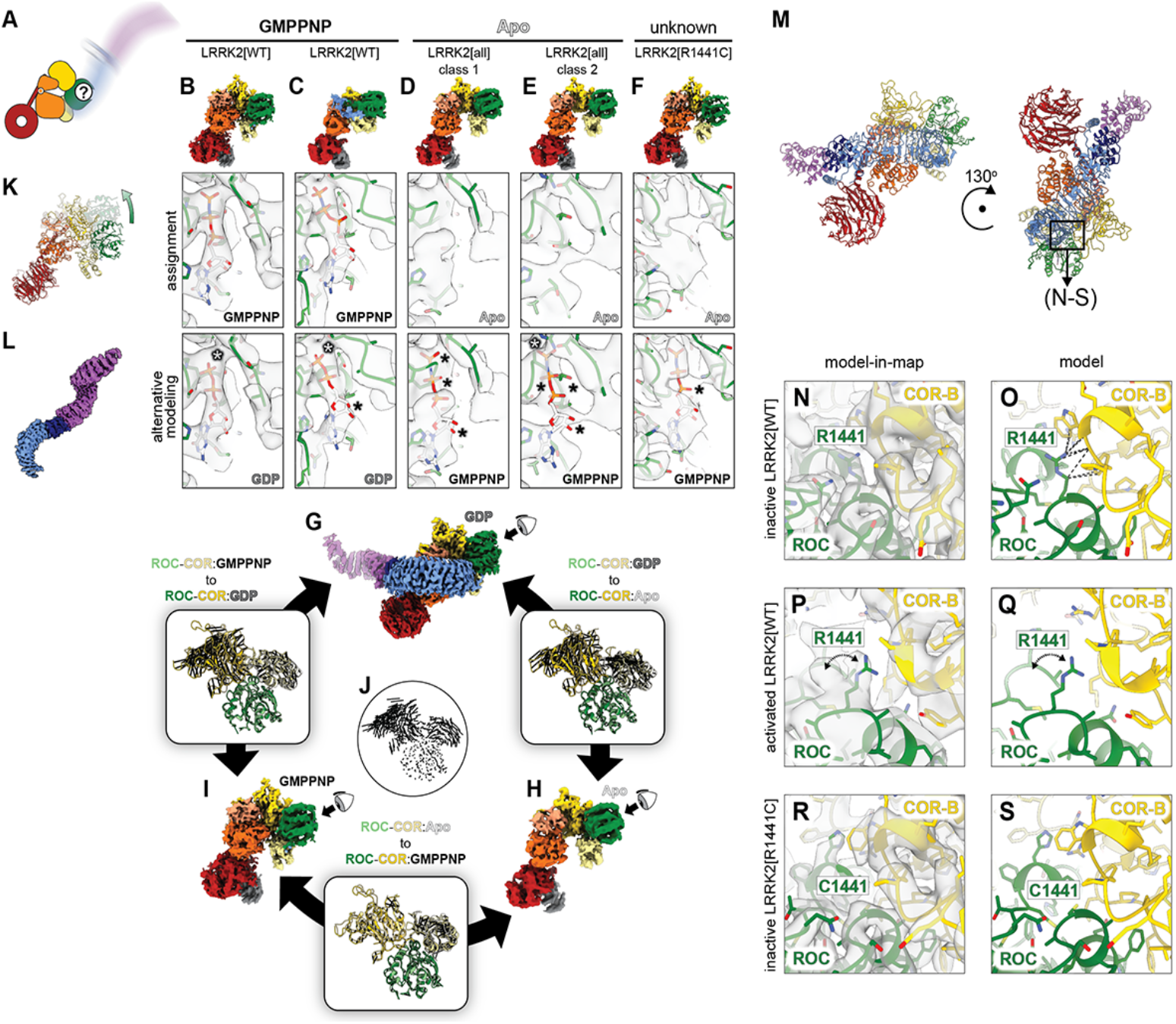
Activated LRRK2 is bound to GMPPNP or lacks nucleotide. **(A)** Schematic representation of activated LRRK2, with its repeats undocked. **(B-F)** Guanine nucleotide states of the ROC domains of activated LRRK2. Maps are shown for 5 different forms of activated LRRK2. Below the maps, we show close-ups of the nucleotide binding pocket of the ROC domain. The top row shows the nucleotide assignments we made based on the analysis of the structures. The bottom row shows an alternative modeling to highlight the reasons why it was rejected. The asterisks highlight problems with the fitting: black asterisks mark portions of the nucleotide that do not fit inside the density while white asterisks mark parts of the map that are not accounted for by the nucleotide or come too close to it. The models on the top row were refined in the presence of GMPPNP (B, C) or absence of nucleotide (D-F). The panels on the bottom row show the same maps and protein models as in the top row, with the alternative nucleotide modeled in by aligning the structure that contains it to the one shown in the panel and then displaying only the nucleotide. For panels labeled ‘GDP’, we aligned the ROC domain from LRRK2:G10:GDP (Figure 5E). For those labeled ‘GMPPNP’, we aligned the structure of LRRK2[WT]:GMPPNP (this figure, panel (B)). **(G-J)**, Conformational changes in ROC-COR between different nucleotide states of LRRK2. We aligned the ROC-COR modules of autoinhibited LRRK2:GDP (G), activated LRRK2:Apo (H), and activated LRRK2:GMPPNP (I) by their GTPase domains. The panels between the structures show the ROC-COR modules from the two structures along with vectors, in black, connecting equivalent alpha carbons between the structures (shown by themselves as an example in (J)). **(K)** Superposition of the model for activated LRRK2 with partial LRR density (C), and the C-terminal half from autoinhibited LRRK2 (LRRK2:G10:GDP, Figure 5E) to highlight that the former is in the activated conformation. The autoinhibited structure is shown in dim colors, and the green arrow shows the different positions of the ROC-COR module in the two models. **(L)** Cryo-EM map, colored by domain, of the undocked N-terminal repeats of LRRK2. The images contributing to this map originated from the same particles that gave rise to the maps of the C-terminal half of LRRK2 shown in this figure. **(M-S)** R1441 is involved in stabilizing the ROC:COR-B interface (the area within the box in (M)). **(N, O)** Model-in-map (N) and model alone (O) for close-ups of the ROC:COR-B interface in autoinhibited LRRK2 (the LRRK2:G10:GDP structure) showing interactions between R1441 and COR-B. **(P, Q)** The same interface in the activated form of LRRK2[WT]:GMPPNP. Because the conformational changes in ROC-COR in activated LRRK2 involve a movement of COR-B away from ROC, R1441 no longer interacts with COR-B. The cryo-EM map of activated LRRK2[WT]:GMPPNP (P) shows density for two conformations of R1441, neither of which interacts with COR-B. The dashed arrow indicates the two conformations. **(R, S)** Model-in-map and model alone for activated LRRK2[R1441C] showing the R1441C mutation. A comparison with panels (N, O) shows how the stabilizing interaction between ROC and COR-B in autoinhibited LRRK2 cannot happen in the R1441C mutant.

We obtained five volumes representing activated LRRK2 for which we were able to build models (Figure 3B-3F, and Figures S4-S6) and could determine the nucleotide state of the ROC domain for four of them; two volumes contained GMPPNP (Figure 3B, 3C) while two lacked nucleotide (Figure 3D, 3E). These structures suggest that LRRK2 activation requires the loss of GDP, which our data show correlates with the autoinhibited form, and that the activated state was maintained either in the absence of nucleotide or when GTP (GMPPNP here) was bound. One GMPPNP-bound structure of activated LRRK2 was unique because we observed density for the C-terminal portion of the LRR, which connects to the GTPase (Figure 3C), possibly representing an intermediate of activation or autoinhibition (Figure 3K, and Figure S9B, S9C).

We wanted to test the idea that removal of GDP was sufficient to activate LRRK2. We first tested this structurally: we applied our nucleotide-stripping protocol to LRRK2 without subsequently adding back a nucleotide. Data processing of images collected from this sample showed that a significant fraction of the LRRK2 molecules were in their active conformation (i.e. the N-terminal repeats are not seen) (Figure S10A-S10C). Next, we tested this functionally by measuring the effect of GDP removal on LRRK2 phosphorylation of Rab8a in vitro (Figure 4A). The 24h incubation in the ‘mock’ nucleotide stripping (i.e. without EDTA) leads to some loss of LRRK2 and an associated reduction in activity relative to an untreated sample (Figure 4B and Figure S10D). Incubation with EDTA, however, leads to increased phosphorylation of Rab8a relative to both untreated and ‘mock’ treated LRRK2 (Figure 4B and Figure S10D).

**Figure 4.**
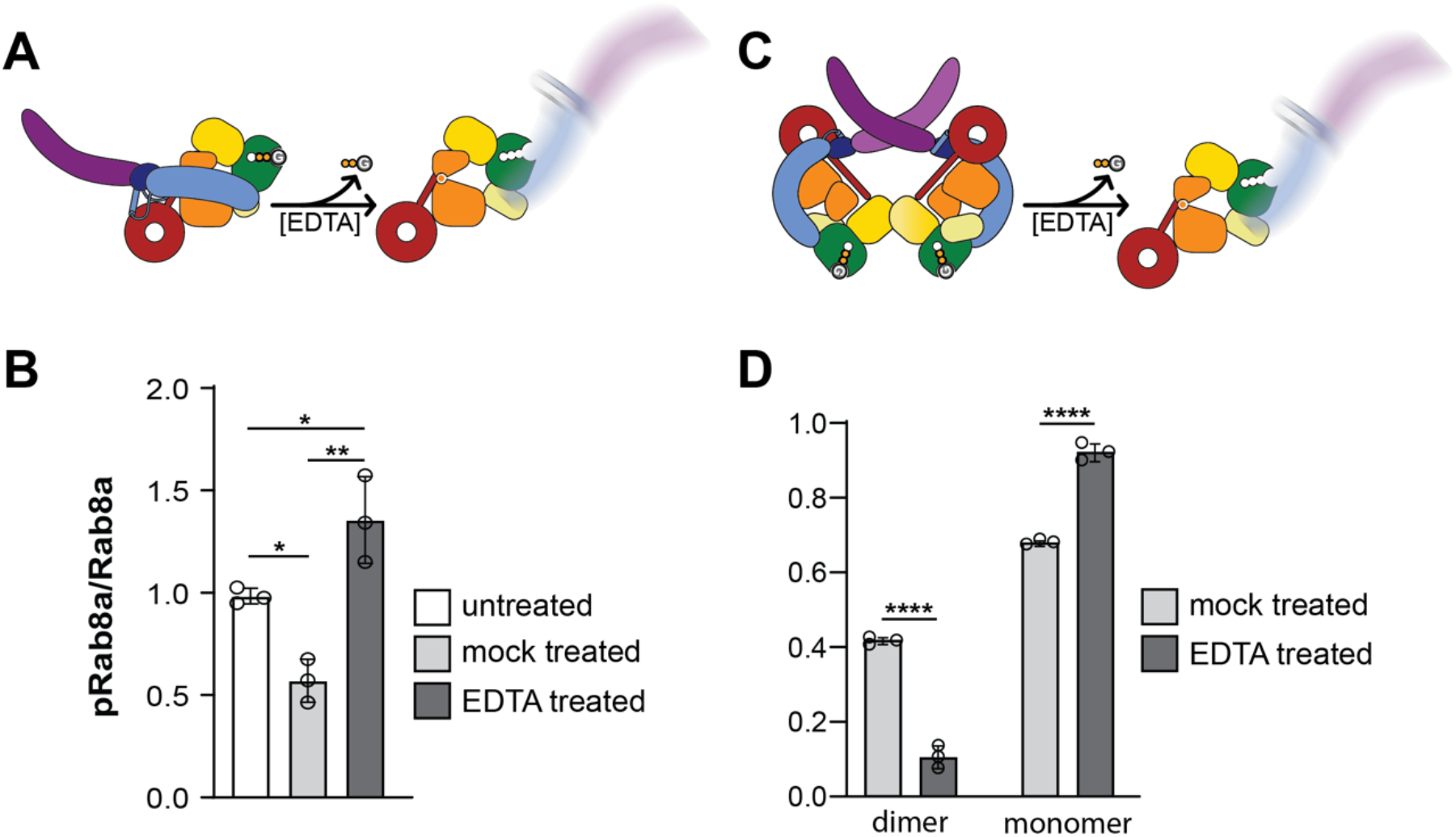
Removal of GDP activates LRRK2 and disrupts the dimer form. **(A)** Schematic representation of the hypothesis being tested: that removal of GDP by exposing LRRK2 to the nucleotide stripping protocol leads to its activation. **(B)** LRRK2[WT] was incubated with (‘EDTA treated) or without EDTA (‘mock treated’) for 24 hours followed by addition of 20mM MgCl_2_ and further incubation for 24 hours. Untreated LRRK2 (‘untreated’) was used immediately upon thawing from frozen aliquots. All LRRK2 samples were incubated with Rab8a in the presence of ATP, with one negative control lacking ATP. Background-corrected pRab8a was normalized to the average pRab8a in the untreated LRRK2 + ATP condition from [n=3] independent Western blots. Images were processed in ImageJ. All data were analyzed using a one-way ANOVA with Tukey’s multiple comparison correction. *p = 0.0246 for untreated vs ‘mock’; *p = 0.0378 for untreated vs ‘EDTA’; **p = 0.0011 for ‘mock’ vs ‘EDTA’. (See Figure S15 for representative immunoblot.) **(C)** Schematic representation of the hypothesis being tested: that removal of GDP by exposing LRRK2 to the nucleotide stripping protocol destabilizes the dimeric form. **(D)** Fraction of LRRK2 monomers and dimers relative to (monomer + dimer) for LRRK2 exposed to the nucleotide stripping protocol or mocked-stripped (see Methods). Total particle counts corresponding to the molecular weights of either monomer or dimer were determined from mass photometry traces. The fraction of dimers contributing to the total (monomer + dimer) number of LRRK2 landing events was analyzed with one-way analysis of variance (ANOVA) and corrected for multiple comparisons using Tukey’s test. Individual data points represent 3 technical repeats. ****p < 0.0001. See Figure S18A for mass photometry traces and particle counts.

One molecular species of LRRK2 that was conspicuously absent from our data sets was a dimer of activated LRRK2s (Figure 1F). We observed dimers of the autoinhibited form, and mixed dimers with one autoinhibited and one activated protomer. However, none of our data sets contained a dimer where both LRRK2 protomers had their repeats undocked. We wondered if activation of LRRK2 might destabilize the dimer. This was indeed the case: activating LRRK2 with the nucleotide-stripping protocol resulted in a shift from dimers to monomers, as measured by mass photometry (Figure 4C and 4D and Figure S18A).

We also solved a structure of the undocked N-terminal repeats (Figure 3L, and Figure S7), which contained a significant fraction of particles coming from monomeric activated LRRK2. Our ability to obtain this structure shows that the repeats remain structured when undocked from the C-terminal half of LRRK2. Importantly, modeling suggested that the N-terminal repeats could not redock when LRRK2 is in its activated conformation (Figure S11). Without LRRK2 going back to the conformation seen in its GDP-bound state, docking would require flexibility in the repeats that we have not seen in our data (Figure S11). Thus, the combination of conformational changes at the ROC-COR interface and the rigidity of the N-terminal repeats appear to be key elements in LRRK2’s regulation.

### The PD-linked R1441C/G/H mutations likely promote undocking of the N-terminal repeats

R1441 is located at the interface between the ROC GTPase and the COR-B domain (Figure 1A, 1B), a region where several PD-linked mutations cluster. To understand how the R1441C/G/H mutations increase Rab phosphorylation by LRRK2^27^, we first compared structures of LRRK2[WT] in its autoinhibited and activated conformations (Figure 3M-3S). In GDP-bound, autoinhibited LRRK2[WT], R1441 packs against the COR-B domain, likely contributing to the stabilization of this interface, and thus to autoinhibition (Figure 3N, 3O). In contrast, the structure of GMPPNP-bound, activated LRRK2[WT] revealed density that can accommodate R1441 in two different rotamers; one pointing towards COR-B and the other away from it (Figure 3P, 3Q); importantly, neither rotamer interacts with COR-B in the activated state, where COR-B has moved away from ROC. We hypothesize that R1441, which is far away from the nucleotide binding pocket and thus unlikely to directly affect binding or hydrolysis, acts as a latch that helps stabilize the ROC-COR interactions as COR-B approaches ROC when LRRK2 returns to its autoinhibited conformation. This interaction between R1441 and COR-B is absent in the PD-linked R1441C mutant (Figure 3R, 3S), which would therefore favor the activated state of LRRK2. A comparison of cryo-EM data obtained from untreated LRRK2 (i.e. not exposed to the nucleotide-stripping protocol) either WT or carrying the R1441C mutation showed a moderate increase in the fraction of particles with their N-terminal repeats undocked in LRRK2[R1441C] (Figure S12). Thus, our data suggest that the increased Rab phosphorylation seen in LRRK2[R1441C/G/H] is a result of a shift in the LRRK2 population towards its activated conformation rather than a direct effect on the catalytic activity of the kinase. A test of this hypothesis is discussed below.

### Development of DARPins to control LRRK2’s activation

The data presented above show that autoinhibited LRRK2 is bound to GDP and that its loss leads to LRRK2 activation. Our data also suggest that the coupling between nucleotide and activation state is mediated by the conformation of the ROC-COR module, whose flexibility is constrained when the N-terminal repeats are docked. Given this, we wondered if the conformation (docked or undocked) of the N-terminal repeats would in turn regulate nucleotide binding and hydrolysis in the ROC domain. To test this, we needed a way to promote or prevent the undocking of the N-terminal repeats that did not require directly exchanging the nucleotide of the GTPase. We had previously identified Designed Ankyrin Repeat Proteins (DARPins) that bound to LRRK2^26^. The first DARPin we characterized^26^, E11, bound to the WD40 domain in an area where it does not interact or interfere with any other part of LRRK2, and did not affect LRRK2 kinase activity in vitro^26^ (Figure 5K). Here, we screened for other DARPins that might alter LRRK2’s activation state and identified two additional DARPins that bind to the WD40 domain: one that activates LRRK2 (C12), and one that stabilizes its autoinhibited conformation (G10) (Figure 5). DARPin C12 binds at a location on the WD40 domain that is sterically incompatible with the docked conformation of the N-terminal repeats (Figure 5A-4D, Figure S13). DARPin G10 binds both to the WD40 domain and to the N-terminal repeats, bridging them (Figure 5E-4H, Figure S14). As predicted from the structures (Figure 5I, 4J), DARPin C12 increased LRRK2s kinase activity towards Rab8a in vitro, whereas DARPin G10 decreased it (Figure 5K, Figure S15).

**Figure 5.**
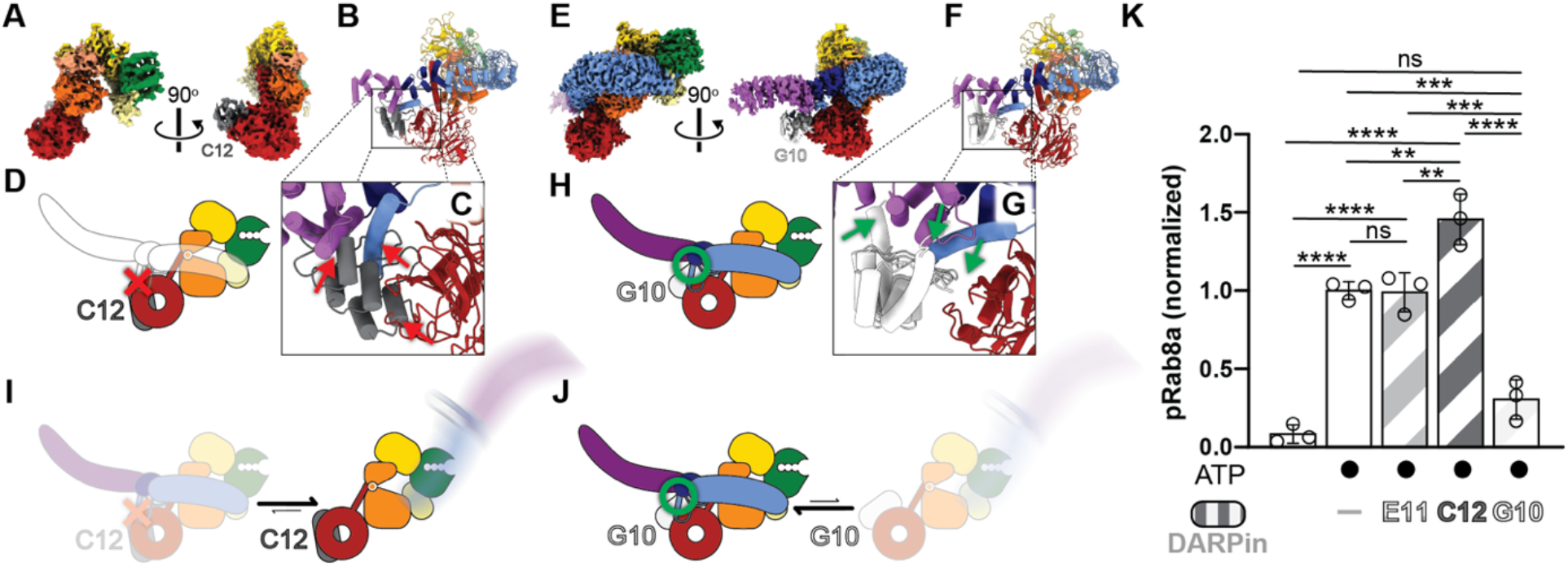
Cryo-EM structures of LRRK2 and LRRK2^RCKW^ bound to DARPins C12 and G10. **(A)** Cryo-EM map of LRRK2^RCKW^ with DARPin C12 bound to its WD40 domain. **(B, C)** Model of full-length LRRK2 bound to C12 showing the clashes that would result between the N-terminal repeats and C12 (B), and close-up of those clashes (C). **(D)** Caroon illustration of the clashes. **(E)** Cryo-EM map of LRRK2 with DARPin G10 bound to the WD40 domain. **(F, G)** Structure of LRRK2 bound to G10 showing that G10 interacts both with the WD40 domain and the N-terminal repeats (F), and close-up of the interactions (G). **(H)** Cartoon illustration of the interactions. **(I)** DARPin C12 is predicted to activate LRRK2 by promoting the conformation with the N-terminal repeats undocked. **(J)** DARPin G10 is predicted to inhibit LRRK2 by stabilizing the conformation with the N-terminal repeats docked. **(K)** In vitro Rab8a phosphorylation assays. LRRK2[WT] was incubated with Rab8a in the presence of ATP (with a no ATP control) plus one of the following: no DARPin, E11, C12, or G10. Background-corrected pRab8a was normalized to the average of pRab8a in the LRRK2 + ATP condition from [n=3] independent Western blots. Images were processed in ImageJ; all data were analyzed using a one-way ANOVA with Tukey’s multiple comparison correction. p = non-significant for No ATP vs LRRK2 + G10, all other comparisons with No ATP ****p<0.0001. LRRK2 vs LRRK2+E11 were non-significant; **p=0.0042 for LRRK2 vs LRRK2+C12; ***p=0.0001 for LRRK2 vs LRRK2+G10; **p=0.0036 for LRRK2+E11 vs LRRK2+C12; ***p=0.0002 for LRRK2+E11 vs LRRK2+G10; and ****p<0.0001 for LRRK2+C12 vs LRRK2+G10.

### G2019S and R1441H increase Rab phosphorylation by LRRK2 via different mechanisms

Our structural analysis suggested that R1441C/G/H mutations activate LRRK2 by favoring the undocked conformation of the N-terminal repeats (Figure 3M-3S). G2019S, on the other hand, is in the catalytic DYG triad in the kinase’s active site. Thus, we hypothesized that these two PD-linked mutations lead to increased Rab phosphorylation in cells by acting in different steps of the LRRK2 activation/phosphorylation pathway (Figure 6A): R1441C/G/H mutations enrich the population of activated LRRK2, while G2019S activates the kinase itself. We tested this in two different ways.

**Figure 6.**
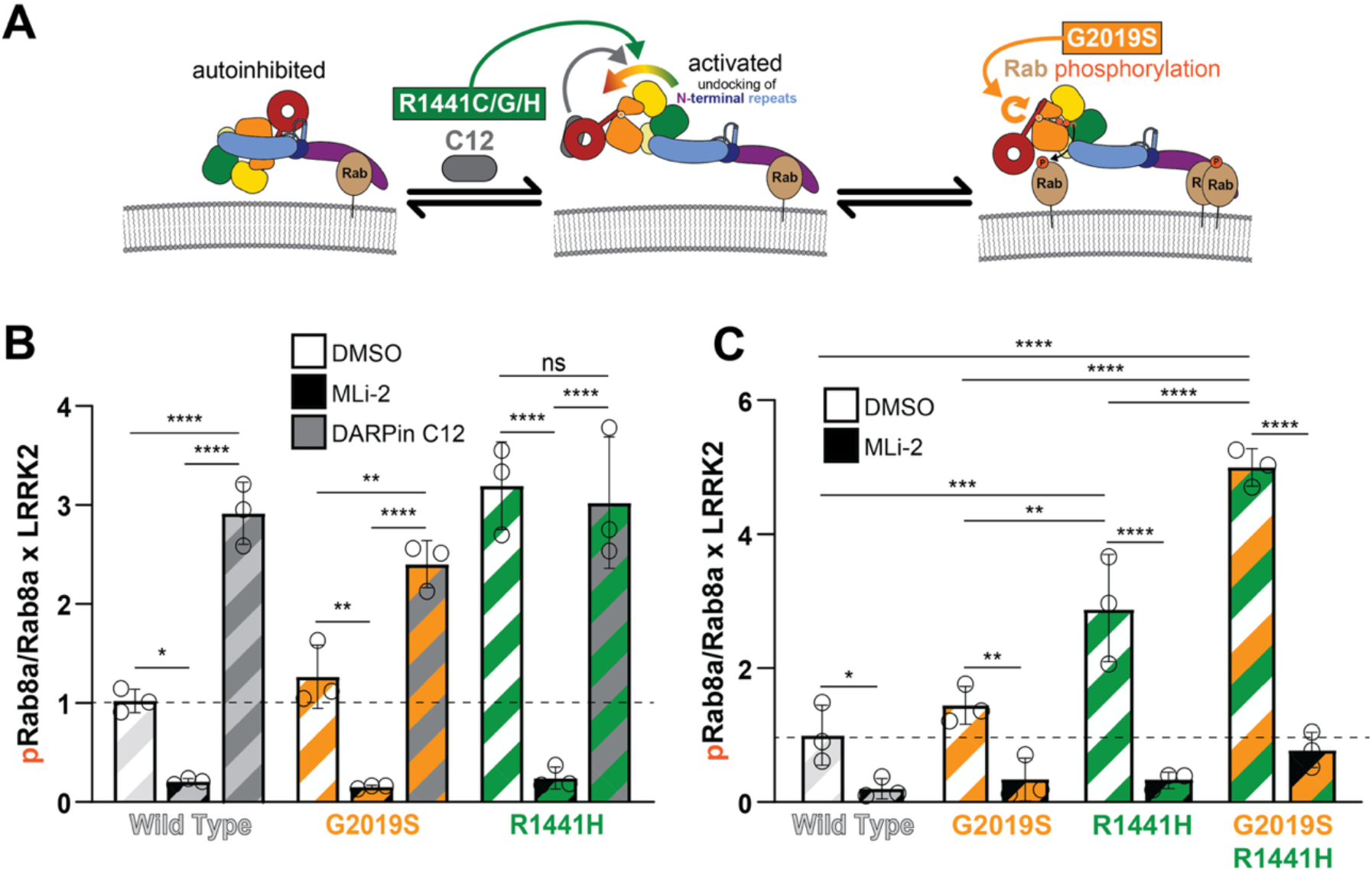
The PD mutants R1441C/G/H and G2019S increase Rab phosphorylation by different mechanisms. **(A)** Proposed mechanism of activation of LRRK2 by the PD-linked mutations R1441C/G/H and G2019S. R1441C/G/H act by promoting the release of autoinhibition, while G2019S increases the activity of the kinase itself. Like R1441C/G/H, DARPin C12 also promotes the release of autoinhibition. **(B)** Quantification of pRab8a (pT72) / Rab8a x LRRK2, normalized to the mean of LRRK2[WT] + DMSO condition from [n=3] independent Western blots. HEK 293T co-expressing GFP-11-tagged LRRK2[WT], LRRK2[G2019S], or LRRK2[R1441H] and GFP-Rab8a with or without Flag-tagged DARPin C12 were treated with either DMSO, MLi-2, or nothing in the C12 condition. Samples were immunoblotted for LRRK2, phospho-Rab8a (pT72), GFP, and Flag. Images were processed on ImageJ, and all data were analyzed using a two-way ANOVA with Tukey’s multiple comparison correction. ****p<0.0001 for WT+C12 vs DMSO and MLi-2 and *p=0.0161 for WT+DMSO vs MLi-2; **p=0.0011 and ****p<0.0001 G2019S+C12 vs DMSO and MLi-2; **p=0.0013 G2019S+DMSO vs MLi-2. ****p<0.0001 R1441H+MLi-2 vs C12 and DMSO; non-significant R1441H DMSO vs C12. Error bars ± SD. **(C)** Quantification of pRab8a (pT72) / Rab8a x LRRK2 normalized to the mean of LRRK2[WT] + DMSO condition. HEK 293T co-expressing LRRK2[WT], LRRK2[G2019S], LRRK2[R1441H] or the double mutant LRRK2[G2019S/R1441H] and GFP-Rab8a were treated, processed and analyzed as before (B). DMSO vs MLi-2: *p=0.0229 WT, **p=0.0031 G2019S, and ***p<0.0001 R1441H and DM; non-significant G2019S+DMSO vs WT+DMSO; **p=0.0016 G2019S+DMSO vs R1441H+DMSO; ***p=0.0001 R1441H+DMSO vs WT+DMSO; ****p<0.0001 all comparisons with the G2019S/R1441H double mutant. Error bars ± SD.

First, we used the C12 DARPin, which activates LRRK2 by favoring the undocking of the N-terminal repeats. Mirroring our results in vitro (Figure 5K), expressing DARPin C12 in cells led to a significant increase in the phosphorylation of Rab8a by LRRK2[WT] (Figure 6B). This effect is not exclusive to Rab8a; we observed a similar increase in the phosphorylation of Rab10 (Figure S16). If R1441C/G/H mutations increase activity by favoring undocking of the repeats, as we hypothesized, the presence of C12 should not lead to a significant additional increase in Rab phosphorylation. LRRK2 G2019S, on the other hand, acts on the kinase itself and should be further activated by C12. This was indeed the case when we measured Rab8a phosphorylation in cells co-expressing C12 and either LRRK2[G2019S] or LRRK2[R1441H] (Figure 6B and Figure S17A).

A second prediction from our hypothesis (Figure 6A) is that the activating effects of the R1441C/G/H and G2019S mutations should be additive to some extent, as they act in series, with R1441C/G/H mutations increasing the pool of activated LRRK2, and G2019S increasing the kinase activity of that pool. Indeed, the increase in Rab8a phosphorylation in cells by the double mutant LRRK2[R1441H/G2019S] was close to the sum of the effects seen with the single mutants (Figure 6C and Figure S17B).

Taken together, these data support the hypothesis that even though R1441C/G/H and G2019S both lead to an increase in Rab phosphorylation in cells, they do so by acting on different steps in the LRRK2 pathway.

### Nucleotide exchange and hydrolysis by ROC are regulated both by the N-terminal repeats and the Rab substrate

We next tested the effect of DARPins C12 and G10 on the GTPase activity of LRRK2. Our hypothesis (Figure 7A) was that G10 would stabilize the autoinhibited conformation of LRRK2 and thus prevent cycles of nucleotide exchange and hydrolysis, while C12 would stimulate them by promoting the undocking of the N-terminal repeats. Our initial data agreed with our prediction that G10 would inhibit the GTPase activity of LRRK2; however, C12 had no effect on it (Figure 7B). We wondered whether undocking of the N-terminal repeats was not sufficient to stimulate GTP hydrolysis, and if adding a Rab substrate might. Indeed, the combination of C12 and Rab8a (the Q67L variant that is locked in its GTP-bound state) led to a significant activation of the GTPase (Figure 7C). Importantly, Rab8a[Q67L] on its own had no effect on the baseline GTPase activity of LRRK2 (Figure 7C), suggesting that undocking of the N-terminal repeats by C12 is required for Rab8a to exert its effect on hydrolysis.

**Figure 7.**
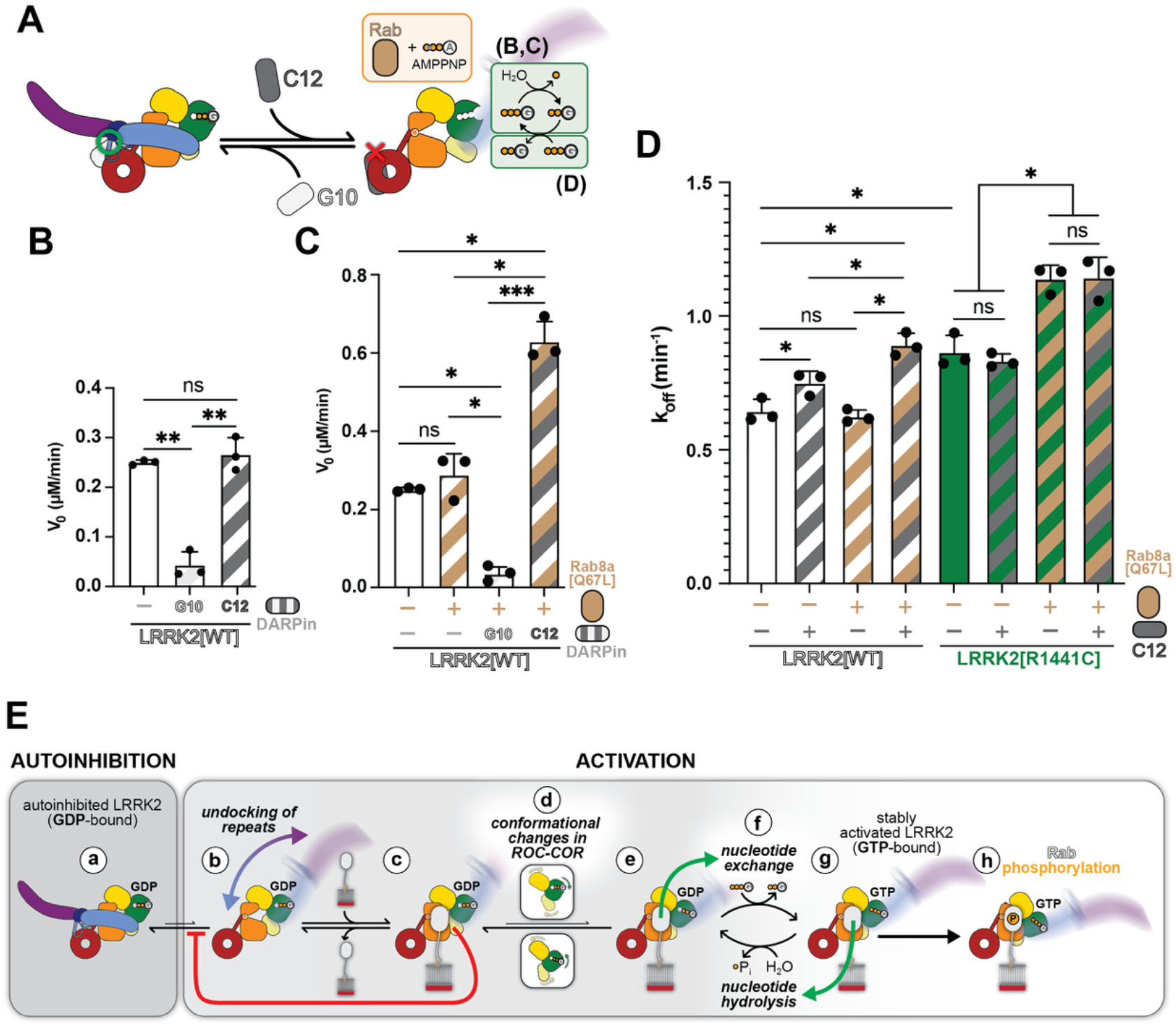
The conformation of the N-terminal repeats regulates LRRK2’s GTPase. **(A)** Schematic of the GTPase and GDP release assays. LRRK2’s rate of GTP hydrolysis (B and C) or GDP release (D) were measured for LRRK2 alone or with the addition of DARPin C12 or G10 and/or the substrate Rab8a (the GTP-locked mutant Q67L). DARPin C12 promotes the activated form of LRRK2 while G10 stabilizes the autoinhibited conformation (see Figure 5). **(B)** Initial GTP hydrolysis velocities (V_0_, µM/min) of LRRK2-catalyzed reactions were measured in the presence of G10 or C12. Data are shown as mean ± SEM from [n = 3] independent experiments. Statistical analysis was performed using ANOVA with post hoc test / Student’s t-test; **p < 0.01, ns = not significant. **(C)** Initial GTP hydrolysis velocities (V_0_, µM/min) of LRRK2-catalyzed reactions were measured with Rab8 Q67L alone, or with Rab8 Q67L in combination with G10 or C12. Data are presented as mean ± SEM from [n = 3] independent experiments. Statistical analysis was performed using ANOVA with post hoc test / Student’s t-test; ***p < 0.001, *p < 0.05, ns = not significant. **(D)** GDP dissociation rates (k_off_) for WT and LRRK2[R1441C]. Measurements were performed with or without DARPin C12 and with and without Rab8a[Q67L]. Data are presented as mean ± SEM from [n = 3] independent experiments. Statistical analysis was performed using [ANOVA with post hoc test / Student’s t-test]; *p < 0.05, ns = not significant. **(E)** Schematic representation of our model for how LRRK2’s GTPase controls its activation and autoinhibition. (a) Autoinhibited LRRK2 is in the GDP-bound state. (b) The N-terminal repeats can occasionally undock, but this state is disfavored in the GDP-bound state. (c) Binding of a Rab substrate to LRRK2 sterically prevents re-docking of the N-terminal repeats (red arrow), transiently stabilizing the undocked conformation. Once undocking of the N-terminal repeats relieves the conformational constraints on the ROC-COR module (d), the latter can switch to the active conformation (e). This conformational change in ROC-COR lowers the affinity for GDP in the ROC GTPase, leading to GDP release and binding of GTP, which is more abundant in the cell (f). The Rab substrate stimulates the release of GDP (green arrow in (e)). Binding of GTP stabilizes the undocked conformation of LRRK2 (g). The Rab substrate also stimulates GTP hydrolysis (green arrow in (g)), which returns LRRK2 to the GDP-bound state and would allow re-docking of the N-terminal repeats if a Rab substrate is not present to sterically prevent re-docking of the N-terminal repeats. The undocked, GTP-bound form of LRRK2 is responsible for the bulk of the Rab phosphorylation (h).

The effect of Rab8a on the rate of GTP hydrolysis could be the result of an increase in hydrolysis itself, an increase in the rate of GDP release, or a combination of both. To address this, we measured the rate of GDP release from LRRK2 pre-loaded with fluorescently labeled nucleotide in a single-turnover assay (see Methods). Consistent with our model where undocking of the N-terminal repeats lowers the affinity for GDP, the rate of GDP release from LRRK2[WT] was increased by the addition of DARPin C12 (Figure 7D). Addition of Rab8a on its own had no effect on GDP release, but the combination of C12 and Rab8a led to a small but significant increase relative to C12 alone, indicating that Rab8a is promoting GDP release (Figure 7D).

Lastly, we wanted to confirm that the effects we were seeing with C12 were due to its promoting the undocked conformation of LRRK2. For this, we turned to LRRK2[R1441C], which our data showed favors the undocked conformation of LRRK2 (Figure 6). Consistent with those results, GDP release from LRRK2[R1441C] was faster than from LRRK2[WT] and was in turn insensitive to the addition of C12 (Figure 7D). Finally, addition of Rab8a to LRRK2[R1441C] significantly increased GDP release, and was again insensitive to C12, supporting the idea that Rab8a is acting downstream of the undocking of the N-terminal repeats. Taken together, our data showed that Rab8a stimulates both GDP release and GTP hydrolysis in activated LRRK2.

## DISCUSSION

### The nucleotide state of the ROC GTPase controls LRRK2 activation and autoinhibition

The work presented here, together with contributions from several other groups in the field, has led us to propose a model for how the nucleotide state of the ROC domain regulates the activation and autoinhibition of LRRK2 (Figure 7E). Our data also showed that the common PD-linked mutations R1441C/G/H and G2019S act on different steps of the activation mechanism and provided an explanation for how PD-associated mutations found on either side of this ROC-COR interface (e.g. N1437H, Y1699C) lead to increased Rab phosphorylation. Our structural data showed that the autoinhibited conformation of LRRK2 was always accompanied by a ROC domain bound to GDP (Figure 2), in agreement with all structures of this species published to date where the nucleotide state could be determined^9,16,17^. This suggests that the docked conformation of the N-terminal repeats stabilizes a conformation of ROC with high affinity for GDP (Figure 7E(a)). This is consistent with our observation that complete stripping of GDP from autoinhibited LRRK2 requires long treatment with EDTA (Figure 2H and. Figure S8).

Our model proposes that undocking of the N-terminal repeats and the resulting increased conformational flexibility in the ROC-COR module leads to a decrease in the ROC domain’s affinity for GDP (Figure 7E(b-e)). This is supported by the increase in the rate of GDP release from LRRK2[R1441C], a mutant we have shown favors the undocked conformation of the N-terminal repeats (Figure 3N-3S and Figure 6). It is likely that any protein that interacts with LRRK2 and stabilizes its N-terminal domains in the undocked conformation will act as an activator by promoting nucleotide exchange, leading to stably active GTP-bound LRRK2. While a few activators have been identified—including Rab12^28,29^, Rab32^30,31^, and GABARAP^32,33^—we do not yet understand whether they exert their effects by recruiting LRRK2 to membranes or additionally by somehow stabilizing the undocked conformation once LRRK2 is on the membrane.

Our data also revealed that the Rab substrate plays a role in stimulating the GTPase cycle in the ROC domain, accelerating both GDP release and GTP hydrolysis (Figure 7C and 7D, Figure 7E(e-g)). This mechanism allows the N-terminal repeats to act as a substrate sensor; promoting GTP hydrolysis to return LRRK2 to its GDP-bound state puts LRRK2 in a state compatible with a return to autoinhibition, a process that is prevented if Rab substrate is available to sterically block redocking of the N-terminal repeats (Figure 7E(c)). If re-docking is prevented, ROC-COR remains in the conformation with low affinity for GDP (Figure 7E(e)), leading to GDP release, GTP binding, and a return of LRRK2 to its stably active conformation (Figure 7E(g-h)).

Several studies over the years have addressed the role of the GTPase in regulating LRRK2 activity. While some are consistent with the work we presented here, others are not. In general, work done in cells or with lysates agrees with our observations: addition of GTP or non-hydrolyzable GTP analogs to cell lysates correlates with increased LRRK2 kinase activity^11,12^. Similarly, LRRK2 localization and association with microtubules in cells correlate more closely with the ability to bind GTP than with kinase activity^13^, and we now understand that binding to microtubules requires LRRK2 to be in its active conformation^10,34,35^. Also in agreement with our data are observations that pathogenic mutations in the ROC–COR region can enhance GTP binding ^11,13^ and affect GTP hydrolysis^14,15^, and that disruption of guanine nucleotide binding by mutations in the P-loop of the GTPase domain (e.g., K1347A or T1348N) generally impaired kinase activity and showed cellular phenotypes across multiple systems^12,36,37^. However, while nucleotide-dependent effects were frequently observed in lysate-based assays, they were reduced or absent when nucleotides were added to immunoprecipitated or purified LRRK2, indicating a strong dependence on experimental and cellular context^11,12^. In addition, measurements of GTPase activity have yielded inconsistent results: while some studies reported low intrinsic hydrolysis and predominantly GTP-bound LRRK2 in cells^36^, others showed that mutations that either increase or decrease GTP hydrolysis reduce kinase activity^37^, suggesting that nucleotide binding is more important than hydrolysis in regulating kinase activity. Finally, there is evidence pointing to a role for autophosphorylation of the ROC domain in regulating GTP hydrolysis^15^. It is difficult to determine where discrepancies arise from as the studies are not directly comparable. They differ in whether work was done in cells, with lysates, or, when using purified protein, on whether LRRK2 was immunoprecipitated or recombinantly expressed. They also report on a variety of assays (nucleotide binding, hydrolysis, Michaelis-Menten kinetics) and both auto- and substrate phosphorylation. Our work has shown that the activation state of LRRK2 is sensitive to how the protein is treated in guanine nucleotide exchange protocols. Given this, it is likely that some of the disagreements in the literature reflect differences in the distribution of active and autoinhibited populations of LRRK2 in cells and lysates vs. purified protein, or because of differences in the nucleotide exchange protocols used.

### The active form of LRRK2 is likely a monomer

Several lines of evidence lead us to propose that the active form of LRRK2 is a monomer. Previous work has shown that disruption of dimerization is activating^7,35^. Stripping LRRK2 of its bound GDP, a process that is activating and leads both to undocking of the N-terminal repeats (Figure S10) and an increase in kinase activity in vitro (Figure 4A and 4B, and Figure S11), correlates with a reduction in the dimeric form of LRRK2 (Figure 4C and 4D, and Figure S18A). Finally, in agreement with our mass photometry data, modeling a dimer in which both LRRK2s are activated showed that the conformational changes in the COR-A and COR-B domains involved in activation lead to significant changes and clashes at the dimer interface (Figure S18B-S18H).

In contrast to our observations and proposal, it has been suggested that LRRK2 is activated in the context of a tetramer^18^. This model was based on a cryo-EM structure where LRRK2 formed a dimer of dimers, with each dimer containing one protomer of autoinhibited LRRK2 and one protomer of activated LRRK2, where its repeats were undocked. LRRK2 is known to dimerize and structural work suggests that the highest affinity dimerization interface is the one mediated by interactions between its C-terminal Of ROC (COR) domains (COR-A and COR-B)^9,10^ (Figure 1D); this was the interface mediating the dimers that formed the tetramer. Given this, the model predicts that disrupting dimerization at the COR interface, which in turn would disrupt the tetramer, should result in a decrease in LRRK2 activity. However, the opposite is true: disrupting the dimer interface increases Rab phosphorylation, both in vitro^7^ and in cells^7,35^. While the physiological relevance of the reported tetramer remains to be determined, it is possible that its appearance was a consequence of the high concentrations of LRRK2 used for cryo-EM (21-28μM)^18^.

We hypothesize that one role of dimerization in LRRK2 may be, as we discuss further below, to stabilize the autoinhibited state by preventing the conformational changes in the ROC-COR module linked to activation.

### The ROC-COR module as a regulatory nexus

Our work points to the ROC-COR module as a regulatory nexus in LRRK2, with the conformation of ROC-COR being coupled to the nucleotide state of the ROC GTPase. A recent cryo-EM structure of a bacterial Roco protein (which contain the LRR-ROC-COR module but lacks a kinase) showed undocking of its LRR domain when its GTPase was in a GTP-like state^38^. We propose that stabilizing the ROC-COR module in a specific conformation, as we suggest LRRK2 dimerization does for the autoinhibited form, may be a strategy that other regulatory factors use. Identifying such factors may lead to novel targets for PD therapeutics.

The best-established regulators of LRRK2 are 14-3-3 proteins^39–42^. These proteins, which function as dimers, bind to phosphorylated residues (pS910 and pS935) in a flexible loop in the LRR domain, and are understood to keep LRRK2 in its autoinhibited form in the cytosol^21^. The pS910 and pS935 marks are used as biomarkers in the diagnosis of LRRK2-linked PD, as the majority of pathogenic mutations that increase LRRK2 activity suppress biomarker site phosphorylation^22^.

A structure of LRRK2 bound to 14-3-3 was recently reported^19^. In this structure, a 14-3-3 dimer was bound simultaneously to the COR-A and COR-B domains of LRRK2 and densities for two phosphoserines were observed interacting with 14-3-3. The authors proposed that 14-3-3 maintains LRRK2 in an autoinhibited state by restricting the movement of the N-terminal repeats, as these are tethered to 14-3-3 via the loop containing pS910 and pS935. The authors also noted that 14-3-3 could only fully engage with autoinhibited LRRK2; the conformational changes in COR-A and COR-B during activation would prevent 14-3-3 from binding to both in the active form of LRRK2^19^.

Based on the data presented here, we propose that the role of the pS910/pS935 residues in the loop is not to restrict the conformation of the N-terminal repeats, but rather to recruit 14-3-3 to LRRK2, as the loop is long enough to accommodate undocking even when bound to 14-3-3. In this model, 14-3-3’s main function would be to clamp together the COR-A and COR-B domains to prevent the conformational changes that lead to GDP release and LRRK2 activation.

Interestingly, binding of 14-3-3 to LRRK2 prevents dimerization by blocking the COR-COR interface^19^. Since LRRK2 dimerization and 14-3-3 binding both engage COR-A and COR-B in their autoinhibited conformation, they could be alternative mechanisms to prevent the conformational changes required for LRRK2 activation. Understanding the interplay among these different regulatory mechanisms could also lead to novel therapeutic targets.

### Nucleotide control of the docking and undocking of the N-terminal repeats

Our structural data suggest a mechanistic model for how the nucleotide state of ROC might control the docking and undocking of the N-terminal repeats (Figure S19). The LRR are connected to the ROC-COR module in the autoinhibited state of LRRK2 via a short two-stranded β-sheet (Figure S19A, S19B). At one end, both β-strands connect to the ROC domain. At the other end, they connect to different domains: one goes from the LRR to ROC, and the other from ROC to COR-A. This arrangement means that changes in the relative orientations of ROC and COR-A, as happens during activation of LRRK2, would destabilize this β-sheet. In fact, the β-sheet is absent from all our structures of activated LRRK2. We only saw it in the structure of GMPPNP-bound activated LRRK2 where partial density for the LRR is present (Figure S19C-S19F). The potential importance of this β-sheet is highlighted by the high conservation of the sequences that encompass it (Figure S19G, S19H). A closer look at changes in the ROC domain that correlate with its nucleotide state suggests that a conformational switch may be controlling the presence or absence of the β-sheet, with GDP stabilizing it and GMPPNP leading to its disruption (Figure S19I-S19L).

### Roadmap for therapeutics

The conformation of the ROC-COR module changes when the N-terminal repeats undock and no longer constrain it (Figure 7E(c-e)). Additional changes in the interfaces between the ROC and COR domains occur in response to the nucleotide bound to ROC (Figure 3G-3J). These changes provide a roadmap for identifying small molecules that stabilize either the GDP-bound autoinhibited LRRK2 or the GTP-bound or Apo activated LRRK2. The importance of the interface between ROC and COR-B for PD has been appreciated for a long time as it is a hot spot for disease-linked mutations (Figure 1A, 1B). Molecules that stabilize the autoinhibited LRRK2 would represent a novel category of allosteric LRRK2 kinase inhibitors for the treatment of PD, distinct from the Type-1 kinase inhibitors that are and were previously in clinical trials (clinicaltrials.gov). Molecules that activate LRRK2 could also be interesting therapeutically, as loss of LRRK2 activity has been linked to both lung fibrosis and lung tumorigenesis in rodents^23–25^.

## Author Contributions

All authors were involved in experimental design. AVS, TB, MS-M, EM, KHVN, and DR performed the cryo-EM work (data collection and analysis). KSH and KJS performed the cell-based assays. WG performed the GTPase and GDP release assays. EM performed the mass photometry experiments. LZ and KMS developed the nucleotide-exchange assay. AVS, KSH, TB, MS-M, KS, EX, VD, SM, SK, SLR-P, and AEL were involved in the development and characterization of the DARPins. AVS, KSH, TB, MS-M, EM, KHVN, WG, KS, RF, DR, SLR-P, and AEL performed data analysis. JI made the model animation. AVS, SLR-P, and AEL wrote the manuscript, and AVS, KSH, TB, KJS, SK, KMS, SLR-P, and AEL edited it.

## Acknowledgments

We thank the Cryo-EM Facility and the Physics Computing Facility at UC San Diego. We also thank our funding sources: AVS was supported by the Molecular Biophysics Training Grant (NIH grant T32 GM139795) and a Postgraduate Fellowship from the National Science Foundation (DGE-2038238); KSH is supported by an F31 Fellowship from the National Institutes of Health (F31NS155649); TB and KJS are supported by postdoctoral fellowships from the Parkinson’s Foundation (PRF-1443704 and PF-PRF-1439460, respectively); SK is funded by the Structural Genomics Consortium (SGC), a registered charity (No:1097737) that received funds from Bayer AG, Boehringer Ingelheim, Bristol Myers Squibb, Genentech, Genome Canada through Ontario Genomics Institute, (OGI-196), EU/EFPIA/OICR/McGill/KTH/Diamond Innovative Medicines Initiative 2 Joint Undertaking (EUbOPEN grant 875510), Janssen, Merck KGaA, Pfizer, and Takeda. KMS was funded by MJFF; KMS and SLR-P were funded by the Howard Hughes Medical Institute; SK, KMS, SLR-P and AEL were supported by the LRRK2 Investigative Therapeutics Exchange (MJFF-LITE).

## Data availability

All cryo-EM maps and models have been deposited in the EM Data Bank and Protein Data Bank, respectively. All accession numbers are listed in Table S1.

## REFERENCES

1. Krüger, C., Lim, S.-Y., Buhrmann, A., Fahrig, F.L., Gabbert, C., Bahr, N., Madoev, H., Marras, C., Klein, C., and Lohmann, K. (2025). Updated MDSGene review on the clinical and genetic spectrum of LRRK2 variants in Parkinson’s disease. NPJ Parkinsons Dis 11, 30. 10.1038/s41531-025-00881-9.

2. Steger, M., Tonelli, F., Ito, G., Davies, P., Trost, M., Vetter, M., Wachter, S., Lorentzen, E., Duddy, G., Wilson, S., et al. (2016). Phosphoproteomics reveals that Parkinson’s disease kinase LRRK2 regulates a subset of Rab GTPases. eLife 5, 809.

3. Steger, M., Diez, F., Dhekne, H.S., Lis, P., Nirujogi, R.S., Karayel, O., Tonelli, F., Martinez, T.N., Lorentzen, E., Pfeffer, S.R., et al. (2017). Systematic proteomic analysis of LRRK2-mediated Rab GTPase phosphorylation establishes a connection to ciliogenesis. eLife 6.

4. Hutagalung, A.H., and Novick, P.J. (2011). Role of Rab GTPases in membrane traffic and cell physiology. Physiol Rev 91, 119–149. 10.1152/physrev.00059.2009.

5. West, A.B., Moore, D.J., Biskup, S., Bugayenko, A., Smith, W.W., Ross, C.A., Dawson, V.L., and Dawson, T.M. (2005). Parkinson’s disease-associated mutations in leucine-rich repeat kinase 2 augment kinase activity. 102, 16842–16847.

6. Gloeckner, C.J., Kinkl, N., Schumacher, A., Braun, R.J., O’Neill, E., Meitinger, T., Kolch, W., Prokisch, H., and Ueffing, M. (2006). The Parkinson disease causing LRRK2 mutation I2020T is associated with increased kinase activity. Human molecular genetics 15, 223–232.

7. Kalogeropulou, A.F., Purlyte, E., Tonelli, F., Lange, S.M., Wightman, M., Prescott, A.R., Padmanabhan, S., Sammler, E., and Alessi, D.R. (2022). Impact of 100 LRRK2 variants linked to Parkinson’s disease on kinase activity and microtubule binding. Biochem J 479, 1759–1783. 10.1042/BCJ20220161.

8. Di Maio, R., Hoffman, E.K., Rocha, E.M., Keeney, M.T., Sanders, L.H., De Miranda, B.R., Zharikov, A., Van Laar, A., Stepan, A.F., Lanz, T.A., et al. (2018). LRRK2 activation in idiopathic Parkinson’s disease. Science translational medicine 10.

9. Myasnikov, A., Zhu, H., Hixson, P., Xie, B., Yu, K., Pitre, A., Peng, J., and Sun, J. (2021). Structural analysis of the full-length human LRRK2. Cell 184, 3519–3527.e10.

10. Deniston, C.K., Salogiannis, J., Mathea, S., Snead, D.M., Lahiri, I., Matyszewski, M., Donosa, O., Watanabe, R., Böhning, J., Shiau, A.K., et al. (2020). Structure of LRRK2 in Parkinson’s disease and model for microtubule interaction. Nature 588, 344–349.

11. West, A.B., Moore, D.J., Choi, C., Andrabi, S.A., Li, X., Dikeman, D., Biskup, S., Zhang, Z., Lim, K.-L., Dawson, V.L., et al. (2007). Parkinson’s disease-associated mutations in LRRK2 link enhanced GTP-binding and kinase activities to neuronal toxicity. Human molecular genetics 16, 223–232.

12. Taymans, J.-M., Vancraenenbroeck, R., Ollikainen, P., Beilina, A., Lobbestael, E., De Maeyer, M., Baekelandt, V., and Cookson, M.R. (2011). LRRK2 kinase activity is dependent on LRRK2 GTP binding capacity but independent of LRRK2 GTP binding. PLoS ONE 6, e23207.

13. Blanca Ramírez, M., Lara Ordóñez, A.J., Fdez, E., Madero-Pérez, J., Gonnelli, A., Drouyer, M., Chartier-Harlin, M.-C., Taymans, J.-M., Bubacco, L., Greggio, E., et al. (2017). GTP binding regulates cellular localization of Parkinson’s disease-associated LRRK2. Human molecular genetics 26, 2747–2767.

14. Guo, L., Gandhi, P.N., Wang, W., Petersen, R.B., Wilson-Delfosse, A.L., and Chen, S.G. (2007). The Parkinson’s disease-associated protein, leucine-rich repeat kinase 2 (LRRK2), is an authentic GTPase that stimulates kinase activity. Experimental cell research 313, 3658–3670.

15. Gilsbach, B.K., Ho, F.Y., Riebenbauer, B., Zhang, X., Guaitoli, G., Kortholt, A., and Gloeckner, C.J. (2024). Intramolecular feedback regulation of the LRRK2 Roc G domain by a LRRK2 kinase-dependent mechanism. Elife 12, RP91083. 10.7554/eLife.91083.

16. Sanz Murillo, M., Villagran Suarez, A., Dederer, V., Chatterjee, D., Alegrio Louro, J., Knapp, S., Mathea, S., and Leschziner, A.E. (2023). Inhibition of Parkinson’s disease-related LRRK2 by type I and type II kinase inhibitors: Activity and structures. Sci Adv 9, eadk6191. 10.1126/sciadv.adk6191.

17. Zhu, H., Hixson, P., Ma, W., and Sun, J. (2024). Pharmacology of LRRK2 with type I and II kinase inhibitors revealed by cryo-EM. Cell Discov 10, 10. 10.1038/s41421-023-00639-8.

18. Zhu, H., Tonelli, F., Turk, M., Prescott, A., Alessi, D.R., and Sun, J. (2023). Rab29-dependent asymmetrical activation of leucine-rich repeat kinase 2. Science 382, 1404–1411. 10.1126/science.adi9926.

19. Martinez Fiesco, J.A., Beilina, A., Alvarez de la Cruz, A., Li, N., Metcalfe, R.D., Cookson, M.R., and Zhang, P. (2025). 14-3-3 binding maintains the Parkinson’s associated kinase LRRK2 in an inactive state. Nat Commun 16, 7226. 10.1038/s41467-025-62337-1.

20. Kett, L.R., Boassa, D., Ho, C.C.-Y., Rideout, H.J., Hu, J., Terada, M., Ellisman, M., and Dauer, W.T. (2012). LRRK2 Parkinson disease mutations enhance its microtubule association. Human molecular genetics 21, 890–899.

21. Dzamko, N., Deak, M., Hentati, F., Reith, A.D., Prescott, A.R., Alessi, D.R., and Nichols, R.J. (2010). Inhibition of LRRK2 kinase activity leads to dephosphorylation of Ser(910)/Ser(935), disruption of 14-3-3 binding and altered cytoplasmic localization. Biochemical Journal 430, 405–413.

22. Doggett, E.A., Zhao, J., Mork, C.N., Hu, D., and Nichols, R.J. (2012). Phosphorylation of LRRK2 serines 955 and 973 is disrupted by Parkinson’s disease mutations and LRRK2 pharmacological inhibition. Journal of Neurochemistry 120, 37–45. 10.1111/j.1471-4159.2011.07537.x.

23. Tian, Y., Lv, J., Su, Z., Wu, T., Li, X., Hu, X., Zhang, J., and Wu, L. (2021). LRRK2 plays essential roles in maintaining lung homeostasis and preventing the development of pulmonary fibrosis. Proc Natl Acad Sci U S A 118, e2106685118. 10.1073/pnas.2106685118.

24. Herzig, M.C., Kolly, C., Persohn, E., Theil, D., Schweizer, T., Hafner, T., Stemmelen, C., Troxler, T.J., Schmid, P., Danner, S., et al. (2011). LRRK2 protein levels are determined by kinase function and are crucial for kidney and lung homeostasis in mice. Hum Mol Genet 20, 4209–4223. 10.1093/hmg/ddr348.

25. Lebovitz, C., Wretham, N., Osooly, M., Milne, K., Dash, T., Thornton, S., Tessier-Cloutier, B., Sathiyaseelan, P., Bortnik, S., Go, N.E., et al. (2021). Loss of Parkinson’s susceptibility gene LRRK2 promotes carcinogen-induced lung tumorigenesis. Sci Rep 11, 2097. 10.1038/s41598-021-81639-0.

26. Dederer, V., Sanz Murillo, M., Karasmanis, E.P., Hatch, K.S., Chatterjee, D., Preuss, F., Abdul Azeez, K.R., Nguyen, L.V., Galicia, C., Dreier, B., et al. (2024). A designed ankyrin-repeat protein that targets Parkinson’s disease-associated LRRK2. J Biol Chem 300, 107469. 10.1016/j.jbc.2024.107469.

27. Liu, Z., Bryant, N., Kumaran, R., Beilina, A., Abeliovich, A., Cookson, M.R., and West, A.B. (2018). LRRK2 phosphorylates membrane-bound Rabs and is activated by GTP-bound Rab7L1 to promote recruitment to the trans-Golgi network. Hum Mol Genet 27, 385–395. 10.1093/hmg/ddx410.

28. Dhekne, H.S., Tonelli, F., Yeshaw, W.M., Chiang, C.Y., Limouse, C., Jaimon, E., Purlyte, E., Alessi, D.R., and Pfeffer, S.R. (2023). Genome-wide screen reveals Rab12 GTPase as a critical activator of Parkinson’s disease-linked LRRK2 kinase. Elife 12, e87098. 10.7554/eLife.87098.

29. Wang, X., Bondar, V.V., Davis, O.B., Maloney, M.T., Agam, M., Chin, M.Y., Cheuk-Nga Ho, A., Ghosh, R., Leto, D.E., Joy, D., et al. (2023). Rab12 is a regulator of LRRK2 and its activation by damaged lysosomes. Elife 12, e87255. 10.7554/eLife.87255.

30. McGrath, E., Waschbüsch, D., Baker, B.M., and Khan, A.R. (2021). LRRK2 binds to the Rab32 subfamily in a GTP-dependent manner via its armadillo domain. Small GTPases 12, 133–146. 10.1080/21541248.2019.1666623.

31. Follett, J., Deng, I.B., Sharp, R.C., Wall, S., Mamais, A., and Farrer, M.J. (2026). Peripheral inflammation mediates midbrain Lrrk2 kinase activity via Rab32 expression. Preprint at bioRxiv, https://doi.org/10.1101/2025.09.05.674552 10.1101/2025.09.05.674552.

32. Bentley-DeSousa, A., Roczniak-Ferguson, A., and Ferguson, S.M. (2025). A STING-CASM-GABARAP pathway activates LRRK2 at lysosomes. J Cell Biol 224, e202310150. 10.1083/jcb.202310150.

33. Clegg, D., Bentley-DeSousa, A., Roczniak-Ferguson, A., and Ferguson, S.M. (2025). LRRK2 integrates Rab and GABARAP interactions to sense and respond to distinct lysosomal stresses. bioRxiv, 2025.11.19.689251. 10.1101/2025.11.19.689251.

34. Watanabe, R., Buschauer, R., Bohning, J., Audagnotto, M., Lasker, K., Lu, T.W., Boassa, D., Taylor, S., and Villa, E. (2020). The In Situ Structure of Parkinson’s Disease-Linked LRRK2. Cell 182, 1508–1518.e16.

35. Snead, D.M., Matyszewski, M., Dickey, A.M., Lin, Y.X., Leschziner, A.E., and Reck-Peterson, S.L. (2022). Structural basis for Parkinson’s disease-linked LRRK2’s binding to microtubules. Nat Struct Mol Biol 29, 1196–1207. 10.1038/s41594-022-00863-y.

36. Ito, G., Okai, T., Fujino, G., Takeda, K., Ichijo, H., Katada, T., and Iwatsubo, T. (2007). GTP binding is essential to the protein kinase activity of LRRK2, a causative gene product for familial Parkinson’s disease. Biochemistry 46, 1380–1388.

37. Biosa, A., Trancikova, A., Civiero, L., Glauser, L., Bubacco, L., Greggio, E., and Moore, D.J. (2013). GTPase activity regulates kinase activity and cellular phenotypes of Parkinson’s disease-associated LRRK2. Hum Mol Genet 22, 1140–1156. 10.1093/hmg/dds522.

38. Galicia, C., Guaitoli, G., Fislage, M., Gloeckner, C.J., and Versées, W. (2024). Structural insights into the GTP-driven monomerization and activation of a bacterial LRRK2 homolog using allosteric nanobodies. Elife 13, RP94503. 10.7554/eLife.94503.

39. Giusto, E., Yacoubian, T.A., Greggio, E., and Civiero, L. (2021). Pathways to Parkinson’s disease: a spotlight on 14-3-3 proteins. NPJ Parkinsons Dis 7, 85. 10.1038/s41531-021-00230-6.

40. Manschwetus, J.T., Wallbott, M., Fachinger, A., Obergruber, C., Pautz, S., Bertinetti, D., Schmidt, S.H., and Herberg, F.W. (2020). Binding of the Human 14-3-3 Isoforms to Distinct Sites in the Leucine-Rich Repeat Kinase 2. Frontiers in neuroscience 14, 302.

41. Obsilova, V., and Obsil, T. (2020). The 14-3-3 Proteins as Important Allosteric Regulators of Protein Kinases. Int J Mol Sci 21, 8824. 10.3390/ijms21228824.

42. Obsilova, V., and Obsil, T. (2022). Structural insights into the functional roles of 14-3-3 proteins. Front Mol Biosci 9, 1016071. 10.3389/fmolb.2022.1016071.

43. Vides, E.G., Adhikari, A., Chiang, C.Y., Lis, P., Purlyte, E., Limouse, C., Shumate, J.L., Spínola-Lasso, E., Dhekne, H.S., Alessi, D.R., et al. (2022). A feed-forward pathway drives LRRK2 kinase membrane recruitment and activation. Elife 11, e79771. 10.7554/eLife.79771.

44. Adhikari, A., Tripathi, A., Chiang, C.Y., Sherpa, P., and Pfeffer, S.R. (2025). Allosteric regulation of the Golgi-localized PPM1H phosphatase by Rab GTPases modulates LRRK2 substrate dephosphorylation in Parkinson’s disease. J Biol Chem 301, 110679. 10.1016/j.jbc.2025.110679.

45. Waschbüsch, D., Berndsen, K., Lis, P., Knebel, A., Lam, Y.P., Alessi, D.R., and Khan, A.R. (2021). Structural basis for the specificity of PPM1H phosphatase for Rab GTPases. EMBO Rep 22, e52675. 10.15252/embr.202152675.

46. Berndsen, K., Lis, P., Yeshaw, W.M., Wawro, P.S., Nirujogi, R.S., Wightman, M., Macartney, T., Dorward, M., Knebel, A., Tonelli, F., et al. (2019). PPM1H phosphatase counteracts LRRK2 signaling by selectively dephosphorylating Rab proteins. Elife 8, e50416. 10.7554/eLife.50416.

47. Punjani, A., Rubinstein, J.L., Fleet, D.J., and Brubaker, M.A. (2017). cryoSPARC: algorithms for rapid unsupervised cryo-EM structure determination. Nature Methods.

48. Bepler, T., Morin, A., Rapp, M., Brasch, J., Shapiro, L., Noble, A.J., and Berger, B. (2019). Positive-unlabeled convolutional neural networks for particle picking in cryo-electron micrographs. Nat Methods 16, 1153–1160. 10.1038/s41592-019-0575-8.

49. Pettersen, E.F., Goddard, T.D., Huang, C.C., Meng, E.C., Couch, G.S., Croll, T.I., Morris, J.H., and Ferrin, T.E. (2021). UCSF ChimeraX: Structure visualization for researchers, educators, and developers. Protein Sci 30, 70–82. 10.1002/pro.3943.

50. Abramson, J., Adler, J., Dunger, J., Evans, R., Green, T., Pritzel, A., Ronneberger, O., Willmore, L., Ballard, A.J., Bambrick, J., et al. (2024). Accurate structure prediction of biomolecular interactions with AlphaFold 3. Nature 630, 493–500. 10.1038/s41586-024-07487-w.

51. Emsley, P., and Cowtan, K. (2004). Coot: model-building tools for molecular graphics. Acta Crystallographica Section D: Biological Crystallography 60, 2126–2132.

52. He, J., Li, T., and Huang, S.-Y. (2023). Improvement of cryo-EM maps by simultaneous local and non-local deep learning. Nat Commun 14, 3217. 10.1038/s41467-023-39031-1.

53. Blaise, A.M., Corcoran, E.E., Wattenberg, E.S., Zhang, Y.-L., Cottrell, J.R., and Koleske, A.J. (2022). In vitro fluorescence assay to measure GDP/GTP exchange of guanine nucleotide exchange factors of Rho family GTPases. Biol Methods Protoc 7, bpab024. 10.1093/biomethods/bpab024.

54. Huang, B., Hubber, A., McDonough, J.A., Roy, C.R., Scidmore, M.A., and Carlyon, J.A. (2010). The Anaplasma phagocytophilum-occupied vacuole selectively recruits Rab-GTPases that are predominantly associated with recycling endosomes. Cell Microbiol 12, 1292–1307. 10.1111/j.1462-5822.2010.01468.x.

55. Raig, N.D., Surridge, K.J., Sanz-Murillo, M., Dederer, V., Krämer, A., Schwalm, M.P., Lattal, N.M., Elson, L., Chatterjee, D., Mathea, S., et al. (2025). Type II kinase inhibitors that target Parkinson’s disease-associated LRRK2. Sci Adv 11, eadt2050. 10.1126/sciadv.adt2050.

56. Fitzgerald, D.J., Berger, P., Schaffitzel, C., Yamada, K., Richmond, T.J., and Berger, I. (2006). Protein complex expression by using multigene baculoviral vectors. Nat Methods 3, 1021–1032. 10.1038/nmeth983.

57. Schindelin, J., Arganda-Carreras, I., Frise, E., Kaynig, V., Longair, M., Pietzsch, T., Preibisch, S., Rueden, C., Saalfeld, S., Schmid, B., et al. (2012). Fiji: an open-source platform for biological-image analysis. Nat Methods 9, 676–682. 10.1038/nmeth.2019.

58. Punjani, A., Rubinstein, J.L., Fleet, D.J., and Brubaker, M.A. (2017). cryoSPARC: algorithms for rapid unsupervised cryo-EM structure determination. Nature Methods 14, 290–296. 10.1038/nmeth.4169.

59. Bepler, T., Morin, A., Rapp, M., Brasch, J., Shapiro, L., Noble, A.J., and Berger, B. (2019). Positive-unlabeled convolutional neural networks for particle picking in cryo-electron micrographs. Nat Methods 16, 1153–1160. 10.1038/s41592-019-0575-8.

60. Liebschner, D., Afonine, P.V., Baker, M.L., Bunkóczi, G., Chen, V.B., Croll, T.I., Hintze, B., Hung, L.W., Jain, S., McCoy, A.J., et al. (2019). Macromolecular structure determination using X-rays, neutrons and electrons: recent developments in Phenix. Acta Crystallogr D Struct Biol 75, 861–877. 10.1107/S1959798319011471.

61. Abramson, J., Adler, J., Dunger, J., Evans, R., Green, T., Pritzel, A., Ronneberger, O., Willmore, L., Ballard, A.J., Bambrick, J., et al. (2024). Accurate structure prediction of biomolecular interactions with AlphaFold 3. Nature 630, 493–500. 10.1038/s41586-024-07487-w.

