## Supplementary Material for "The structural basis for LRRK2’s activation and autoinhibition"

### Table S1 – Cryo-EM data collection, refinement and validation statistics

| | LRRK2 autoinhibited bound to G10 DARPin (EMDB-70981) (PDB 9OXH) | LRRK2 autoinhibited bound to G10 DARPin map focused (top) (EMDB-73497) | LRRK2 autoinhibited bound to G10 DARPin map focused (WD40-DARPin) (EMDB-73499) | LRRK2 autoinhibited (EMDB-70604) (PDB 9OM2) |
|---|---|---|---|---|
| **Data collection and processing** | | | | |
| Magnification | 130000 | 13000 | 130000 | 130000 |
| Voltage (kV) | 300 | 300 | 300 | 300 |
| Electron exposure (e–/Å$^2$) | 50 | 50 | 56 | 55 |
| Defocus range (μm) | -1.0 to -3.0 | -1.0 to -3.0 | -1.0 to -3.0 | -1.0 to -3.0 |
| Pixel size (Å) | 0.935 | 0.935 | 0.935 | 0.935 |
| Symmetry imposed | C1 | C1 | C1 | C1 |
| Initial particle images (no.) | 2889148 | 2889148 | 2889148 | 2889148 |
| Final particle images (no.) | 122184 | 122184 | 122184 | 206812 |
| Map resolution (Å) | 3.15 | 3.00 | 3.21 | 3.31 |
| FSC threshold | 0.143 | 0.143 | 0.143 | 0.143 |
| Map resolution range (Å) | 2.5-4.5 | 2.5-3.5 | 2.8-5 | 3-5 |
| **Refinement** | | | | |
| Initial model used (PDB code) | 7LHW | --- | --- | 9OXH |
| Model resolution (Å) | --- | --- | --- | --- |
| FSC threshold | --- | --- | --- | --- |
| Model resolution range (Å) | --- | --- | --- | --- |
| Map sharpening *B* factor (Å$^2$) | --- | --- | --- | --- |
| Model composition | | | | |
| Non-hydrogen atoms | 14799 | --- | --- | 13980 |
| Protein residues | 1912 | --- | --- | 1782 |
| Ligands | 1 | --- | --- | 0 |
| *B* factors (Å$^2$) | | | | |
| Protein | 49.32 | --- | --- | 80.09 |
| Nucleotide | 41.31 | --- | --- | 58.07 |
| Ligand | 33.05 | --- | --- | --- |
| R.m.s. deviations | | | | |
| Bond lengths (Å) | 0.004 | --- | --- | 0.003 |
| Bond angles (°) | 0.933 | --- | --- | 0.660 |
| Validation | | | | |
| MolProbity score | 1.98 | --- | --- | 2.16 |
| Clashscore | 7.07 | --- | --- | 12.71 |
| Poor rotamers (%) | 0.06 | --- | --- | 0.64 |
| Ramachandran plot | | | | |
| Favored (%) | 88.35 | --- | --- | 90.23 |
| Allowed (%) | 10.96 | --- | --- | 9.60 |
| Disallowed (%) | 0.69 | --- | --- | 0.17 |

| | LRRK2 GMP-PNP bound active state (EMDB-72756) (PDB 9Y7I) | LRRK2 GMP-PNP bound active state focused map ROC-COR (EMDB-73485) | LRRK2 GMP-PNP bound active state focused map ROC-COR resolveEM (EMDB-73485) |
|---|---|---|---|
| **Data collection and processing** | | | |
| Magnification | 13000 | 13000 | 13000 |
| Voltage (kV) | 300 | 300 | 300 |
| Electron exposure (e–/Å$^2$) | 56 | 56 | 56 |
| Defocus range (μm) | -1.0 to -3.0 | -1.0 to -3.0 | -1.0 to -3.0 |
| Pixel size (Å) | 0.935 | 0.935 | 0.935 |
| Symmetry imposed | C1 | C1 | C1 |
| Initial particle images (no.) | 273568 | 273568 | 273568 |
| Final particle images (no.) | 94877 | 94877 | 94877 |
| Map resolution (Å) | 3.73 | 3.73 | 3.73 |
| FSC threshold | 0.143 | 0.143 | 0.143 |
| Map resolution range (Å) | 3-5 | 3-5 | 3-5 |
| **Refinement** | | | |
| Initial model used (PDB code) | 9OXH, 8TXZ | --- | --- |
| Model resolution (Å) | --- | --- | --- |
| FSC threshold | --- | --- | --- |
| Model resolution range (Å) | --- | --- | --- |
| Map sharpening *B* factor (Å$^2$) | --- | --- | --- |
| Model composition | | | |
| Non-hydrogen atoms | 8192 | --- | --- |
| Protein residues | 1101 | --- | --- |
| Ligands | 2 | --- | --- |
| *B* factors (Å$^2$) | | | |
| Protein | 139.15 | --- | --- |
| Nucleotide | --- | --- | --- |
| Ligand | 142.72 | --- | --- |
| R.m.s. deviations | | | |
| Bond lengths (Å) | 0.010 | --- | --- |
| Bond angles (°) | 1.187 | --- | --- |
| Validation | | | |
| MolProbity score | 2.16 | --- | --- |
| Clashscore | 7.93 | --- | --- |
| Poor rotamers (%) | 0.38 | --- | --- |
| Ramachandran plot | | | |
| Favored (%) | 90.63 | --- | --- |
| Allowed (%) | 9.09 | --- | --- |
| Disallowed (%) | 0.28 | --- | --- |

| | LRRK2 GMP-PNP bound partial active state (EMDB-72649) (PDB 9Y7A) | LRRK2 GMP-PNP bound partial active state Refinement (EMDB-73487) | LRRK2 GMP-PNP bound partial active state focus map ROC-COR (EMDB-73491) | LRRK2 GMP-PNP bound partial active state focus map ROC-COR resolveEM (EMDB-73492) |
|---|---|---|---|---|
| **Data collection and processing** | | | | |
| Magnification | 130000 | 130000 | 130000 | 130000 |
| Voltage (kV) | 300 | 300 | 300 | 300 |
| Electron exposure (e–/Å$^2$) | 56 | 56 | 56 | 56 |
| Defocus range (μm) | -1.0 to -3.0 | -1.0 to -3.0 | -1.0 to -3.0 | -1.0 to -3.0 |
| Pixel size (Å) | 0.935 | 0.935 | 0.935 | 0.935 |
| Symmetry imposed | C1 | C1 | C1 | C1 |
| Initial particle images (no.) | 273568 | 273568 | 273568 | 273568 |
| Final particle images (no.) | 56741 | 56741 | 56741 | 56741 |
| Map resolution (Å) | 4.40 | 4.40 | 4.40 | 4.40 |
| FSC threshold | 0.143 | 0.143 | 0.143 | 0.143 |
| Map resolution range (Å) | 4-7 | 4-7 | 3-7 | 3-7 |
| **Refinement** | | | | |
| Initial model used (PDB code) | 9OXH | --- | --- | --- |
| Model resolution (Å) | --- | --- | --- | --- |
| FSC threshold | --- | --- | --- | --- |
| Model resolution range (Å) | --- | --- | --- | --- |
| Map sharpening *B* factor (Å$^2$) | --- | --- | --- | --- |
| Model composition | | | | |
| Non-hydrogen atoms | 6384 | --- | --- | --- |
| Protein residues | 1028 | --- | --- | --- |
| Ligands | 1 | --- | --- | --- |
| *B* factors (Å$^2$) | | | | |
| Protein | 184.06 | --- | --- | --- |
| Nucleotide | --- | --- | --- | --- |
| Ligand | 170.47 | --- | --- | --- |
| R.m.s. deviations | | | | |
| Bond lengths (Å) | 0.004 | --- | --- | --- |
| Bond angles (°) | 0.880 | --- | --- | --- |
| Validation | | | | |
| MolProbity score | 1.99 | --- | --- | --- |
| Clashscore | 7.38 | --- | --- | --- |
| Poor rotamers (%) | 0 | --- | --- | --- |
| Ramachandran plot | | | | |
| Favored (%) | 82.26 | --- | --- | --- |
| Allowed (%) | 16.83 | --- | --- | --- |
| Disallowed (%) | 0.91 | --- | --- | --- |

| | LRRK2 active state (R1441C) (EMDB-72768) (PDB 9YCD) | LRRK2 active state (R1441C) Focused map ROC-COR (EMDB-73051) | LRRK2 active state (R1441C) Focused map ROC-COR (EMDB-73051) | LRRK2 autoinhibited flex-ARM (EMDB-70982) (PDB 9OXI) |
|---|---|---|---|---|
| **Data collection and processing** | | | | |
| Magnification | 130000 | 130000 | 130000 | 130000 |
| Voltage (kV) | 300 | 300 | 300 | 300 |
| Electron exposure (e–/Å$^2$) | 50 | 50 | 50 | 55 |
| Defocus range (μm) | -1.0 to -3.0 | -1.0 to -3.0 | -1.0 to -3.0 | -1.0 to -3.0 |
| Pixel size (Å) | 0.935 | 0.935 | 0.935 | 0.935 |
| Symmetry imposed | C1 | C1 | C1 | C1 |
| Initial particle images (no.) | 286935 | 286935 | 286935 | 2889148 |
| Final particle images (no.) | 63806 | 63806 | 63806 | 144198 |
| Map resolution (Å) | 3.63 | 3.63 | 3.63 | 3.72 |
| FSC threshold | 0.143 | 0.143 | 0.143 | 0.143 |
| Map resolution range (Å) | 3-9 | 3-8 | 3- | 3-6 |
| **Refinement** | | | | |
| Initial model used (PDB code) | 9OXH, 8TXZ | --- | --- | 9OXH |
| Model resolution (Å) | --- | --- | --- | --- |
| FSC threshold | --- | --- | --- | --- |
| Model resolution range (Å) | --- | --- | --- | --- |
| Map sharpening *B* factor (Å$^2$) | --- | --- | --- | --- |
| Model composition | | | | |
| Non-hydrogen atoms | 8602 | --- | --- | 12100 |
| Protein residues | 1117 | --- | --- | 1531 |
| Ligands | 1 | --- | --- | 0 |
| *B* factors (Å$^2$) | | | | |
| Protein | 140.67 | --- | --- | 92.50 |
| Nucleotide | --- | --- | --- | 47.60 |
| Ligand | 56.29 | --- | --- | --- |
| R.m.s. deviations | | | | |
| Bond lengths (Å) | 0.004 | --- | --- | 0.003 |
| Bond angles (°) | 0.978 | --- | --- | 0.812 |
| Validation | | | | |
| MolProbity score | 2.14 | --- | --- | 1.92 |
| Clashscore | 10.52 | --- | --- | 7.09 |
| Poor rotamers (%) | 0.11 | --- | --- | 0 |
| Ramachandran plot | | | | |
| Favored (%) | 88.41 | --- | --- | 90.74 |
| Allowed (%) | 11.22 | --- | --- | 8.99 |
| Disallowed (%) | 0.37 | --- | --- | 0.27 |

| | LRRK2 after symmetry expansion (G2019S) (EMDB-72556) (PDB 9Y68) | LRRK2 after symmetry expansion (R1441C) (EMDB-71012) (PDB 9OYA) | LRRK2 asymmetry dimer (EMDB-72751) (PDB 9YBL) | LRRK2 after symmetry expansion (EMDB-72555) (PDB 9Y67) |
|---|---|---|---|---|
| **Data collection and processing** | | | | |
| Magnification | 130000 | 130000 | 130000 | 13000 |
| Voltage (kV) | 300 | 300 | 300 | 300 |
| Electron exposure (e–/Å$^2$) | 55 | 55 | 55 | 55 |
| Defocus range (μm) | -1.0 to -3.0 | -1.0 to -3.0 | -1.0 to -3.0 | -1.0 to -3.0 |
| Pixel size (Å) | 0.935 | 0.935 | 0.935 | 0.935 |
| Symmetry imposed | C1 | C1 | C1 | C1 |
| Initial particle images (no.) | 2889148 | 2889148 | 2889148 | 2889148 |
| Final particle images (no.) | 83730 | 98448 | 79889 | 41214 |
| Map resolution (Å) | 3.25 | 3.44 | 3.44 | 3.72 |
| FSC threshold | 0.143 | 0.143 | 0.143 | 0.143 |
| Map resolution range (Å) | 3-5 | 3-5 | 3-6 | 3-6 |
| **Refinement** | | | | |
| Initial model used (PDB code) | 9OXH | 9OXH | 9OXH | 9OXH |
| Model resolution (Å) | --- | --- | --- | --- |
| FSC threshold | --- | --- | --- | --- |
| Model resolution range (Å) | --- | --- | --- | --- |
| Map sharpening *B* factor (Å$^2$) | --- | --- | --- | --- |
| Model composition | | | | |
| Non-hydrogen atoms | 13046 | 14270 | 15523 | 13173 |
| Protein residues | 1700 | 1801 | 2071 | 1713 |
| Ligands | 0 | 0 | 0 | 0 |
| *B* factors (Å$^2$) | | | | |
| Protein | 109.89 | 102.36 | 117.93 | 126.71 |
| Nucleotide | 76.92 | 75.05 | 38.41 | 85.35 |
| Ligand | --- | --- | -- | --- |
| R.m.s. deviations | | | | |
| Bond lengths (Å) | 0.003 | 0.004 | 0.004 | 0.004 |
| Bond angles (°) | 0.660 | 0.809 | 0.743 | 0.946 |
| Validation | | | | |
| MolProbity score | 1.93 | 1.92 | 1.98 | 2.04 |
| Clashscore | 7.17 | 6.74 | 8.25 | 8.77 |
| Poor rotamers (%) | 0 | 0 | 0.06 | 0.14 |
| Ramachandran plot | | | | |
| Favored (%) | 90.56 | 90.15 | 88.25 | 89.07 |
| Allowed (%) | 9.38 | 9.79 | 11.35 | 10.51 |
| Disallowed (%) | 0.06 | 0.06 | 0.40 | 0.42 |

| | LRRK2 active state 1 (APO ROC) EMDB-72762) (PDB 9YC8) | LRRK2 active state 1 (APO ROC) focus map ROC-COR EMDB-73496) | LRRK2 active state 2 (APO ROC) EMDB-72756) (PDB 9YBP) | LRRK2 active state 2 (APO ROC) EMDB-72756) (PDB 9YBP) |
|---|---|---|---|---|
| Data collection and processing | | | | |
| Magnification | 130000 | 130000 | 13000 | 13000 |
| Voltage (kV) | 300 | 300 | 300 | 300 |
| Electron exposure (e–/Å2) | 55 | 55 | 55 | 55 |
| Defocus range (μm) | -1.0 to -3.0 | -1.0 to -3.0 | -1.0 to -3.0 | -1.0 to -3.0 |
| Pixel size (Å) | 0.935 | 0.935 | 0.935 | 0.935 |
| Symmetry imposed | C1 | C1 | C1 | C1 |
| Initial particle images (no.) | 2889148 | 2889148 | 2889148 | 2889148 |
| Final particle images (no.) | 76274 | 76274 | 86982 | 86982 |
| Map resolution (Å) | 4.02 | 4.01 | 4 | 4.05 |
| FSC threshold | 0.143 | 0.143 | 0.143 | 0.143 |
| Map resolution range (Å) | 3.5-7.2 | 3.5-7.4 | 3.4-8 | 3-8 |
| Refinement | | | | |
| Initial model used (PDB code) | 9OXH | --- | 9OXH | --- |
| Model resolution (Å) | --- | --- | --- | --- |
| FSC threshold | --- | --- | --- | --- |
| Model resolution range (Å) | --- | --- | --- | --- |
| Map sharpening B factor (Å2) | --- | --- | --- | --- |
| Model composition | | | | |
| Non-hydrogen atoms | 7822 | --- | 6587 | --- |
| Protein residues | 1051 | --- | 990 | --- |
| Ligands | 1 | --- | 0 | --- |
| B factors (Å2) | | | | |
| Protein | 127.40 | --- | 119.59 | --- |
| Nucleotide | 86.07 | --- | --- | --- |
| Ligand | --- | --- | --- | --- |
| R.m.s. deviations | | | | |
| Bond lengths (Å) | 0.003 | --- | 0.002 | --- |
| Bond angles (°) | 0.662 | --- | 0.669 | --- |
| Validation | | | | |
| MolProbity score | 2.20 | --- | 1.88 | --- |
| Clashscore | 6.41 | --- | 6.61 | --- |
| Poor rotamers (%) | 0.26 | --- | 0 | --- |
| Ramachandran plot | | | | |
| Favored (%) | 88.81 | --- | 93.55 | --- |
| Allowed (%) | 10.99 | --- | 6.03 | --- |
| Disallowed (%) | 0.20 | --- | 0.42 | --- |

| | LRRK2 autoinhibited (EMDB-72941) (PDB 9YGT) | LRRK2 autoinhibited (EMD-72978) | LRRK2 active state 3 (APO ROC) (EMD-72978) |
|---|---|---|---|
| **Data collection and processing** | | | |
| Magnification | 13000 | 1300 | 1300 |
| Voltage (kV) | 300 | 300 | 300 |
| Electron exposure (e–/Å$^2$) | 55 | 55 | 55 |
| Defocus range (μm) | -1.0 to -3.0 | -1.0 to -3.0 | -1.0 to -3.0 |
| Pixel size (Å) | 0.935 | 0.935 | 0.935 |
| Symmetry imposed | C1 | C1 | C1 |
| Initial particle images (no.) | 2889148 | 2889148 | 2889148 |
| Final particle images (no.) | 129152 | 50446 | 112087 |
| Map resolution (Å) | 3.42 | 4.14 | 3.72 |
| FSC threshold | 0.143 | 0.143 | 0.143 |
| Map resolution range (Å) | 3-5 | 4-6 | 3-6 |
| **Refinement** | | | |
| Initial model used (PDB code) | 9OM2 | -- | -- |
| Model resolution (Å) | --- | --- | --- |
| FSC threshold | --- | --- | --- |
| Model resolution range (Å) | --- | --- | --- |
| Map sharpening *B* factor (Å$^2$) | --- | --- | --- |
| Model composition | | | |
| Non-hydrogen atoms | 13934 | --- | --- |
| Protein residues | 1774 | --- | --- |
| Ligands | 0 | --- | --- |
| *B* factors (Å$^2$) | | | |
| Protein | 55.10 | --- | --- |
| Nucleotide | 48.81 | --- | --- |
| Ligand | --- | --- | --- |
| R.m.s. deviations | | | |
| Bond lengths (Å) | 0.005 | --- | --- |
| Bond angles (°) | 0.939 | --- | --- |
| Validation | | | |
| MolProbity score | 2.04 | --- | --- |
| Clashscore | 8.79 | --- | --- |
| Poor rotamers (%) | 0 | --- | --- |
| Ramachandran plot | | | |
| Favored (%) | 89.49 | --- | --- |
| Allowed (%) | 10.29 | --- | --- |
| Disallowed (%) | 0.23 | --- | --- |

| | LRRK2$^{RCKW}$ bound to C12 DARPin (EMDB-73338) (PDB-9YQK) | LRRK2$^{RCKW}$ map focused on C-terminal kinase domain, WD40 and C12 DARPin (EMD-73280) | LRRK2$^{RCKW}$ map focused on ROC, COR-A, COR-B and N-terminal kinase domain (EMD-73281) | LRRK2$^{RCKW}$ bound to DARPin C12 consensus map (EMD-73282) |
|---|---|---|---|---|
| **Data collection and processing** | | | | |
| Magnification | 36000 | 36000 | 36000 | 36000 |
| Voltage (kV) | 200 | 200 | 200 | 200 |
| Electron exposure (e–/Å$^2$) | 51 | 51 | 51 | 51 |
| Defocus range (μm) | -1.0 to -3.0 | -1.0 to -3.0 | -1.0 to -3.0 | -1.0 to -3.0 |
| Pixel size (Å) | 1.16 | 1.16 | 1.16 | 1.16 |
| Symmetry imposed | C1 | C1 | C1 | C1 |
| Initial particle images (no.) | 591732 | 591732 | 591732 | 591732 |
| Final particle images (no.) | 392016 | 392016 | 392016 | 392016 |
| Map resolution (Å) | | 3.48 | 3.6 | 3.72 |
| FSC threshold | | 0.143 | 0.143 | 0.143 |
| Map resolution range (Å) | 3-5 | 3-5 | 3-5 | 3-5 |
| **Refinement** | | | | |
| Initial model used (PDB code) | 6VP7 | -- | -- | -- |
| Model resolution (Å) | --- | --- | --- | --- |
| FSC threshold | --- | --- | --- | --- |
| Model resolution range (Å) | --- | --- | --- | --- |
| Map sharpening *B* factor (Å$^2$) | --- | --- | --- | --- |
| Model composition | | | | |
| Non-hydrogen atoms | 8925 | --- | --- | --- |
| Protein residues | 1123 | --- | --- | --- |
| Ligands | 0 | --- | --- | --- |
| *B* factors (Å$^2$) | | | | |
| Protein | 105.88 | --- | --- | --- |
| Nucleotide | --- | --- | --- | --- |
| Ligand | --- | --- | --- | --- |
| R.m.s. deviations | | | | |
| Bond lengths (Å) | 0.004 | --- | --- | --- |
| Bond angles (°) | 0.799 | --- | --- | --- |
| Validation | | | | |
| MolProbity score | 2.18 | --- | --- | --- |
| Clashscore | 15.40 | --- | --- | --- |
| Poor rotamers (%) | 0 | --- | --- | --- |
| Ramachandran plot | | | | |
| Favored (%) | 91.98 | --- | --- | --- |
| Allowed (%) | 8.02 | --- | --- | --- |
| Disallowed (%) | 0.00 | --- | --- | --- |

### SUPPLEMENTARY FIGURES

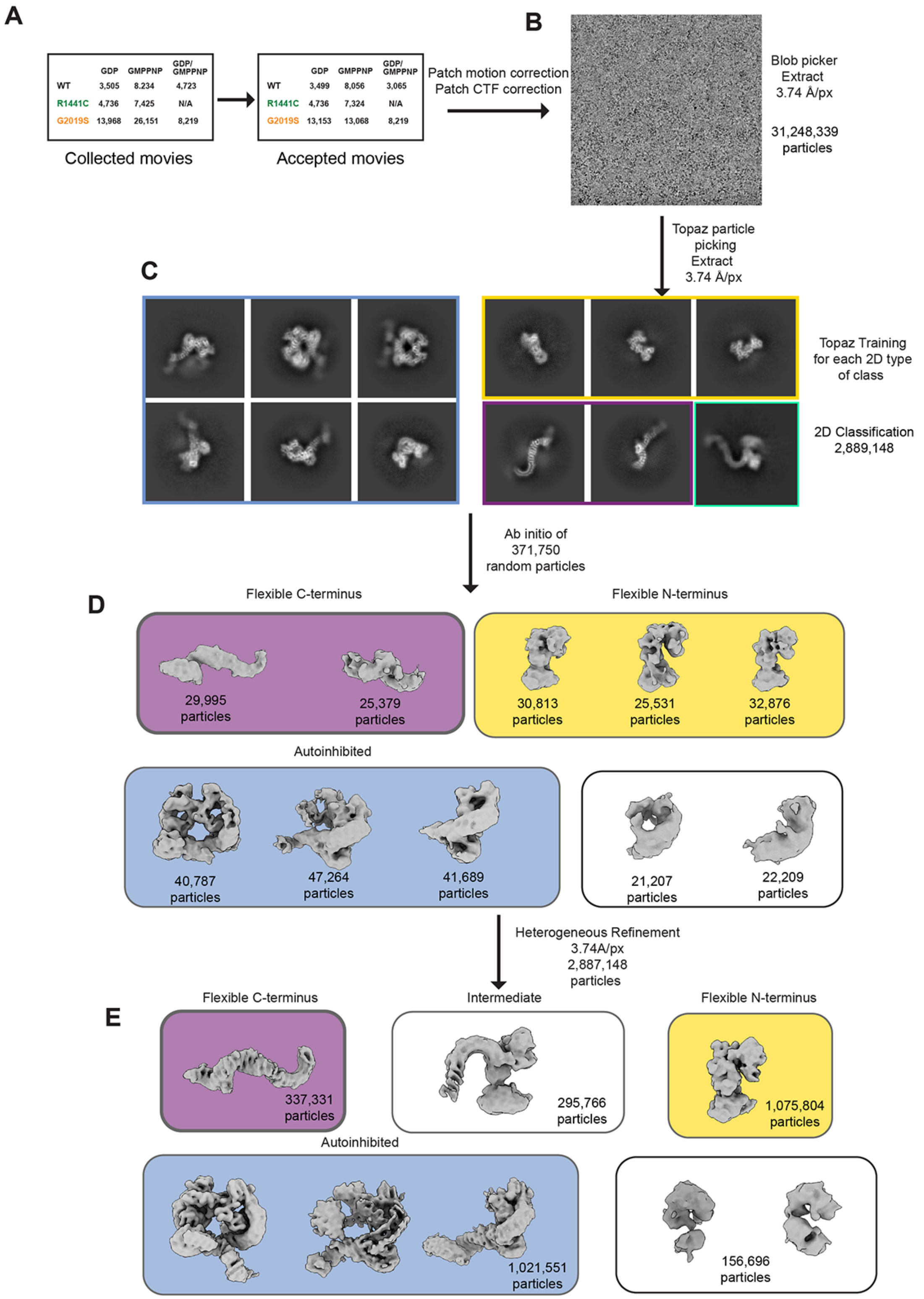

**Figure S1. Overall Cryo-EM processing pipeline for the combined data set. (A)** Number of movies that were collected and kept for further processing. **(B, C)** Representative micrograph (B) and class averages (C). **(D)** Data processing strategy. **(E)** Splitting of data into volumes representing different states/conformations of LRRK2.

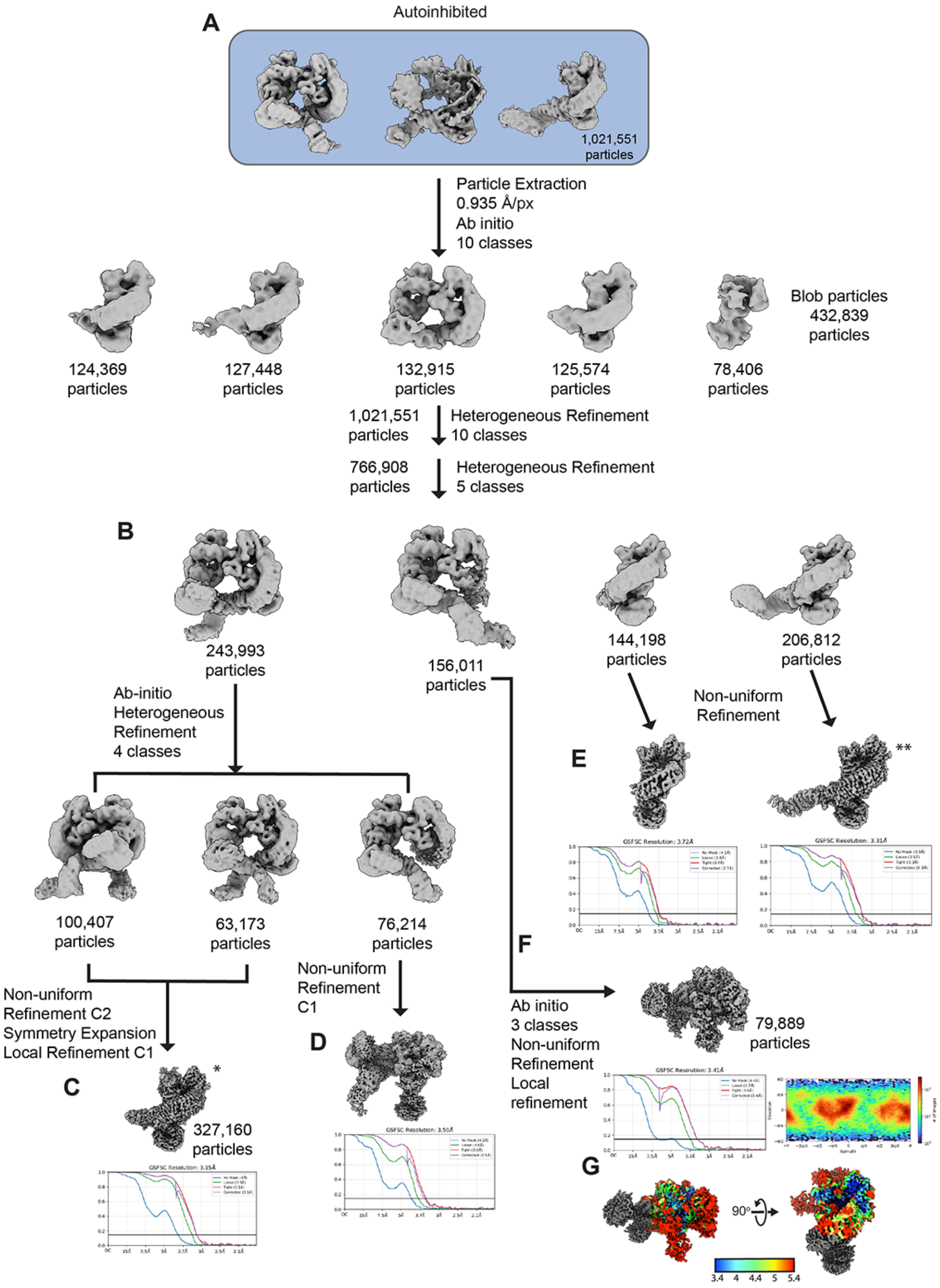

**Figure S2. Cryo-EM processing pipeline for the combined data set: autoinhibited LRRK2 and mixed dimers. (A)** Data processing strategy. **(B)** Processing of the autoinhibited dimer. **(C)** Volume and FSC curve for the dimer after symmetry expansion. **(D)** Volume and FSC curve for the asymmetric dimer (autoinhibited protomer). **(E)** Volumes and FSC curves for autoinhibited monomers. **(F)** Volume, FSC curve and Euler angle distribution for the asymmetric dimer (activated protomer). **(G)** Local resolution map. * and ** indicate volumes whose processing continues in Figure S3.

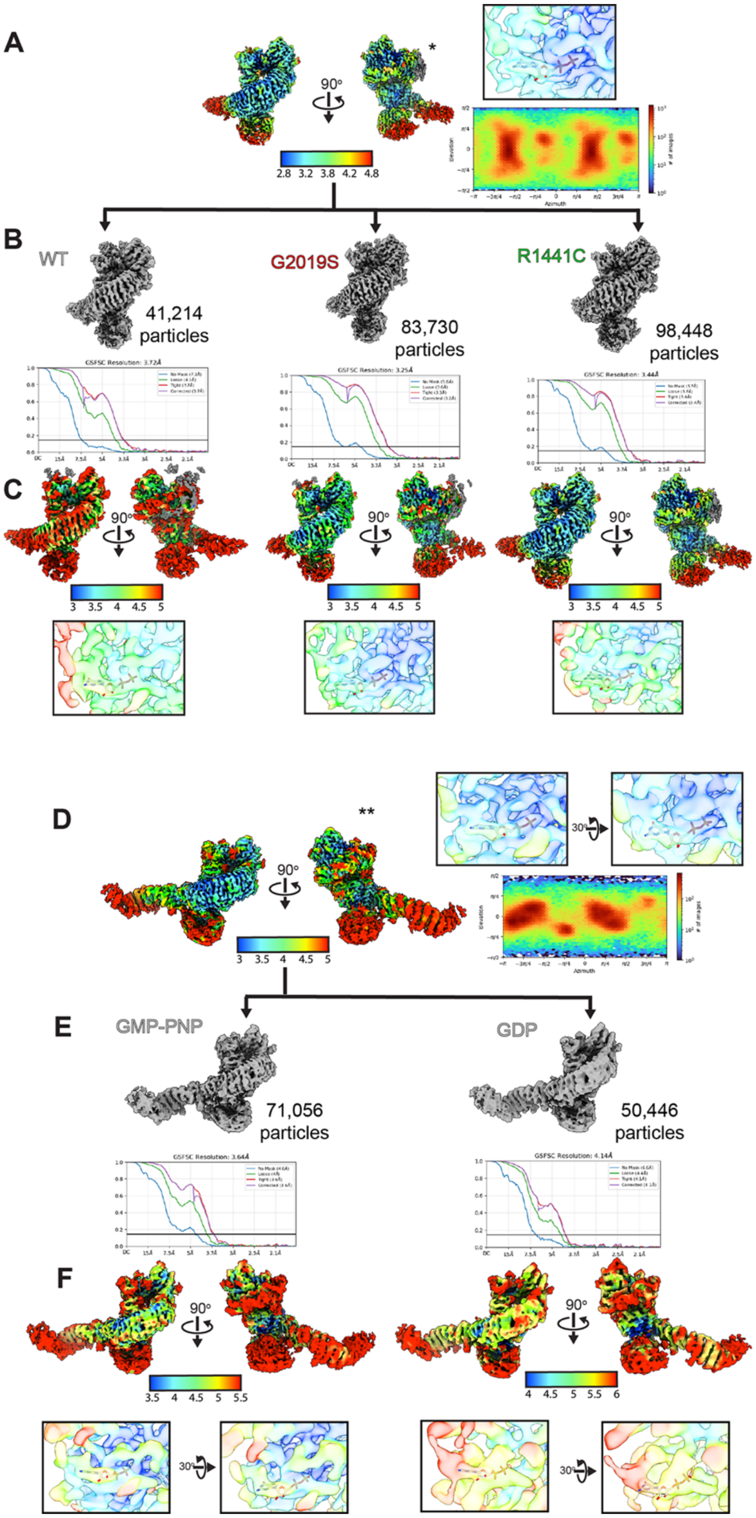

**Figure S3. Cryo-EM processing pipeline for autoinhibited LRRK2 and mixed dimers: splitting by nucleotide added or by LRRK2 variant. (A)** Local resolution map and Euler angle distribution for the dimer after symmetry expansion. **(B)** Data splitting by LRRK2 variant; volumes and FSC curves. **(C)** Local resolution map in the vicinity of the nucleotide-binding pocket of the ROC domain. **(D)** Local resolution map and Euler angle distribution for the monomer. **(E)** Data splitting by the guanine nucleotide added to the sample; volumes and FSC curves. **(F)** Local resolution map in the vicinity of the nucleotide-binding pocket of the ROC domain.

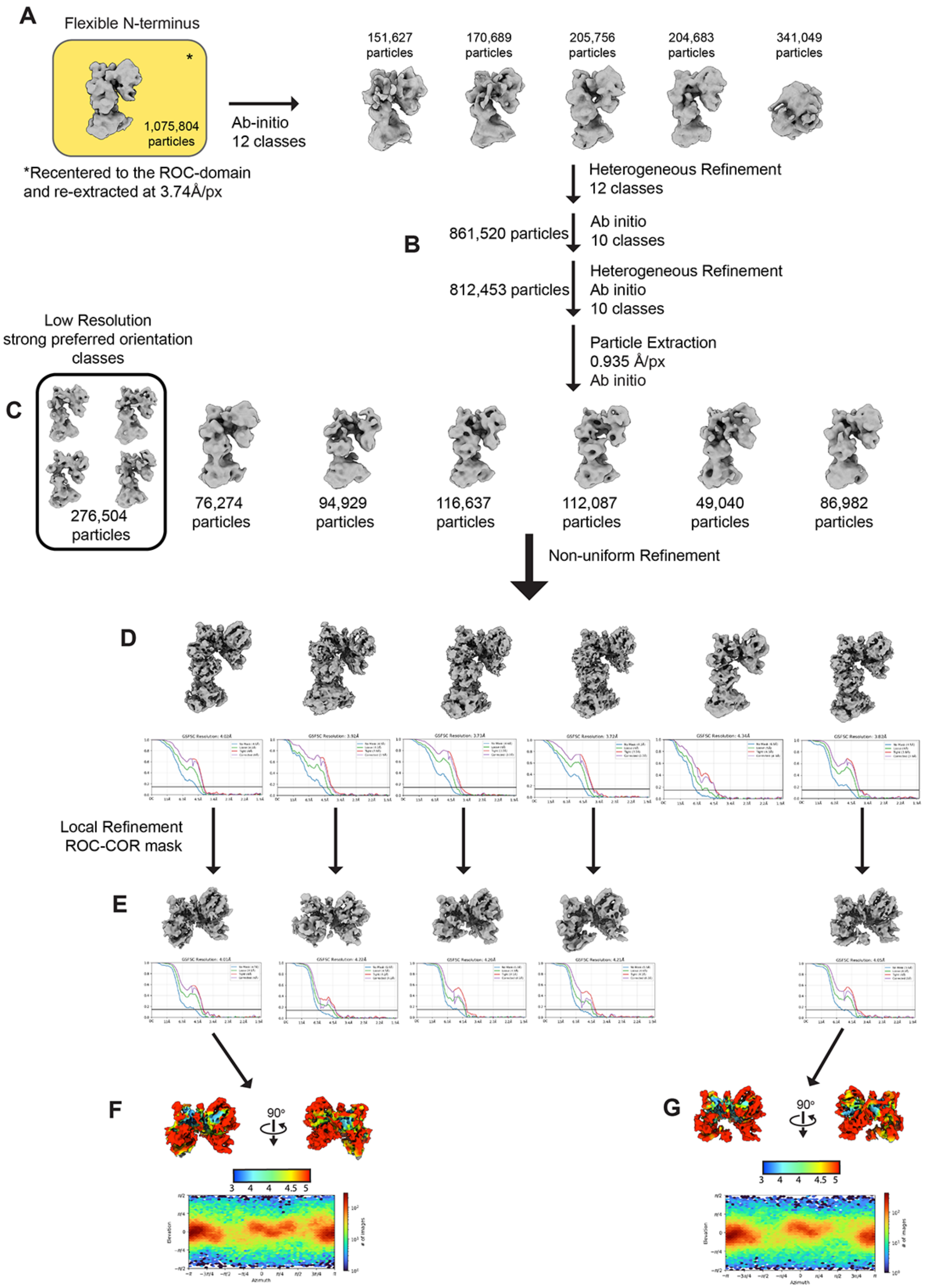

**Figure S4. Cryo-EM processing pipeline for the combined data set: activated LRRK2 (C-terminal half). (A, B)** Data processing strategy. **(C)** Splitting of activated volumes by conformation. **(D)** Volumes and FSC curves of activated LRRK2. **(E)** Local refinement volumes and FSC curves. **(F, G)** Local resolution maps and Euler angle distributions for activated LRRK2 (apo).

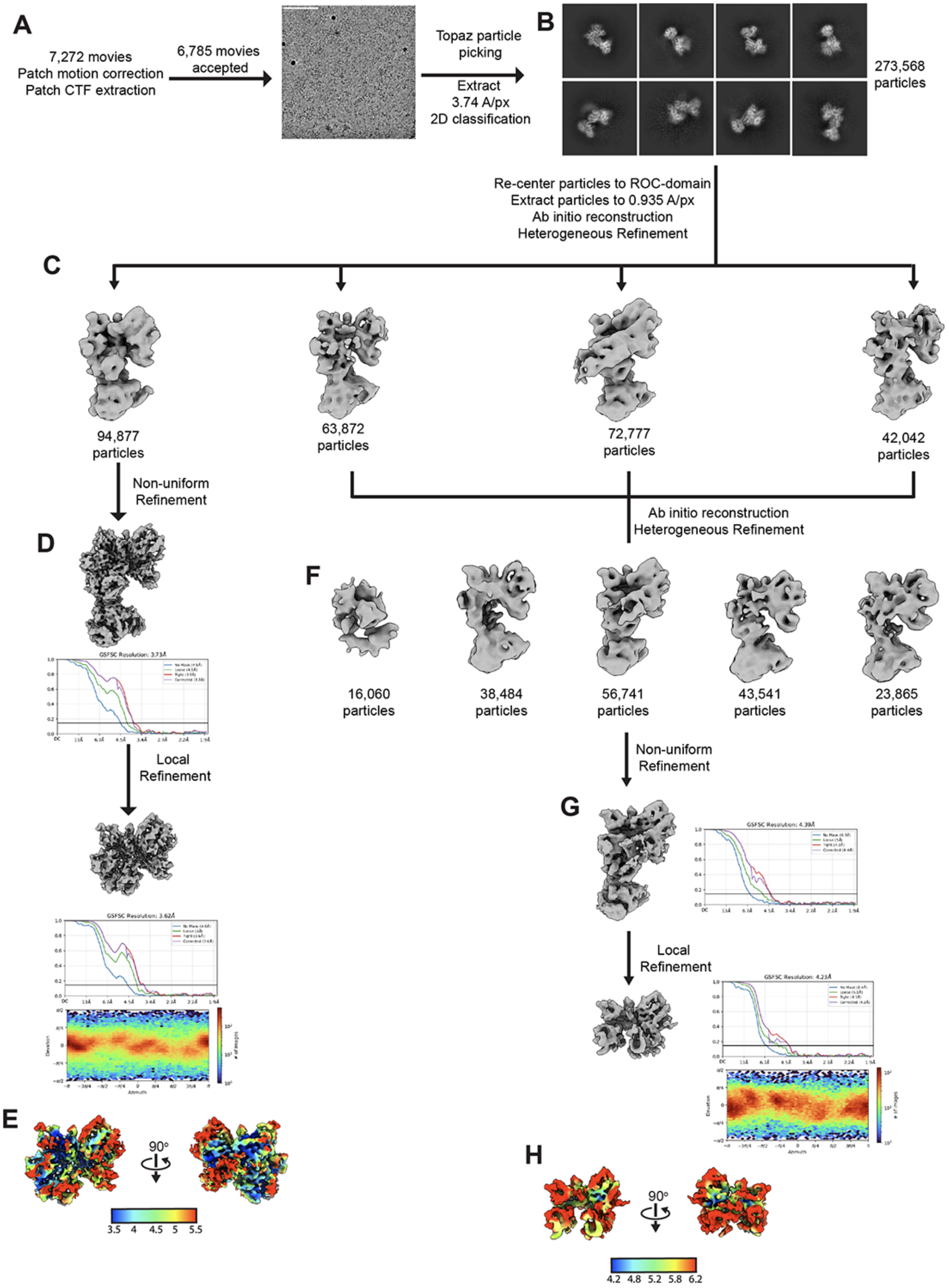

**Figure S5. Cryo-EM data processing pipeline for LRRK2 incubated with GMPPNP for 24h, active and semi-active conformations. (A)** Number of movies collected and kept for further processing, and representative micrograph. **(B)** 2D class averages. **(C, F)** Data processing strategy. **(D)** Volume, FSC curve and Euler angle distribution, and **(E)** local resolution map of the ROC–COR domain of activated LRRK2. **(G)** Volume, FSC curve and Euler angle distribution. **(H)** Local resolution map of the ROC–COR domain of 'semi-activated' LRRK2.

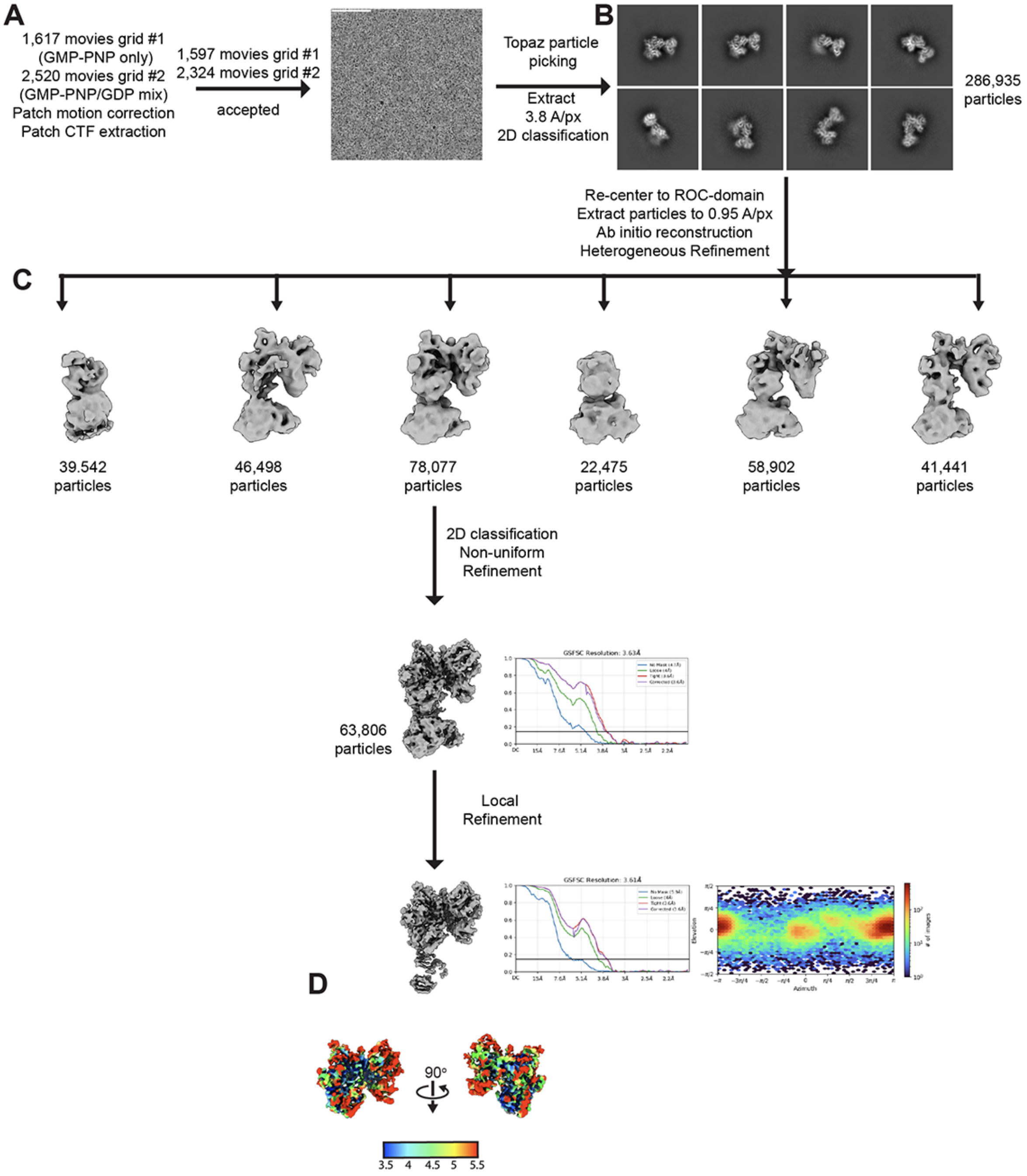

**Figure S6. Cryo-EM processing pipeline for activated LRRK2[R1441C]. (A)** Number of movies collected and kept for further processing, and representative micrograph. **(B)** 2D class averages. **(C)** Data processing strategy. **(D)** Volume, FSC curve, Euler angle distribution, and local resolution map of the ROC–COR domain of activated LRRK2[R1441C].

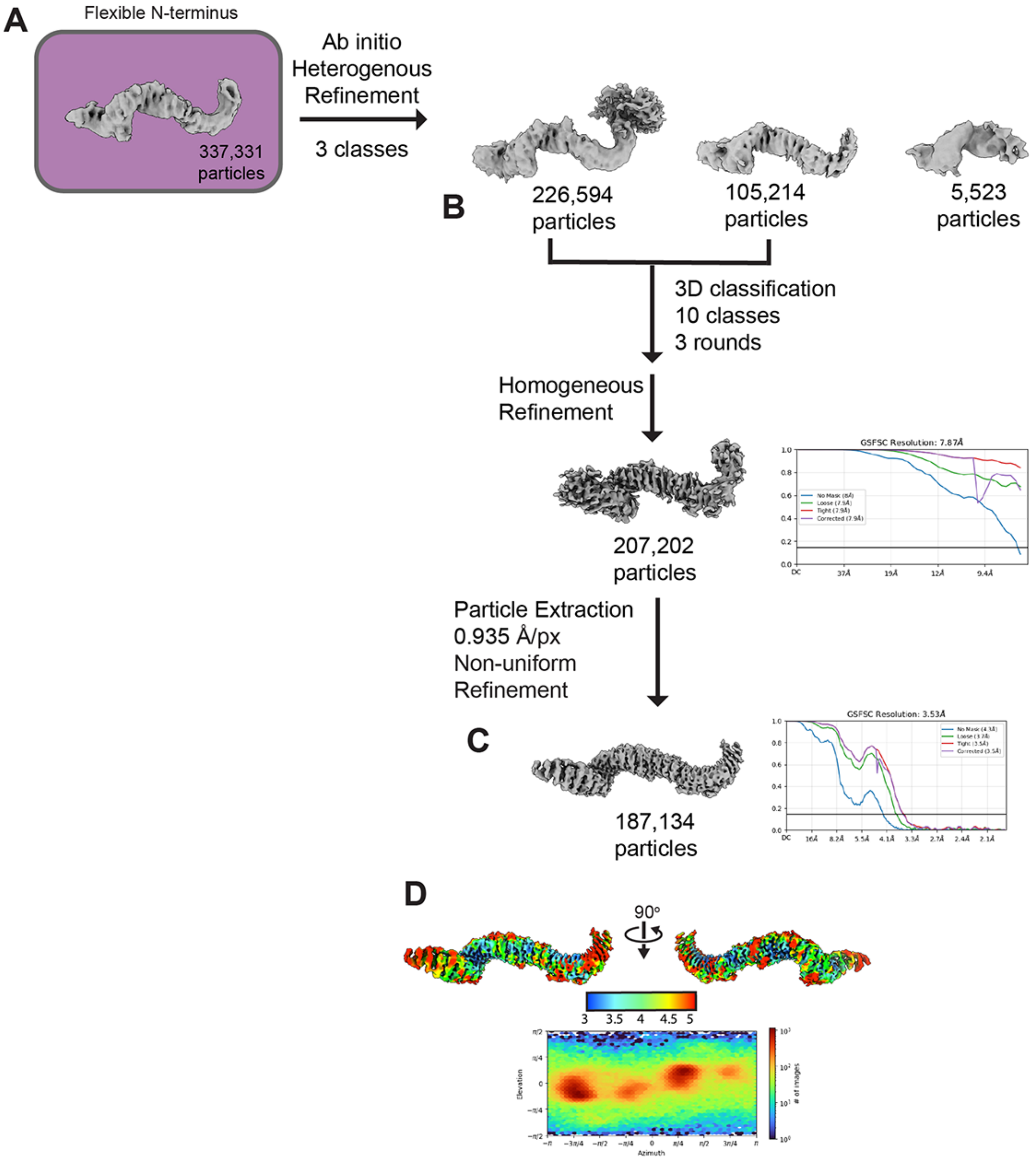

**Figure S7. Cryo-EM data processing pipeline for the N-terminal repeats. (A, B)** Data processing strategy. **(C)** Volume and FSC curve. **(D)** Local resolution map and Euler angle distribution.

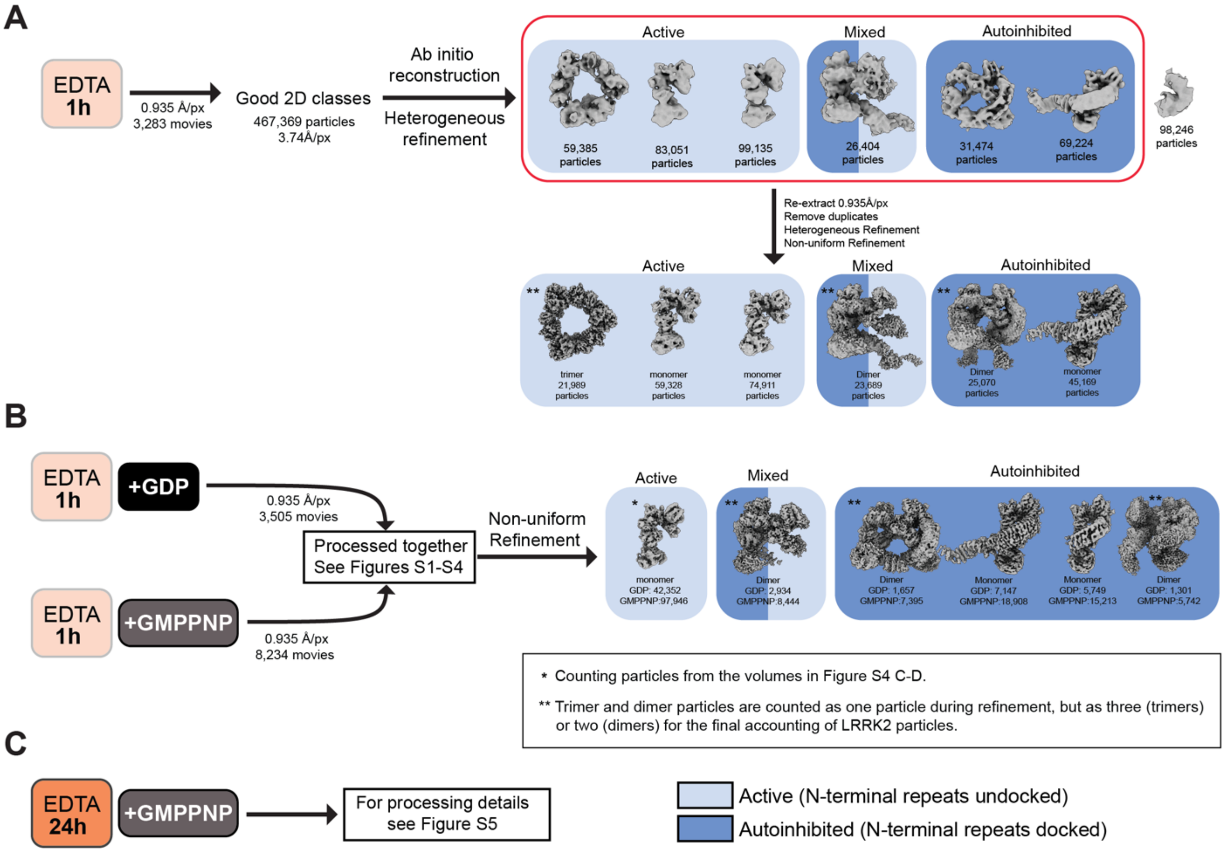

**D**

| EDTA | 1h | 1h | 1h | 24h |
|---|---|---|---|---|
| **Nucleotide** | **none** | **GDP / 1h** | **GMPPNP / 1h** | **GMPPNP / 24h** |
| Active | 223,895 | 45,291 | 230,482 | 257,508 |
| Autoinhibited | 118,998 | 21,751 | 68,839 | 0 |

| EDTA | 1h | 1h | 1h | 24h |
|---|---|---|---|---|
| **Nucleotide** | **none** | **GDP / 1h** | **GMPPNP / 1h** | **GMPPNP / 24h** |
| Active | 65% | 68% | 77% | 100% |
| Autoinhibited | 35% | 32% | 23% | 0% |

**Figure S8. Distribution of autoinhibited and activated LRRK2 in samples that were treated with EDTA for 1h vs. 24h. (A)** Cryo-EM data processing of nucleotide-stripped LRRK2 (EDTA treatment only), yielding active LRRK2 in monomeric, trimeric, and mixed dimer forms, as well as autoinhibited LRRK2 in monomeric, dimeric, and mixed dimer forms. **(B)** Data classification for LRRK2 after nucleotide stripping and 1h incubation with GDP or GMPPNP, yielding active LRRK2 monomers and mixed dimers, and autoinhibited monomers, dimers, and mixed dimers. **(C)** Data classification for LRRK2 after nucleotide stripping for 24h and incubation with GMPPNP for 24h, yielding exclusively active LRRK2 monomers (see Figure S5 for data processing details). **(D)** Number and relative abundance (%) of particles corresponding to the active and inactive conformations of LRRK2 derived from cryo-EM datasets of LRRK2 subjected to EDTA treatment for 1h or 24h and then incubated with one of the following: no nucleotide, GDP, GMPPNP. The graph in Figure 2H corresponds to the relative abundances shown in the table.

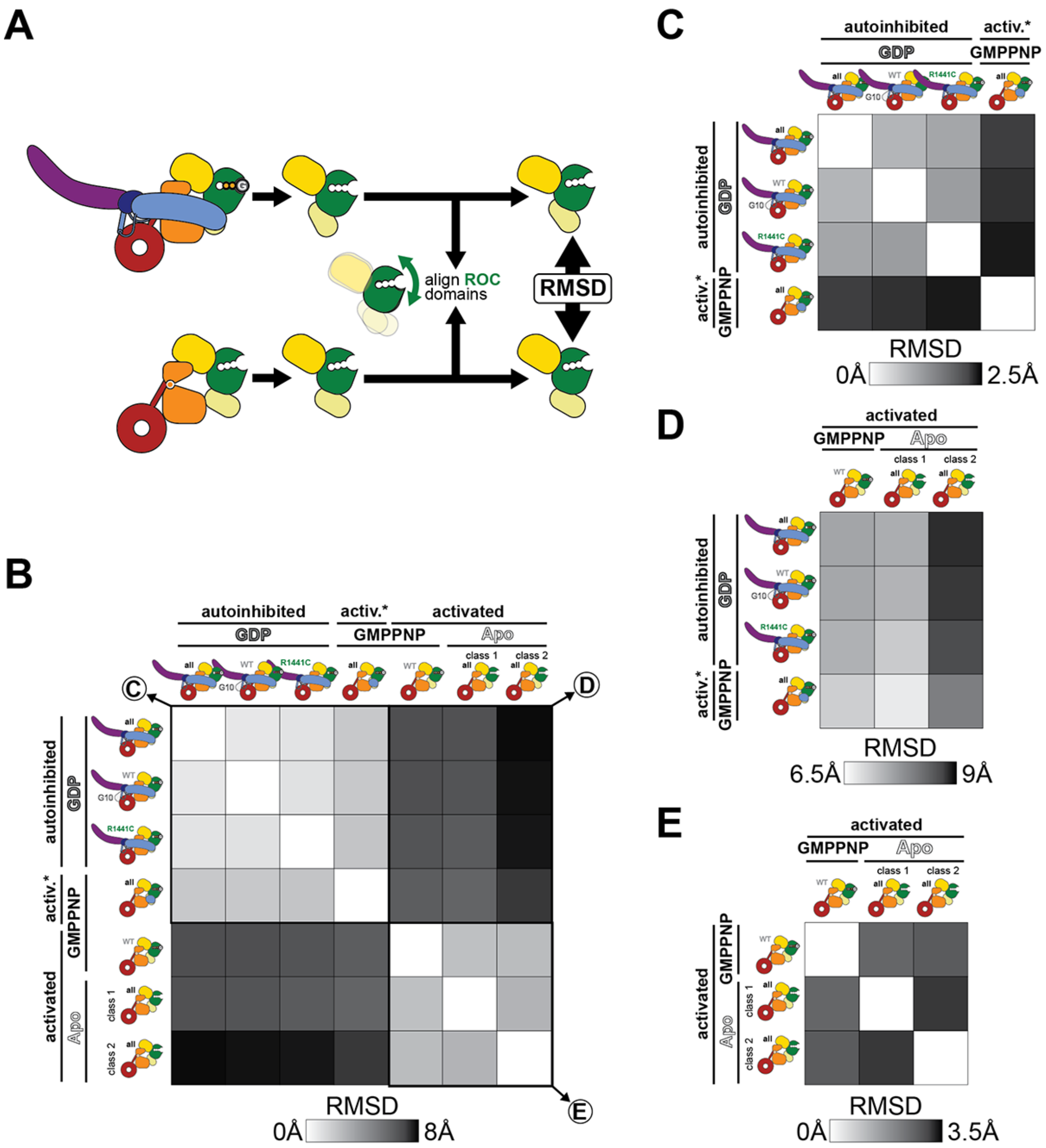

**Figure S9. Quantification of conformational differences among structures of autoinhibited and activated LRRK2. (A)** Schematic representation of the approach used here to quantify conformation differences between structures. We focused our analysis on the ROC-COR module, where the largest conformational changes take place in the C-terminal half of LRRK2 ($LRRK2^{RCKW}$). For any pair of structures—in this example an autoinhibited LRRK2:GDP and an activated LRRK2:Apo—we extracted their ROC-COR modules and aligned them to each other by their ROC domains. We then measured the Root Mean Squared Deviation (RMSD) between the two ROC-COR modules (see Methods). Those values are shown in the matrices in greyscale. **(B)** Matrix of pairwise comparisons (RMSDs) for 7 structures: 3 of autoinhibited LRRK2, 3 of activated LRRK2, and the one structure of activated LRRK2 where we saw the LRR partially docked (labeled 'activated*' here). For all structures, we indicated the LRRK2 variant and the nucleotide state of the ROC domain. The RMSD scale used in this matrix is shown at the bottom. Three quadrants, which are replotted separately with different RMSD scales, are highlighted with thicker lines and their panels indicated. **(C-E)**, The 3 quadrants highlighted in (B) are shown with RMSD scales expanded to reveal differences among each subset of structures.

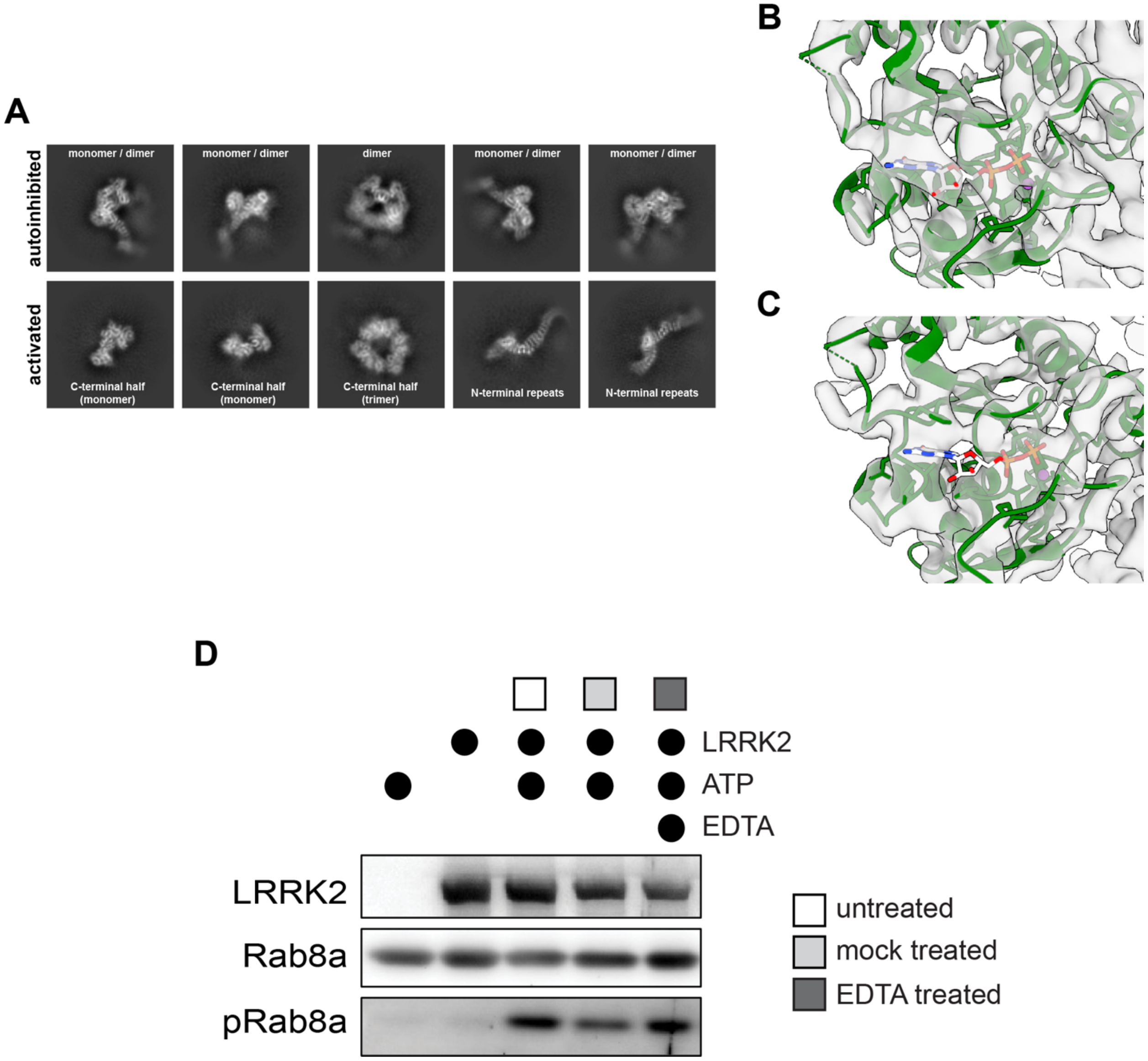

**Figure S10. The nucleotide stripping protocol activates LRRK2. (A)** 2D class averages for LRRK2 that was subjected to the nucleotide-stripping protocol but to which no additional nucleotide was added. The top and bottom rows show species of autoinhibited (top) and activated (bottom) LRRK2. **(B, C)** Close-ups of the ROC domain for reconstructions obtained from the data shown in (A) for autoinhibited (B) and activated (C) LRRK2. The density in autoinhibited LRRK2 (B) can accommodate a nucleotide (GDP was modeled here based on the data discussed in the manuscript) while activated LRRK2 (C) cannot. **(D)** Representative immunoblot for in vitro Rab8a phosphorylation assays with or without treatment with EDTA (Figure 4B). ATP was omitted in one condition as a negative control.

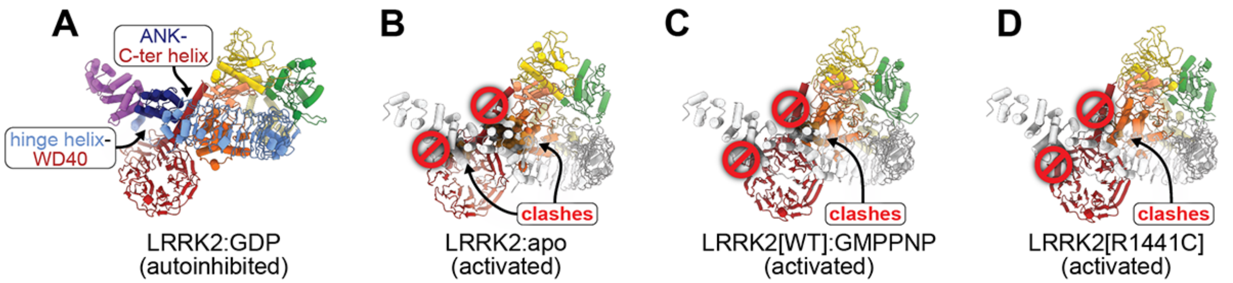

**Figure S11. Modeling the redocking of the N-terminal repeats in the context of activated LRRK2**. **(A)** Structure of GDP-bound, autoinhibited LRRK2 highlighting the two main interactions that stabilize the docked conformation: one between the hinge helix in the LRR and the WD40 domain, and the other between the ANK domain and the C-terminal helix the follows the WD40. **(B-D)**, We modeled the N-terminal repeats into three of our structures of activated LRRK2: LRRK2:Apo (class 2) (Figure 3E), LRRK2[WT]:GMPPNP (Figure 3B), and LRRK2[R1441C] (Figure 3F). In all cases, we took a fragment comprising the N-terminal repeats and the ROC domain from autoinhibited LRRK2 and aligned that to each of the structures of activated LRRK2 via their ROC domains. These panels show that the interactions shown in (A) cannot form when the C-terminal half of LRRK2 is in its active conformation without additional flexibility in the N-terminal repeats, which we did not observe in our data. Furthermore, there are clashes between the repeats and the C-terminal half in all three models.

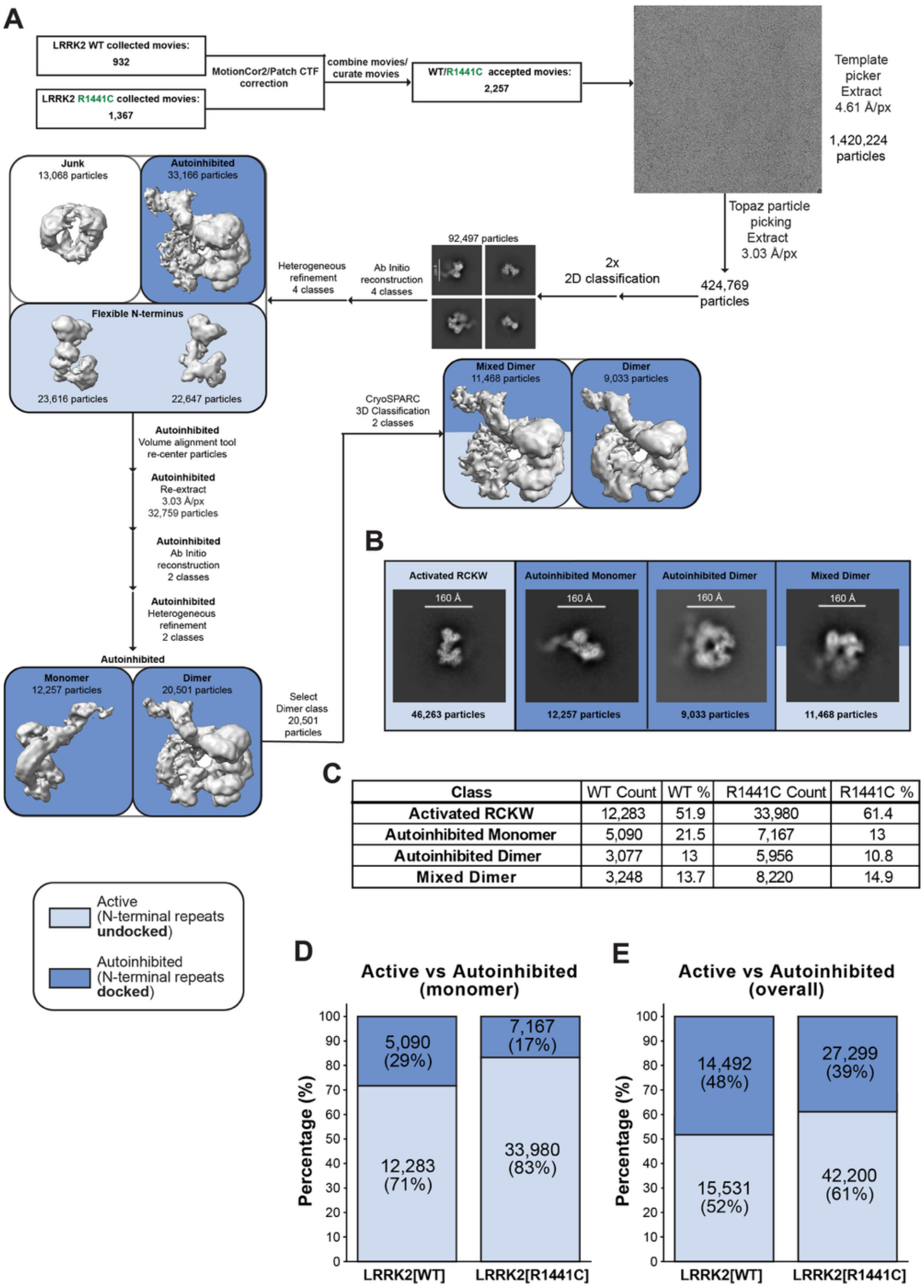

| Class | WT Count | WT % | R1441C Count | R1441C % |
|---|---|---|---|---|
| Activated RCKW | 12,283 | 51.9 | 33,980 | 61.4 |
| Autoinhibited Monomer | 5,090 | 21.5 | 7,167 | 13 |
| Autoinhibited Dimer | 3,077 | 13 | 5,956 | 10.8 |
| Mixed Dimer | 3,248 | 13.7 | 8,220 | 14.9 |

**Figure S12**. **The mutation R1441C increases the fraction of LRRK2 in the active conformation (N-terminal repeats undocked). (A)** Data processing workflow for the combined LRRK2[WT] and LRRK2[R1441C] datasets. Light blue denotes active states and dark blue denotes autoinhibited states. **(B)** Representative 2D class averages for Activated LRRK2 ("RCKW"), Autoinhibited Monomer, Autoinhibited Dimer, and Mixed Dimer states derived from the combined dataset (79,021 total particles). **(C)** Particle counts and distributions across structural classes for LRRK2[WT] and LRRK2[R1441C] datasets. **(D)** Partition of active and autoinhibited states for monomeric LRRK2[WT] and LRRK2[R1441C]. **(E)** Partition of active and autoinhibited states for all species of LRRK2[WT] and LRRK2[R1441C]. Mixed dimers are counted twice, contributing one protomer to the active counts and one protomer to the autoinhibited counts.

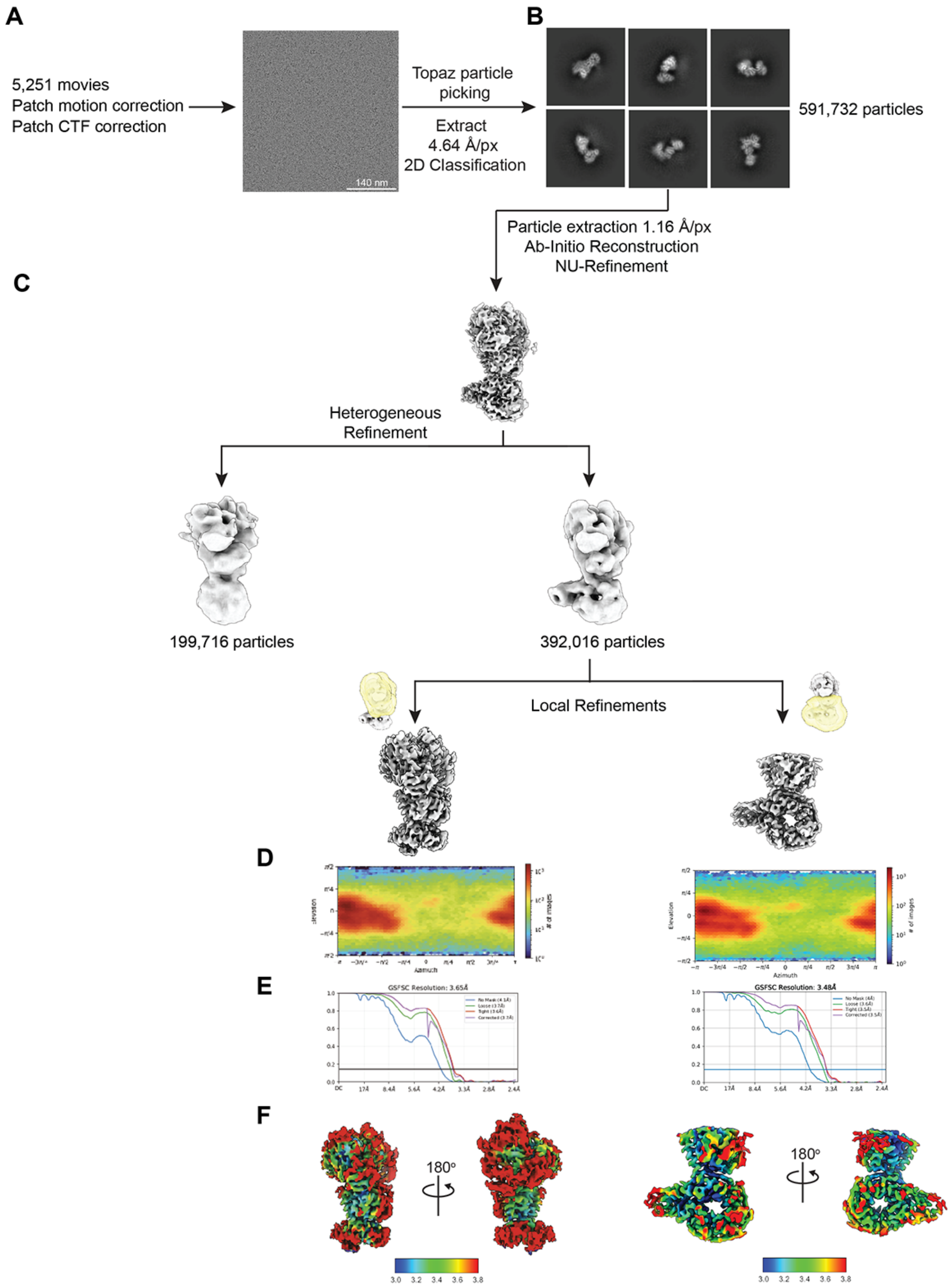

**Figure S13. Cryo-EM data processing pipeline for LRRK2^RCKW bound to DARPin C12. (A)** Number of movies collected and kept for further processing, and representative micrograph. **(B)** 2D class averages. **(C)** Data processing strategy. **(D, E)** Euler angle distribution (D) and FSC curve (E) for the volumes shown above. **(F)** Local resolution map.

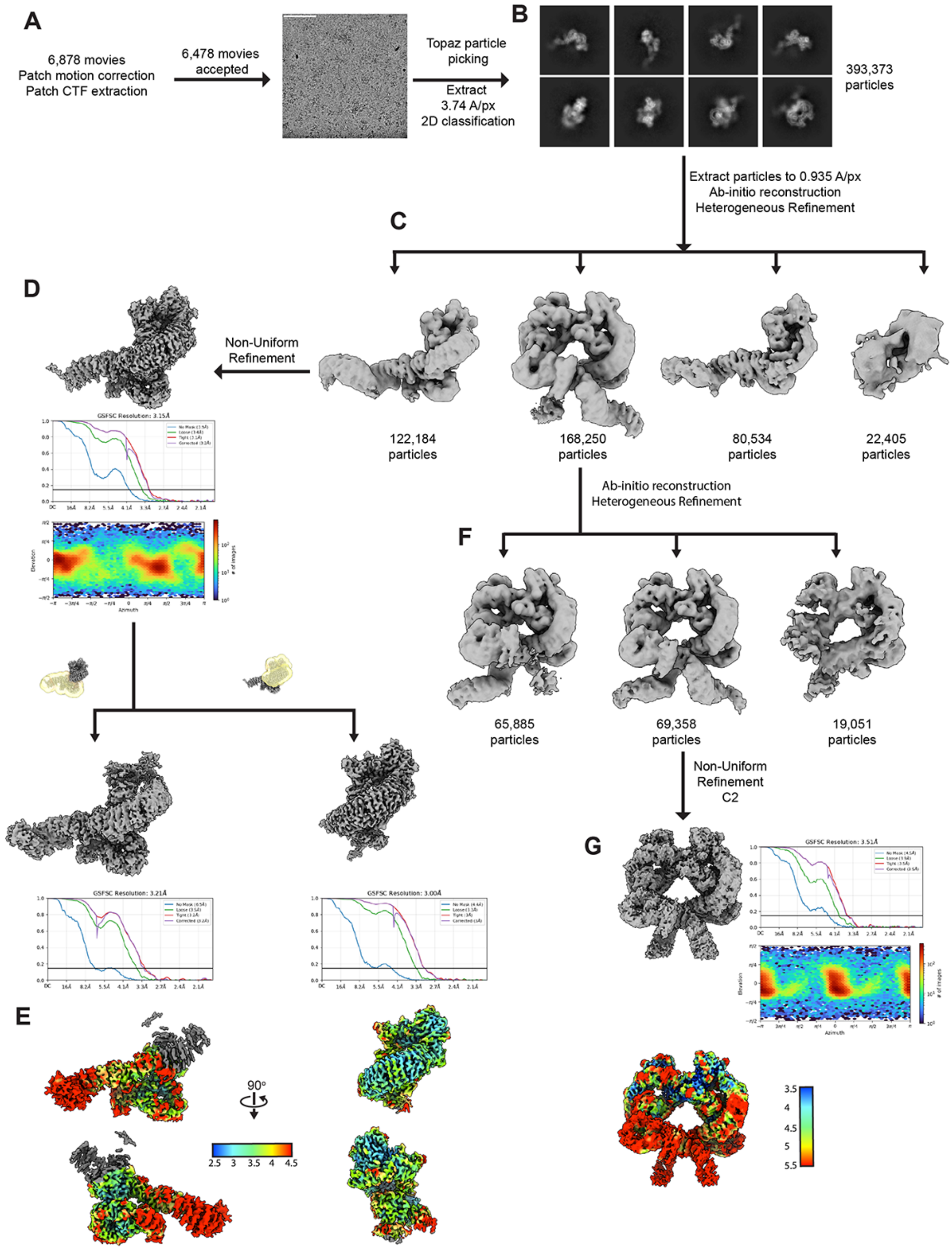

**Figure S14. Cryo-EM data processing pipeline for LRRK2 bound to DARPin G10. (A)** Number of movies collected and kept for further processing, and representative micrograph. **(B)** 2D class averages. **(C)** Data processing strategy. **(D)** Processing of monomeric LRRK2:G10, volume, Euler angle distribution and FSC curve. **(E)** Local resolution map of monomeric LRRK2:G10. **(F)** Processing of dimeric LRRK2:G10. **(G)** Volume, FSC curve, Euler angle distribution and local resolution map of dimeric LRRK2:G10.

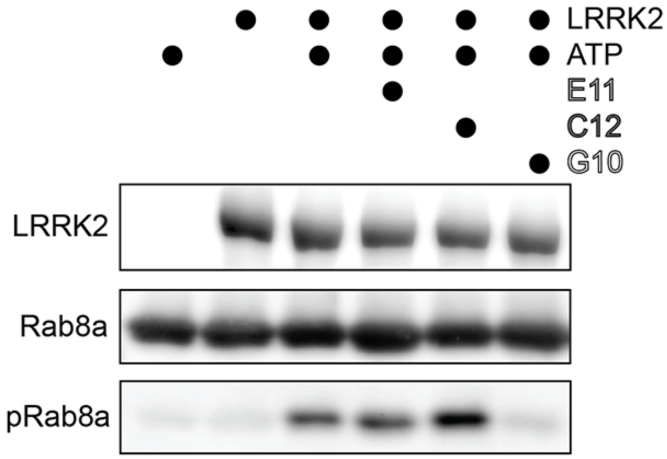

**Figure S15. Representative immunoblot for Figure 5K (in vitro Rab8a phosphorylation assays with DARPins E11, C12, and G10).** LRRK2[WT] was incubated with Rab8a in the presence of ATP and either no DARPin or one of the following DARPins: E11, C12, or G10. ATP was omitted in one condition as a negative control.

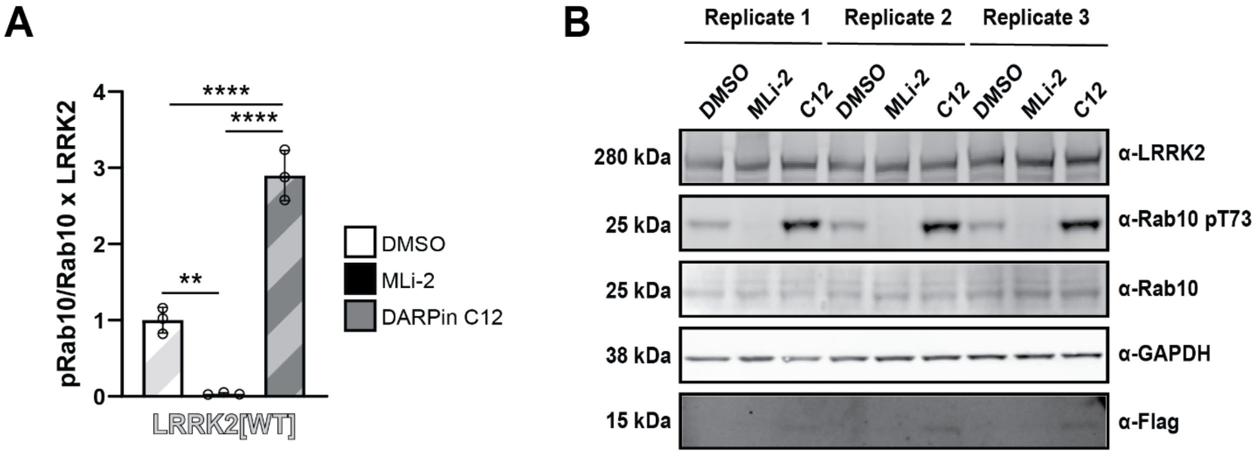

**Figure S16. (A) Quantification and (B) immunoblot of endogenous pRab10 (pT73) / Rab10 x LRRK2 normalized to the mean of LRRK2[WT] + DMSO condition.** HEK 293Ts co-expressing LRRK2[WT] were treated with either DMSO or MLi-2 (5μM) for 1 hour. Cells were lysed and immunoblotted for LRRK2, Rab10, phosphor-Rab10 (T73), GAPDH, and Flag-tagged DARPin C12. All data were analyzed using a two-way ANOVA with Tukey's multiple comparison correction: DMSO vs MLi-2: **p=0.0036; MLi-2 vs DARPin C12: ****p<0.0001; DMSO vs DARPin C12: ****p<0.0001. Error bars ± SD**.**

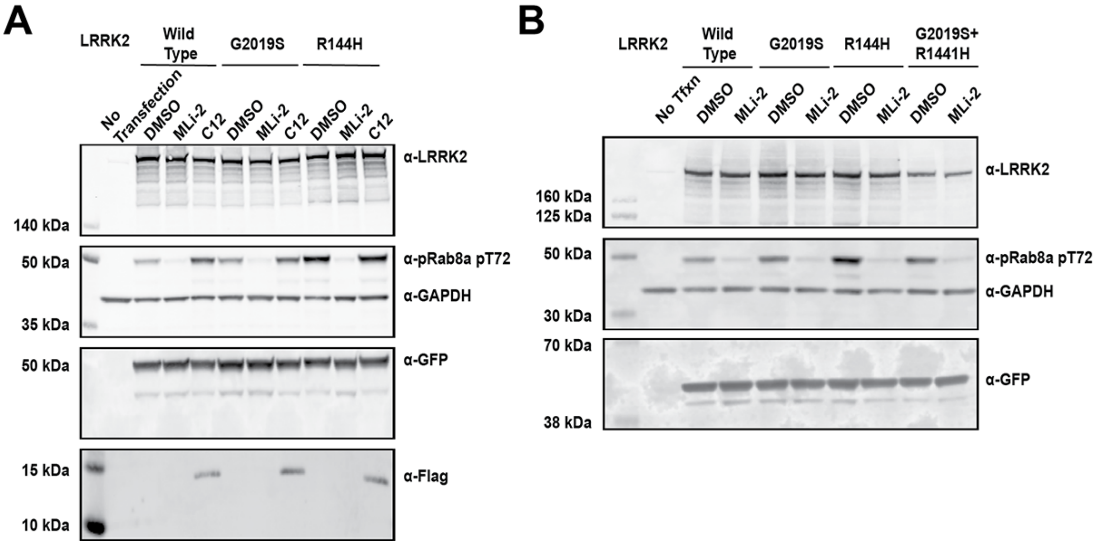

**Figure S17. Representative immunoblots for Figure 6B (effect of DARPin C12 on Rab8a phosphorylation in cells by LRRK2[WT], LRRK2[G2019S], and LRRK2[R1441H]) and Figure 6C (Rab8a phosphorylation in cells by LRRK2[G2019S], LRRK2[R1441H], and LRRK2[G2019S/R1441H]). (A)** Representative immunoblot from 293T cells transiently transfected with GFP 11-tagged LRRK2[WT], LRRK2[R1441H], or LRRK2[G2019S] and its substrate GFP-Rab8a plus or minus Flag-tagged DARPin C12. LRRK2 and its mutants with GFP-Rab8a alone were treated with either DMSO or MLi-2 (5μM) for 1 hour. Lysed cells were immunoblotted for LRRK2, GFP-Rab8a, phospho-Rab8a (pT72), and GAPDH. Samples were immunoblotted for Flag separately. **(B)** Representative immunoblot from 293T cells transiently transfected with GFP 11-tagged LRRK2[WT], LRRK2[R1441H], LRRK2[G2019S], or LRRK2[G2019S/R1441H] and its substrate GFP-Rab8a and treated with either DMSO or MLi-2 (5μM) for 1 hour. Lysed cells were immunoblotted for LRRK2, GFP-Rab8a, phospho-Rab8a (pT72), and GAPDH.

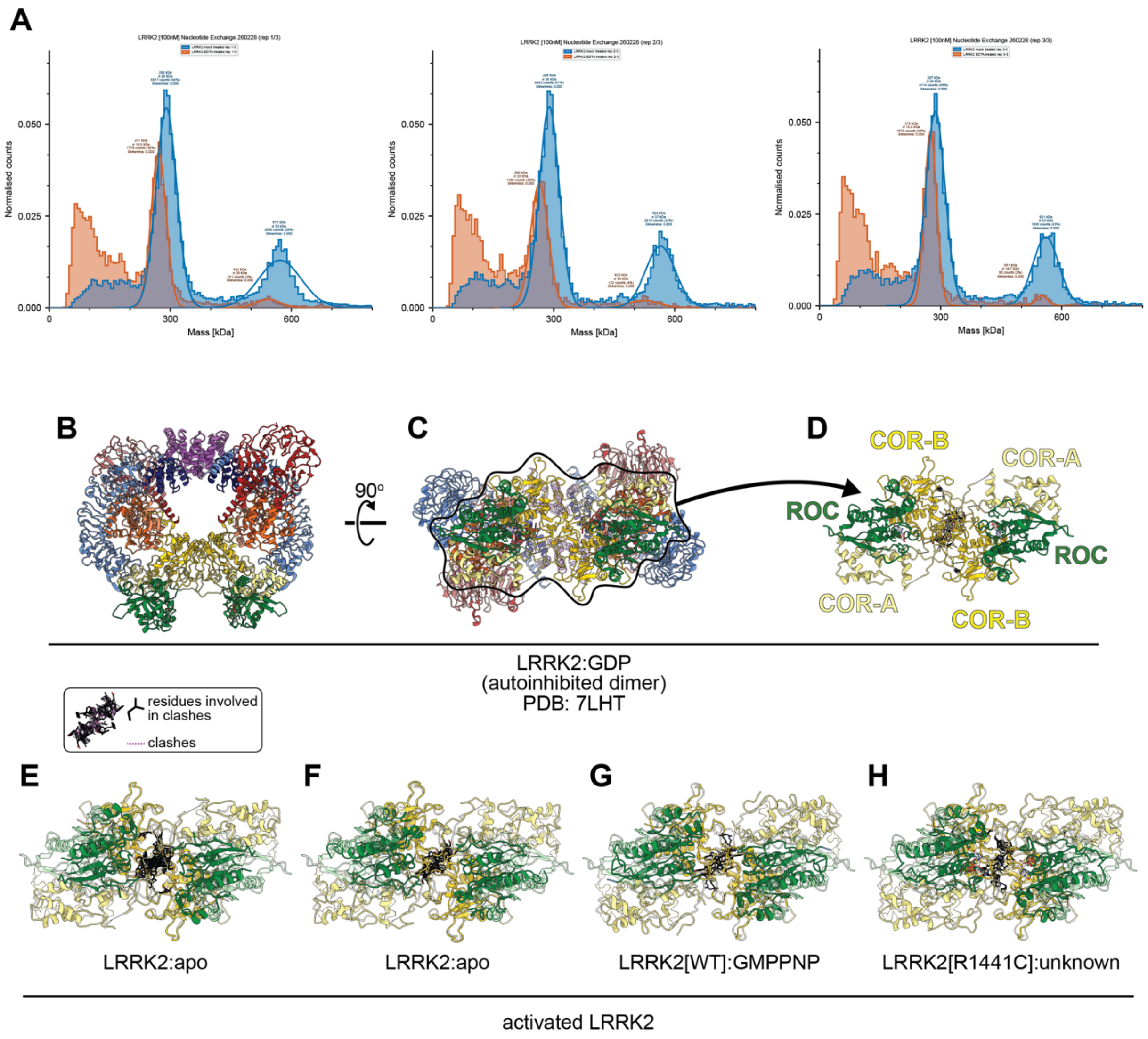

**Figure S18. The active state of LRRK2 is likely to be a monomer. (A)** Treatment of LRRK2 with EDTA shifts the population from the dimeric to the monomeric form. LRRK2 was either treated with EDTA for 2h (as in our nucleotide stripping protocol) or left untreated (mock treatment) (see Methods). The resulting samples were subjected to mass photometry to measure the abundance of LRRK2 monomers and dimers. The three technical replicates of the mass photometry measurements quantified in Figure 4D are shown here. **(B-H)** Modeling of activated LRRK2 in the context of the autoinhibited dimer. **(B,C)** Two views of the dimeric form of autoinhibited LRRK2:GDP (PDB: 7LHT). The view in (C) is from the dimeric interface, which involves the COR-B and COR-A domains. **(D)** Structure of the ROC-COR moieties of the LRRK2 dimer, showing the contacts (dashed black lines) involved in formation of the dimer. **(E-H)** Modeling of dimer interfaces for activated LRRK2. We took the ROC-COR moieties from four structures of activated LRRK2—LRRK2:Apo (1) (E), LRRK2:Apo (2) (F), LRRK2[WT]:GMPPNP (G), and LRRK2[R1441C] (nucleotide undetermined) (H)—and aligned them to the two COR-B domains that form the dimer interface in autoinhibited LRRK2. Residues involved in clashes are shown in black, and the clashes between them, as identified in ChimeraX, as purple dashed lines. Overlayed on these models is a semi- transparent version of panel (D) to highlight the conformational changes between the dimer interface in autoinhibited LRRK2 and the models shown here.

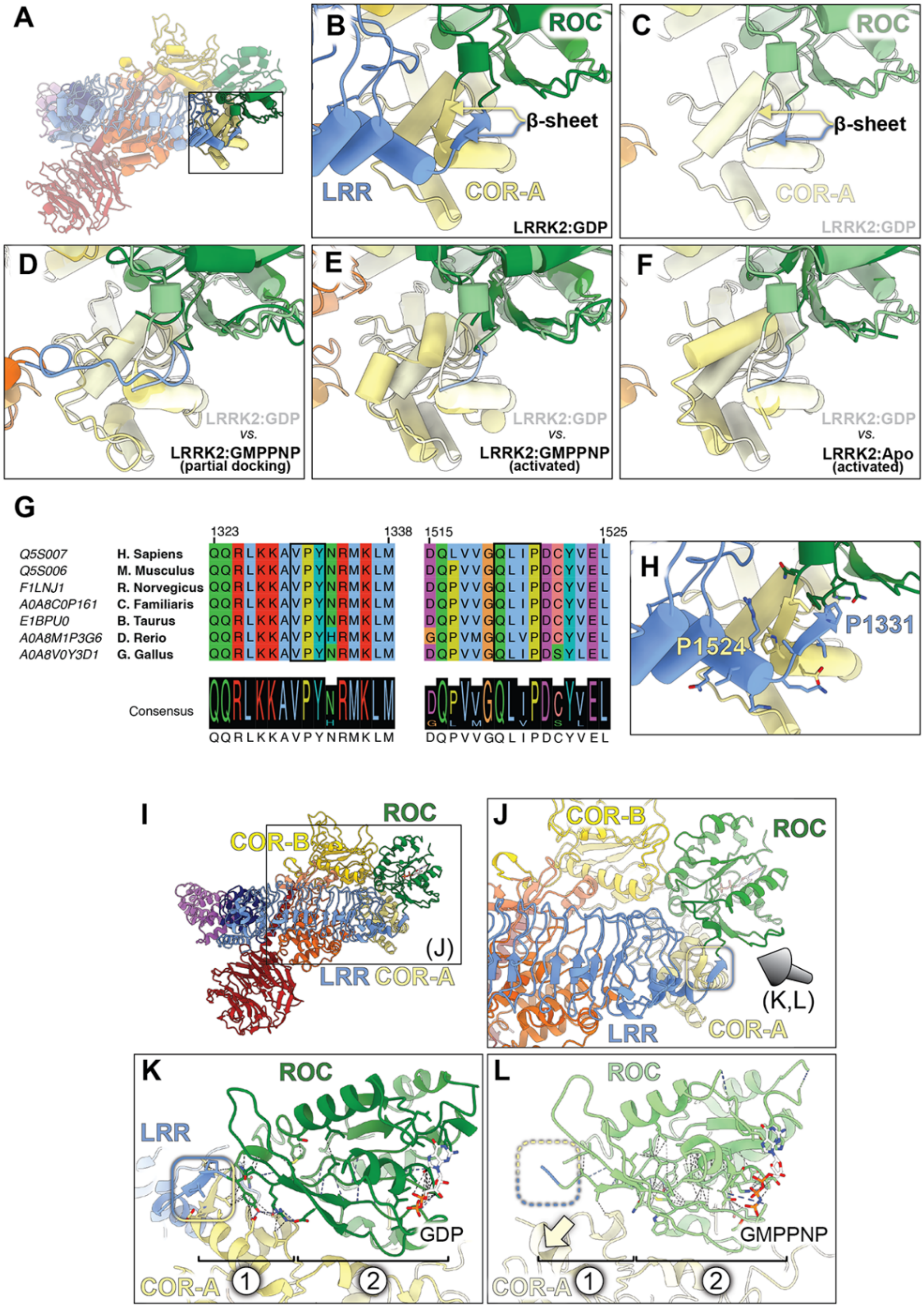

**Figure S19. Model for nucleotide control of the docking and undocking of the N-terminal repeats. (A, B)** In the autoinhibited (GDP-bound) form of LRRK2, the N-terminal repeats of LRRK2 connect to the GTPase (ROC) (A), via one strand of a two-stranded β-sheet (B). The other β-strand connects ROC to the COR-A domain (B). **(C)** Same panel as in (B), but with the N-terminal repeats of LRRK2 removed and the structure shown in lighter shades. This structure will be used as reference in panels (d-f) to illustrate the conformational changes between autoinhibited LRRK2 (in lighter shades) and different structures of activated LRRK2 (in the regular darker shades). **(D)** Comparison between autoinhibited LRRK2 and the structure of GMPPNP-bound LRRK2 (Figure 3C) with partial redocking of the LRR. Formation of the β-sheet is seen in this structure. **(E, F)** Comparison between autoinhibited LRRK2 and GMPPNP-bound (E) or Apo (F) activated LRRK2. The β-sheet is not formed in these structures. **(G)** Sequence alignment of the regions that encompass the two β-strands. **(H)** Close-up of the β-sheet and neighboring region, with the residues included in the alignment shown. The two prolines that mark the ends of the β-strands (P1331 and P1524) are labeled. **(I, J)** Structure of autoinhibited LRRK2, bound to GDP (I), with the region inside the square enlarged in (J). The blue/yellow rounded square in (J) shows the β-sheet connecting the LRR to the C-terminal half of LRRK2. The arrow indicates the viewing direction in panels (K) and (L). **(K, L)** Close-ups of the ROC domain and the region where it connects to the LRR and COR-A domains in the autoinhibited (GDP-bound) (K) and activated (GMPPNP-bound) (L) forms of LRRK2. The blue/yellow rounded square in (K) shows the β-sheet while the dotted version in (L) indicates its absence in the activated form of LRRK2. In addition to the presence or absence of this β-sheet, the two states differ in a set of interactions seen adjacent to the β-sheet when it is formed (region (1) in panel (K)), or close to the nucleotide in the activated (GMPPNP) form of LRRK2 (region (2) in panel (L)). The autoinhibited state of LRRK2 (K) shows several interactions in (1) and few in (2), while the reverse is seen in the activated state (L). The yellow arrow in (L) highlights the movement of the COR-A domain away from ROC responsible for breaking the β-sheet in activated LRRK2.

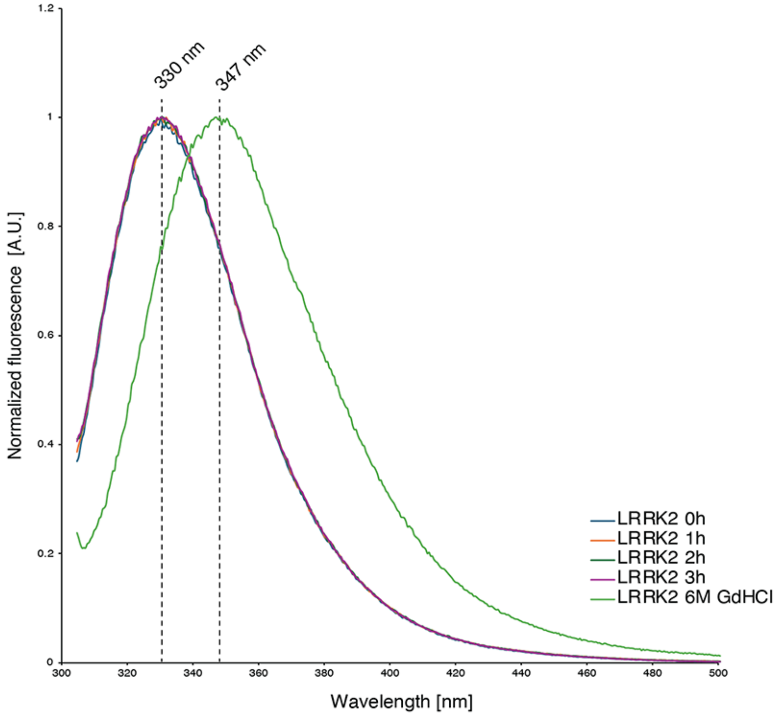

**Figure S20. Fluorescence emission spectra for LRRK2 incubated at room temperature for different times.** To determine the stability of LRRK2 incubated at room temperature for a few hours (needed for the nucleotide-exchange protocol), we measured fluorescence spectra for LRRK2[G2019S] incubated at room temperature at one-hour intervals from 0h (control) to 3h. We also measured the fluorescence spectrum of LRRK2 in 6M guanidinium hydrochloride as an unfolded control. LRRK2 was diluted to 1.56μM (or 40μM tryptophan).

### METHODS

#### Expression and purification of LRRK2

For LRRK2 expression, DNA coding for full-length LRRK2-WT (residues 1-2527) or $LRRK2^{RCKW}$ (residues 1327-2527) (from the Mammalian Gene Collection) was polymerase chain reaction (PCR)-amplified, and the amplicon was inserted into the expression vector pKL: (RRDID:Addgene_110741). The plasmid was used as template for G2019S and R1441C site-directed mutagenesis (Q5 Site-Directed Mutagenesis Kit, NEB). The resulting plasmids were used to generate recombinant baculovirus, according to Bac-to-Bac expression system protocols (Invitrogen). The virus was used to infect SF9 insect cells (RRID:CVCL_0549) with a density of $2x10^6$ cells/ml. Infected cells were cultured in serum-free Insect-XPRESS Medium (Lonza) at 27°C for 3 days post infection. For LRRK2 purification, pelleted Sf9 cells were washed with phosphate-buffered saline (PBS), resuspended in lysis buffer [50mM Hepes (pH 7.4), 500mM NaCl, 20mM imidazole, 0.5mM TCEP, 5% glycerol, 5mM $MgCl_2$, and 20µM GDP] and lysed by homogenization. The supernatant was cleared by centrifugation and loaded onto a Ni-NTA (Qiagen) column. After rinsing with lysis buffer, the $His_6$-Z–tagged protein was eluted in lysis buffer containing 300mM imidazole. The eluate was then diluted to 250mM NaCl with dilution buffer [50mM Hepes (pH 7.4), 0.5mM TCEP, 5% glycerol, 5mM $MgCl_2$, and 20µM GDP] and loaded onto an SP Sepharose column (Cytiva). $His_6$-Z-TEV-$LRRK2^{RCKW}$/$His_6$-Z-TEV-LRRK2 were eluted with a gradient of 250mM to 2.5 M NaCl in dilution buffer and then treated with recombinantly expressed Tobacco Etch Virus (TEV) protease overnight to cleave the $His_6$-Z. Contaminating proteins, the cleaved tag, uncleaved protein, and TEV protease were removed by another combined SP sepharose Ni-NTA (Cytiva) step. Lastly, both LRRK2 and $LRRK2^{RCKW}$ were concentrated and subjected to gel filtration in 20mM Hepes (pH 7.4), 700mM NaCl, 0.5mM TCEP, 5% glycerol, 2.5mM $MgCl_2$, and 20µM GDP, using an ÄKTAxpress system with an S190 16/200 gel filtration column (GE Healthcare). Final yields, as calculated from ultraviolet absorbance, were in the range of 0.9 to 2.2 mg for LRRK2 and $LRRK2^{RCKW}$ respectively per liter of insect cell medium.

#### Expression and purification of DARPins C12 and G10

DARPin E11, C12 and G10 were cloned into a pQE30 vector (Qiagen) containing an N-terminal 8× His tag and a C-terminal 3x-FLAG tag, as previously described[26]. Plasmids were transformed into *E. coli* Rosetta cells. Cells were grown at 37 °C with shaking until the $A_{600}$ reached 1.0. Then, the temperature was reduced to 18 °C and protein expression was induced by adding 0.5mM IPTG. After 18 (H) the cells were harvested by centrifugation and resuspended in lysis buffer (50mM Hepes pH 7.4, 500mM NaCl, 20mM imidazole, 0.5mM TCEP, and 5% glycerol) prior to lysis by sonication. The lysate was cleared by centrifugation and loaded onto preequilibrated Ni-NTA Sepharose beads (GE HealthCare). The beads were washed with 50 column volumes of lysis buffer and protein was eluted in lysis buffer supplemented with 300mM imidazole. The eluate was concentrated to 5 ml and subjected to SEC using an S190 16/200 column (GE HealthCare) in SEC buffer (20mM Hepes pH 7.4, 150mM NaCl, 0.5mM TCEP, and 5% glycerol). Final concentrations for DARPin E11, C12 and G10 were: 6.3mg/mL, 3.6mg/mL and 3mg/mL respectively.

#### Kinase activity assays with DARPins

The impact of DARPin binding on kinase activity was assessed by measuring the phosphorylation of Rab8a. The reaction was set up using 100 nM LRRK2 and incubated for 40 minutes at room temperature with 1.5μM His-Rab8a T22N and with or without 1mM ATP, 10μM DARPin (E11, C12, or G10), or 2.5μM Mli-2 in reaction buffer (50mM HEPES pH 7.4, 150mM NaCl, 0.5mM TCEP, 5mM $MgCl_2$, 5% glycerol, 20μM GTP, 2μM GDP). Reactions were quenched using SDS-PAGE sample buffer, run on an SDS-PAGE gel, and then transferred via Western blot for immunoblot analysis to visualize LRRK2 (1:1000, MJFF2 (c41–2); Abcam ab133474, Lot # GR3445288–3; RRID:AB_2713963), His-Rab8a (1:2000, 6*His Monoclonal antibody; Proteintech 66005, Lot # 10027680; RRID: AB_3086567), and phospho-Rab8a (pT72) (1:500, Abcam 230260, Lot #GR3216587–1;RRID:AB_2814988). All membranes were imaged using the Biorad Chemidoc MP Imaging system. Band intensities were quantified using ImageJ (version 2.14.0). One-way ANOVA with Tukey's multiple comparison correction was performed using Graphpad Prism (version 10.5.0, RRID: SCR_002798) on the ratio of background corrected phospho-Rab in each condition to the average of phospho-Rab in the LRRK2 alone condition (N=3).

#### Kinase activity assay with nucleotide-stripped LRRK2

LRRK2[WT] was incubated with ('EDTA treated) or without EDTA ('mock treated') for 24 hours followed by addition of 20mM $MgCl_2$ and further incubation for 24 hours. Untreated LRRK2 was used immediately upon thawing from frozen aliquots. The effect of nucleotide stripping on kinase activity was assayed by measuring the phosphorylation of Rab8a as described above. The reaction was set up using 100 nM LRRK2 and incubated for 40 minutes at room temperature with 1.5μM His-Rab8a T22N and with or without 1mM ATP, in reaction buffer (50mM HEPES pH 7.4, 150mM NaCl, 0.5mM TCEP, 5mM MgCl2, 5% glycerol). All LRRK2 samples were incubated with Rab8a in the presence of ATP, with one negative control lacking ATP. Background-corrected pRab8a was normalized to the average pRab8a in the untreated LRRK2 + ATP condition from [n=3] independent Western blots. Images were processed in ImageJ.

#### Guanine nucleotide exchange of LRRK2 samples for cryo-EM

We prepared a working stock of 0.1M EDTA (pH 7.5) by diluting 0.2M EDTA pH 7.5 in half with Exchange Buffer (20mM Hepes pH 7.4; 200mM NaCl and 20μM TCEP). Ten μL aliquots of full-length LRRK2 at concentrations of 11–20μM were mixed with 0.5μL of 0.1M EDTA (pH 7.5) (for a final EDTA concentration of 5mM) and incubated for 30 min at 4 °C. After the incubation, a Zeba micro desalting column (Thermo Fisher Scientific) was equilibrated with Exchange Buffer according to the manufacturer's instructions. A total of 10.5 μL of sample was applied to the spin column, followed by 3 μL of Exchange Buffer, and centrifuged for 2 min at 1000 × *g* at 4 °C. After centrifugation, 1.3 μL of 10mM nucleotide (GMP-PNP, Abcam AB146660, or GDP, Sigma-Aldrich G7127) was added followed by 1.3 μL of 20mM $MgCl_2$ to the 13.5 μL sample, for a total volume of 16.1 μL and a final concentration of 0.8mM nucleotide and 1.6mM $MgCl_2$. For mixed-nucleotide samples, 1.3 μL of 750μM GMP-PNP and 250μM GDP were added, followed by 1.3 μL of 20mM $MgCl_2$ to the 13.5 μL sample, for a total volume of 16.1 μL and a final concentration of 0.8mM total nucleotide (200μM GDP and 600μM GMP-PNP) and 1.6mM $MgCl_2$. Samples were incubated for 30–60 min at 4 °C.

Some of our initial cryo-EM analysis revealed that the nucleotide stripping protocol described above was incomplete, and that some LRRK2 remained bound to GDP even in samples that had been stripped and incubated with GMPPNP (Figure 2). To address this issue, we tested a longer EDTA treatment. Ten μL of purified full-length LRRK2 (15μM) was incubated with 5mM EDTA (pH 7.5) for 24 hr at 4 °C. After incubation, a Zeba micro desalting column was equilibrated with Exchange Buffer as described above. A total of 10.5 μL of sample was applied to the spin column, followed by 3 μL of Exchange Buffer, and centrifuged for 2 min at 1000 × *g* at 4 °C. After centrifugation, 1.3 µL of 10mM GMP-PNP was added, followed by 1.3 µL of 20mM $MgCl_2$, and the sample was incubated for 24 hr at 4 °C, for a final concentration of 0.8mM nucleotide and 1.6mM $MgCl_2$ .

**Guanine nucleotide exchange of LRRK2 samples for kinase activity and fluorescence measurements**

Full-length human LRRK2 at 20µM was thawed on ice and diluted to 500nM in a nucleotide stripping buffer (50mM HEPES, 150mM NaCl, 1mM TCEP, 5mM EDTA, and 1mM GMPPNP or GDP). Because of the incomplete nucleotide exchange we detected in our cryo-EM samples that had been stripped of their nucleotide for 30-60min at 4 °C, each sample was incubated at room temperature for 3 hours. We verified that LRRK2 was stable at room temperature for this long by measuring fluorescence emission spectra of LRRK2 incubated at room temperature for up to 3 hours (Figure S20). After the stripping, 10mM $MgCl_2$ was added to neutralize EDTA, then samples were incubated for another 30 minutes at room temperature and nucleotide (GMPPNP or GDP) was added to a final 100µM. Each sample (LRRK2 with GMPPNP or LRRK2 with GDP) was transferred into mini dialysis devices (Thermo Scientific, 69572) and dialyzed against 300mL of nucleotide exchange buffer (50mM HEPES, 150mM NaCl, 5mM $MgCl_2$, 1mM TCEP, and 0.1µM GMPPNP or GDP) overnight in the cold room with gentle stirring. The next day samples were spun down on a tabletop centrifuge at maximum speed and directly used for the measurements.

**Guanidine nucleotide exchange for mass photometry**

12.7µM of LRRK2[WT] in 5 µL of buffer (20mM Hepes pH 7.4, 150mM NaCl, 5% glycerol, 0.5mM TCEP, 20uM GDP and 2.5mM MgCl2 ) was mixed with one of the following: (1) 0.28 µL of 100mM EDTA pH 7.5 and 0.28uL of 100µM GDP ('EDTA treated'), or (2) 0.28 µL of milliQ H2O and 0.28 µL of 100µM GDP ('mock treated'). Samples were incubated on ice for 2 hours, at which point 0.6 µL of 20mM $MgCl_2$ were added and samples were incubated on ice for another 24 hours. Samples were then diluted using with 20mM Hepes pH 7.4, 150mM NaCl, 5% glycerol, 0.5mM TCEP, and 1mM $MgCl_2$, to a final concentration of LRRK2[WT] of 100nM and incubated on ice for 30 minutes before mass photometry measurements.

Total particle counts corresponding to the molecular weights of either monomer or dimer were determined from mass photometry traces. The percentage of dimers contributing to total monomer/dimer LRRK2 landing events were analyzed with one-way analysis of variance (ANOVA) and corrected for multiple comparisons using Tukey's test.

**Sample preparation for Cryo-EM: nucleotide-exchanged LRRK2**

LRRK2, following nucleotide stripping and exchange, was incubated with MLi-2 (10mM; Tocris, RRID: Addgene_226784), and E11-DARPin, purified as previously described[26]. The incubation was performed at a molar ratio of 1:1.5:1.25 (LRRK2:MLi-2:E11) for 10 min at room temperature, followed by 15 min at 4 °C. The resulting complex was diluted to a final concentration of 7µM LRRK2 in Exchange Buffer (20mM HEPES, pH 7.4, 200mM NaCl, 20µM TCEP) and supplemented with 21µM DDM (GOLDBIO: DDM5_69227). A volume of 2.8–3.5 µL was applied to glow-discharged UltraAUFoil Holey Gold 300 mesh R1.2/1.3 grids (Quantifoil, Q350AR13A) and incubated in a FEI Vitrobot IV chamber at 4 °C and 95 % humidity for 5 s. Excess liquid was blotted for 2 s using filter paper 595 at blot force 4, and the grids were vitrified by plunging into liquid ethane cooled to liquid-nitrogen temperature.

One data set for LRRK2$^{R1441C}$ was acquired on a Talos Arctica (FEI) operated at 200 keV and equipped with a Falcon 4i detector (Thermo Fisher Scientific). Images were recorded at a nominal magnification of 150,000× in EF-TEM mode (0.95 Å/pixel) with a cumulative electron exposure of approximately 55 e⁻/Å². The remaining data sets for LRRK2$^{WT}$, LRRK2$^{G2019S}$, and LRRK2$^{R1441C}$ were acquired on a Titan Krios G3 (Thermo Fisher Scientific) operated at 300 keV, equipped with a Falcon 4 direct electron detector and a Gatan BioContinuum energy filter. Images were collected at a nominal magnification of 130,000× in EF-TEM mode (0.935 Å/pixel) using a 20 eV energy filter slit width, with a cumulative electron exposure of approximately 55 e⁻/Å².

#### Sample preparation for Cryo-EM: DARPin-bound LRRK2 and LRRK2$^{RCKW}$

For LRRK2 bound to G10-DARPin, a 1:5 molar ratio (LRRK2:G10) was incubated for 10 min at room temperature, followed by 15 min at 4 °C. The complex was diluted to a final concentration of 5μM LRRK2 in buffer containing 20mM HEPES (pH 7.4), 200mM NaCl, 20μM TCEP, 20μM GDP, 2.5mM $MgCl_2$, and 5 % glycerol, and supplemented with 21μM DDM (GOLDBIO: DDM5_69227). A 2.8 μL volume was applied to glow-discharged UltraAUFoil Holey Gold 300 mesh R1.2/1.3 grids (Quantifoil, Q350AR13A) and incubated in a FEI Vitrobot IV chamber at 4 °C and 95 % humidity for 5 s. Excess liquid was blotted for 2 s using filter paper 595 at blot force 4, and the grids were vitrified by plunging into liquid ethane cooled to liquid-nitrogen temperature. Data was collected under the same conditions described above on a Titan Krios G3.

As previously described[26] LRRK2$^{RCKW}$[R1441H]:C12-DARPin was prepared by desalting LRRK2 into a buffer containing 20mM HEPES (pH 7.4), 150mM NaCl, 20μM TCEP, 20μM GDP, 2.5mM $MgCl_2$, and 5 % glycerol using the Zeba micro desalting column (Thermo Fisher). LRRK2$^{RCKW}$ was then incubated with C12-DARPin in a ratio of 1:1.25 (LRRK2:C12) for 10 min at room temperature, followed by 15 min at 4 °C. The sample was diluted to a final LRRK2$^{RCKW}$ concentration of 6μM. A 4 μL volume was applied to glow-discharged UltrAUFoil Holey Gold 300 mesh R1.2/1.3 grids (Quantifoil, Q350AR13A) and incubated in a FEI Vitrobot IV chamber at 4 °C and 95 % humidity for 20 s. Excess liquid was blotted for 4 s using filter paper 595 at blot force 4, and the grids were vitrified by plunging into liquid ethane cooled to liquid-nitrogen temperature. Images were collected at a nominal magnification of 36000x in EF-TEM mode (1.16 Å/pixel), with a cumulative electron exposure of approximately 51 e⁻/Å². Data was collected on a FEI Talos Arctica operated at 200 kV and equipped with a K2 summit direct electron detector (Gatan).

#### Cryo-EM data data processing and model building

Movies for all datasets were pre-processed using Patch MotionCor and Patch CTF estimation in CryoSPARC[47] (RRID:SCR_016501) to perform motion correction and estimate the contrast transfer function (CTF). Micrographs with a CTF fit worse than 5.5 Å were excluded from further processing. Particles were picked for all datasets using a Topaz[48] model (version 0.2.4, https://cb.csail.mit.edu/topaz/) previously trained on the dataset. For combined datasets, several additional rounds of Topaz training and two-dimensional (2D) classification were performed. Particle stacks were extracted at the original pixel size. Ab initio and heterogeneous refinement jobs were then used to discard poor-quality particles, followed by non-uniform refinement and local refinement (see Figures S1-S6, S9, S13, and S14). To build the initial LRRK2 model, the highest-resolution local maps obtained for the LRRK2:G10 complex were used (EMD-70981). The available structure of full-length LRRK2[9] (PDB 7LHW) was used as the starting point, yielding the final model PDB 9OXH. This model was subsequently split into domains, docked into the corresponding cryo-EM maps using UCSF ChimeraX[49] (version 1.9, RRID:SCR_015872), and merged into one chain. For models containing density for MLi-2, we built the model for the part of the map corresponding to the kinase and WD40 domains starting from our previous structure of MLi-2-bound LRRK2$^{RCKW}$ (PDB 8TXZ)[16]. For DARPin C12 and G10, initial models were generated using AlphaFold 3[50]. Models were fit into the density maps using Coot[51] (version 0.9.8, www2.mrc-lmb.cam.ac.uk/personal/pemsley/coot/, RRID:SCR_014222) and refined in Phenix to generate the final models. Maps for LRRK2$^{WT}$[GMPPNP] and LRRK2$^{R1441C}$ were improved with Resolve in Phenix[52]. Data collection and refinement statistics are summarized in Table S1.

Movies for LRRK2$^{RCKW}$:C12 DARPin complex, were collected and pre-processed using Patch MotionCor and Patch CTF estimation in CryoSPARC[47] (RRID:SCR_016501). Micrographs with CTF fit worse than 4 Å were excluded from further processing. Particles obtained with blob picker in CryoSPARC were used to train a Topaz[48] model. After several rounds of 2D Classification, particles were extracted with a 288-pixel box and used for subsequent processing. Two final local refinements were performed using focused masks. The first refinement targeted the C-lobe of the kinase, the WD40 domain, and the C12 DARPin, while the second focused on the ROC domain, COR-A, COR-B, and the N-lobe of the kinase. These refinements yielded maps at 3.48 Å and 3.6 Å resolution, respectively. Local refinement procedures were conducted to evaluate map quality and directional resolution anisotropy. The available structure of LRRK2$^{RCKW}$ (PDB:6VP7) was fitted into the 3D maps using UCSF ChimeraX[49] and used as a starting point for model building, which was performed in Coot[51]. The built structures were refined using real-space refinement as implemented in Phenix[52] Data collection and refinement statistics are summarized in Table S1.

**Quantitation of the distribution of autoinhibited and activated LRRK2 in samples treated with EDTA for 1h vs. 24h**

This analysis included particle counts from samples subjected to the nucleotide stripping protocol for 1h and then incubated with GDP and/or GMPPNP yielding active volumes (see Figures S4C–4D for monomers and Figure S2F for mixed dimers) or autoinhibited volumes (see Figures S2C–2D and Figure S2F for mixed dimers), excluding data for the mutants (LRRK2[G2019S] and LRRK2[R1441C]). The counts were then further separated according to the nucleotide added after stripping. We also used particle counts from LRRK2 subjected to 1h nucleotide stripping without subsequent addition of nucleotide (see Figure S8) and data from LRRK2 subjected to 24h nucleotide stripping followed by addition of GMPPNP (see Figures S5D–5F). The final relative abundance of particles corresponding to the active and autoinhibited states of LRRK2[WT] in the different samples is presented in Figure 2H and Figure S8D.

**Cryo-EM data collection for untreated (no nucleotide stripping) LRRK2[WT] and LRRK2[R1441C]**

Cryo-EM data were collected on a Talos Arctica transmission electron microscope operated at 200 kV equipped with a Gatan K3 direct electron detector. Images were acquired at a nominal magnification of 36,000×, corresponding to a calibrated super-resolution pixel size of 0.5505 Å/pixel (1.101 Å/pixel in counting mode). Movies were collected using Leginon v3.7 at a dose rate of 21.22 $e^{-}/Å^2/s$ over 2.40 s (total dose 50.92 $e^{-}/Å^2$), fractionated into 40 frames. A total of 2,371 movies were collected over a defocus range of 1.7–2.6 µm. Ice thickness was estimated as described previously.

On-the-fly processing was performed using MotionCor2 for motion correction and CTFFIND4 for CTF estimation within the Appion pipeline. Motion-corrected, dose-weighted micrographs (1.101 Å/pixel) were imported into cryoSPARC. Initial particle picking was performed using Warp, and picks from each dataset (WT and R1441C) were used to generate templates for cryoSPARC-based template picking.

**Cryo-EM data processing for untreated (no nucleotide stripping) LRRK2[WT] and LRRK2[R1441C]**

Motion-corrected micrographs from WT and R1441C datasets were combined prior to downstream processing to ensure consistent classification and minimize bias. After curation based on CTF quality and ice thickness (2,257 micrographs), initial particle picking was performed in cryoSPARC using template-based methods, yielding 1,420,224 particles. Representative particles were selected and used for Topaz-based particle picking, resulting in a refined dataset of 424,769 particles. This was subjected to two rounds of 2D classification to remove junk particles, yielding a final cleaned subset of 92,497 particles for downstream analysis.

These particles were used for ab initio reconstruction and heterogeneous refinement to separate junk, flexible, and autoinhibited conformations. Particles corresponding to autoinhibited states were selected, re-centered, and re-extracted at 3.03 Å/pixel (32,759 particles). These particles were further classified by ab initio reconstruction and heterogeneous refinement to resolve autoinhibited monomeric and dimeric states. The autoinhibited dimer subset was subjected to 3D classification in cryoSPARC into two classes, yielding a mixed monomer/dimer class (11,468 particles) and a dimer class (9,033 particles).

Final particle distributions were determined by tracing particles back to their dataset of origin following joint classification. Particles were grouped into four structural categories: Activated LRRK2 (“RCKW”), Autoinhibited Monomer, Autoinhibited Dimer, and Mixed Dimer. The mixed monomer/dimer class was conservatively treated as contributing equally to active and inactive states for downstream quantification.

**GTP turnover assay**

GTP hydrolysis was monitored using a real-time inorganic phosphate detection assay (EnzCheck Phosphate Assay Kit; Thermo Fisher Scientific/Invitrogen, E6645). This assay relies on the enzymatic conversion of the chromogenic substrate 2-amino-6-mercapto-7-methylpurine ribonucleoside (MESG) by purine nucleoside phosphorylase (PNP) in the presence of inorganic phosphate. Phosphate released during GTP hydrolysis drives the MESG/PNP reaction, resulting in a spectral shift in absorbance from 330 nm to 360 nm, which was continuously recorded using a spectrophotometer (Agilent G6860AA). Reactions were carried out in 60 µL volumes in quartz cuvettes and contained 1x reaction buffer (50mM Tris-HCl, 150mM KCl, 1mM $MgCl_2$, pH 7.5), MESG substrate (0.2mM final), and PNP (1 U/mL final). Absorbance at 360 nm was monitored in real time for 3 min at 22°C. Phosphate release was quantified from the change in absorbance using the extinction coefficient of 2-amino-6-mercapto-7-methylpurine ($\varepsilon$ = 11,000 $M^{-1}cm^{-1}$). To minimize phosphate contamination, all solutions were prepared with $P_i$-free water, and cuvettes were thoroughly rinsed with deionized water between measurements. For DARPin assays, purified LRRK2 protein (1µM final) was incubated with or without individual DARPins (G10 or C12; 1.2µM final) for 30 min at room temperature prior to measurement. For Rab8a Q67L–dependent assays, LRRK2 (1µM) was pre-incubated with G10 or C12 (1.2µM) for 30 min, followed by the addition of Rab8a Q67L (20µM). After an additional 10 min incubation, GTP hydrolysis rates were measured as described above.

#### In vitro guanine nucleotide exchange assay

Nucleotide exchange on LRRK2 was measured using a fluorescence-based assay as described previously[53]. Recombinant LRRK2 protein (1μM) was preloaded with 1mM BODIPY-FL–GDP (G22360, Thermo Fisher Scientific) in exchange buffer containing 20mM HEPES (pH 7.2), 150mM KCl, 5% glycerol, and 1mM DTT. The loading reaction was performed at 4 °C for 5 h. Nucleotide loading was terminated by the addition of $MgCl_2$ to a final concentration of 10mM.
To initiate nucleotide exchange, BODIPY-FL–GDP–loaded LRRK2 was rapidly mixed with C12 (1.2μM) and/or Rab8A Q67L (20μM) in exchange buffer supplemented with 5mM GTP. Reactions were carried out in black 384-well plates. Fluorescence signals were monitored in real time using a microplate reader (Cytation 5, Agilent BioTek) with excitation at 488 nm and emission at 535 nm. Measurements were recorded every 10 s for 6 min.

#### Analysis of phosphorylated Rab8a in cells

For in-cell LRRK2 activity assays, $1.2 \times 10^6$ 293T cells were plated into each well in six well plates and incubated at 37 °C in 5% CO2. After two days, cells were transfected with 500ng GFP-Rab8a (EGFP C1 vector backbone; Addgene; RRID: Addgene_49543) plus or minus 500ng 8xHis-DARPin C12-3xFlag (pcDNA3.1 vector backbone) and 1000ng of full-length LRRK2 wild type, G2019S, R1441H, or the double mutant G2019S+R1441H (all LRRK2 in the pCDNA5 vector backbone) using polyethylenimine (Polysciences, catalog # 23966-1). After 48 hours, control cells not transfected with DARPin C12 were treated with either DMSO or MLi-2 (Cayman Chemical Company, catalog #19305; 5μM) and incubated for an hour. Cells were washed three times in cold DPBS then lysed in 300uL cold RIPA buffer per well (50mM Tris pH8, 150mM NaCl, 1% Triton X-100, 0.1% SDS, 0.5% Sodium Deoxycholate, 1mM DTT in the presence of cOmplete mini EASYpack protease inhibitor cocktail and the PhosSTOP EASYpack phosphatase inhibitor cocktail (Sigma-Aldrich, catalog # 05056489001; Roche, 04906837001)). Cell lysates were vortexed twice for 30 seconds then rotated for 15 minutes at 4ºC. Lysates were clarified by centrifugation at 17,000g for 15 minutes at 4ºC, and the supernatant was moved to a new tube. SDS-PAGE loading buffer and reducing agent (1X, Invitrogen, catalog # NP0007 and NP0009) were added to the supernatant which was then boiled at 95ºC for 10 minutes. Samples were spun again at 13,000g for five minutes, and supernatant was moved to a new tube. Samples were either used immediately or stored at −20°C. For immunoblotting, samples were processed as described previosly[26]. In short, the membrane was cut at the 70kDa molecular weight marker and the top portion was immunoblotted for LRRK2 with a specific antibody (1:1000, MJFF2 (c41–2); Abcam catalog # ab133474, Lot # GR3445288–3; RRID:AB_2713963) in 1% BSA. The lower portion was immunoblotted using antibodies for phospho-Rab8a (pT72) (1:1000, Abcam catalog # 230260, Lot # GR3216587–1;RRID:AB_2814988), GFP (1:2500, Santa Cruz Biotechnology catalog # sc-9996, Lot # C2523; RRID:AB_627695), and GAPDH (1:3000, Cell Signaling Technology catalog # 2118, Lot # 16; RRID:AB_561053) in 1% milk. To visualize DARPin C12, cell lysate was reduced in Tricine SDS sample buffer (Invitrogen, catalog # LC167). Samples were run on Tricine 10%-20% gels (Invitrogen, catalog # EC66252BOX) and run in tricine running buffer (Invitrogen, catalog # LC1675) for 1 hour and 20 mins at 125V. Protein was then transferred onto polyvinylidene fluoride (PVDF) transfer membrane (0.2 μm pore size) for 75 min at 300 mA at 4 °C. Samples were then blocked in 5% milk for an hour and incubated for 16 hours at 4 ºC in primary rabbit anti-FLAG polyclonal antibody (ptg labs, catalog # 20543-1-AP, Lot # 00106090; RRID:AB_11232216) in 1% milk. Samples were washed three times in Tris buffered saline buffer with 0.05% Tween-20 (TBS-T) for 10 minutes then incubated with goat anti-rabbit Licor680 antibody (LI-COR biosciences; catalog # 926-32211, Lot # D40109-05; RRID:AB_621843 ) in 4% milk for 1 hour at room temperature. All membranes were imaged using the Li-Cor imaging system. Intensities of phospho-Rab8a, GFP, and LRRK2 bands were measured using ImageJ (version 2.14.0). Two-way ANOVA were conducted on the ratios of phosphorylated Rab8a to total Rab8a and LRRK2, normalized to the mean of the wild type LRRK2 plus DMSO group using GraphPad Prism (version 10.5.0, RRID: [43]) with Tukey's multiple comparison correction.

#### RMSD analysis

The RMSD values reported in Figure S9 were obtained by doing pairwise comparisons between the ROC-COR modules of the structures shown in the figure. Specifically, we extracted the part of each model corresponding to residues 1332-1878 (ROC, COR-A, and COR-B). For each pairwise comparison, models were aligned to each other in ChimeraX[49] ("match"), using only their backbones. Once aligned, the RMSD for the entire ROC-COR module was calculated in ChimeraX[49] ("rmsd"), also using only the backbone atoms.

#### Figure preparation

All structural representations were prepared using ChimeraX[49].

### QUANTIFICATION AND STATISTICAL ANALYSIS

Statistical tests are noted in figure legends. All data are shown as means ± s.d., and all analyses were performed using Prism 10 software (Graphpad).
